# The erosion of biodiversity and biomass in the Atlantic Forest biodiversity hotspot

**DOI:** 10.1101/2020.09.08.287938

**Authors:** Renato A. F. Lima, Alexandre A. Oliveira, Gregory R. Pitta, André L. de Gasper, Alexander C. Vibrans, Jérôme Chave, Hans ter Steege, Paulo I. Prado

## Abstract

Tropical forests are being deforested worldwide, and the remaining fragments are suffering from biomass and biodiversity erosion. Quantifying this erosion is challenging because ground data on tropical biodiversity and biomass are often sparse. Here, we use an unprecedented dataset of 1,819 field surveys covering the entire Atlantic Forest biodiversity hotspot. We show that 83–85% of the surveys presented losses in forest biomass and tree species richness, functional traits and conservation value. On average, forest fragments had 25–32% less biomass, 23–31% fewer species, and 33%, 36% and 42% fewer individuals of late-successional, large-seeded and endemic species, respectively. Biodiversity and biomass erosion were both lower inside strictly protected conservation units, particularly in large ones. We estimate that biomass erosion across the Atlantic Forest remnants was equivalent to the loss of 55-70 thousand km^2^ of forests or US$2.3-2.6 billion in carbon credits. These figures have direct implications on mechanisms of climate change mitigation.

## Introduction

Tropical forests are major stocks of biodiversity and carbon; and these stocks are declining worldwide. Half of their original cover has already vanished and current deforestation rates are about 1% per year^1^. Human impacts on tropical forests, however, are not restricted to deforestation. Beyond the reduction in habitat availability and connectivity, deforestation triggers a myriad of modifications that can penetrate up to 1.5 km into the remaining fragments^2–4^. In addition, forest fragments are more accessible, increasing their exposure to fire, selective logging, hunting, and biological invasions. These human-induced impacts on forest fragments (*i.e*. forest degradation) impose a long-lasting burden on forest biodiversity and biomass stocks^5–12^ that can be as severe as deforestation^13,14^. Protected areas can mitigate the erosion of biodiversity and biomass^15–17^, but their effectiveness is contingent on the type of management and level of anthropogenic pressure surrounding the protected areas^15–17^.

Forest degradation can be assessed by high-resolution remote sensing (*e.g*. LiDAR^18^), but the coverage of this approach is limited and the impact on biodiversity cannot be measured. This is why large-scale quantifications of the impacts of forest degradation are mostly available for biomass^10–12^. Therefore, field surveys remain essential to quantify the erosion of both biodiversity and biomass^2–4,7,8,10,19^. The simultaneous evaluation of forest degradation on tropical biodiversity and biomass at large-scales provides crucial knowledge for the conservation and climate change agenda^13,17,20^ and to refine regional assessments of biodiversity and ecosystem service (*e.g*. Intergovernmental Science-Policy Platform on Biodiversity and Ecosystem Services - IPBES).

Here, we aim at quantifying the impacts of forest degradation on a major biodiversity hotspot located in eastern South America, the Atlantic Forest (Supplementary Fig. 1). Home to 35% of the South American population, the Atlantic Forest is one of the most fragmented tropical/subtropical forests in the world^12,21^, which may well represent the present or future of other tropical forests worldwide^22^. To achieve our goal, we create one of the largest data sets of forest surveys ever assembled for the tropics and subtropics^23^, both inside and outside protected areas. This data set includes data on forest biomass and tree species richness and/or composition, as well as carefully curated metadata associated with each survey, representing a total of 1,819 field surveys, 1.45 million trees, 3,124 tree species, and 1,238 ha of sampling coverage (Supplementary Table 1, Supplementary Data 1). The data set covers the entire range of environmental conditions, landscape contexts, and disturbance histories of the Atlantic Forest (Supplementary Fig. 2, Supplementary Table 2). It also contains information on multiple species properties, including plant functional traits (i.e., wood density, maximum height, seed mass), ecological groups (or successional status, *e.g*. pioneer) and their conservation value (i.e., threat status and endemism level), which enable to assess human-induced impacts on community composition *sensu lato*.

Using this unprecedented data set, we quantified forest degradation impacts on the aboveground biomass stocks, tree species richness, and multiple species properties. More specifically, we assess the extent and magnitude of those impacts by asking: (i) how pervasive negative impacts are across this biodiversity hotspot? (ii) how much these biodiversity and biomass losses represent compared to low-disturbance Atlantic Forests? And (iii) can protected areas mitigate those losses? Next, we explore the implications of our results to the conservation of what’s left of this biodiversity hotspot by (iv) projecting forest degradation impacts to the remaining Atlantic Forest area to estimate the total amount of carbon lost. We also (v) explore the costs and benefit of two contrasting restoration scenarios: one that focused only on reducing the within-fragment disturbance level (*i.e*. ‘fragment restoration’ scenario) and another focused on increasing fragment size and landscape connectivity (*i.e*. ‘landscape restoration’ scenario).

In general lines, we quantified forest degradation impacts on Atlantic Forest biodiversity and biomass as follows. First, the variation in forest biomass, species richness and species properties were described using linear mixed-effects regression models. These models accounted for the effects of environmental and human-related variables, as well as sampling and biogeographical effects (see Methods and Figs. 3-7), which explained 53% of the variation in biomass, 71% in species richness and 26-44% in species properties (Supplementary Fig. 8, Supplementary Table 3). Next, we used these models to generate baseline predictions in the absence of major human impacts, *i.e*., predictions as if all sites were large, low-disturbance forest patches in landscapes with 100% of forest cover. We validated the precision of these predictions using simulations (Supplementary Table 4). Finally, we defined an index of loss due to human-induced impacts, defined as the standardized difference between observed values and baseline predictions (Supplementary Fig. 9). Values of the index close to zero indicate little human impact and the more negative the value, the greater the impact.

## Results and discussion

### Extent and magnitude of human impacts

We found that the majority of the Atlantic Forest surveys (83%) presented losses of species richness and biomass (Fig. 1), *i.e*. negative indices of loss. These losses were correlated with each other (Fig. 1), meaning that fragments that suffer greater losses of biomass also lose more species. In absolute terms, human-induced impacts corresponded to average declines of 23-32% of the richness and biomass relative to low-disturbance Atlantic Forests (Table 1, Supplementary Table 5). Similar estimates (18-57%) have been reported at smaller spatial scales for Neotropical rainforests^8,10,13^ and at global scale^11,14,17^, suggesting that our estimates are representative of the Atlantic Forest. We did not explicitly model the distances to forest edge (see Supplementary Methods), where biomass erosion can be even greater^2,6–8^. But assuming that researchers tend to avoid edges while establishing plots, our estimates are probably conservative.

**Figure 1.**
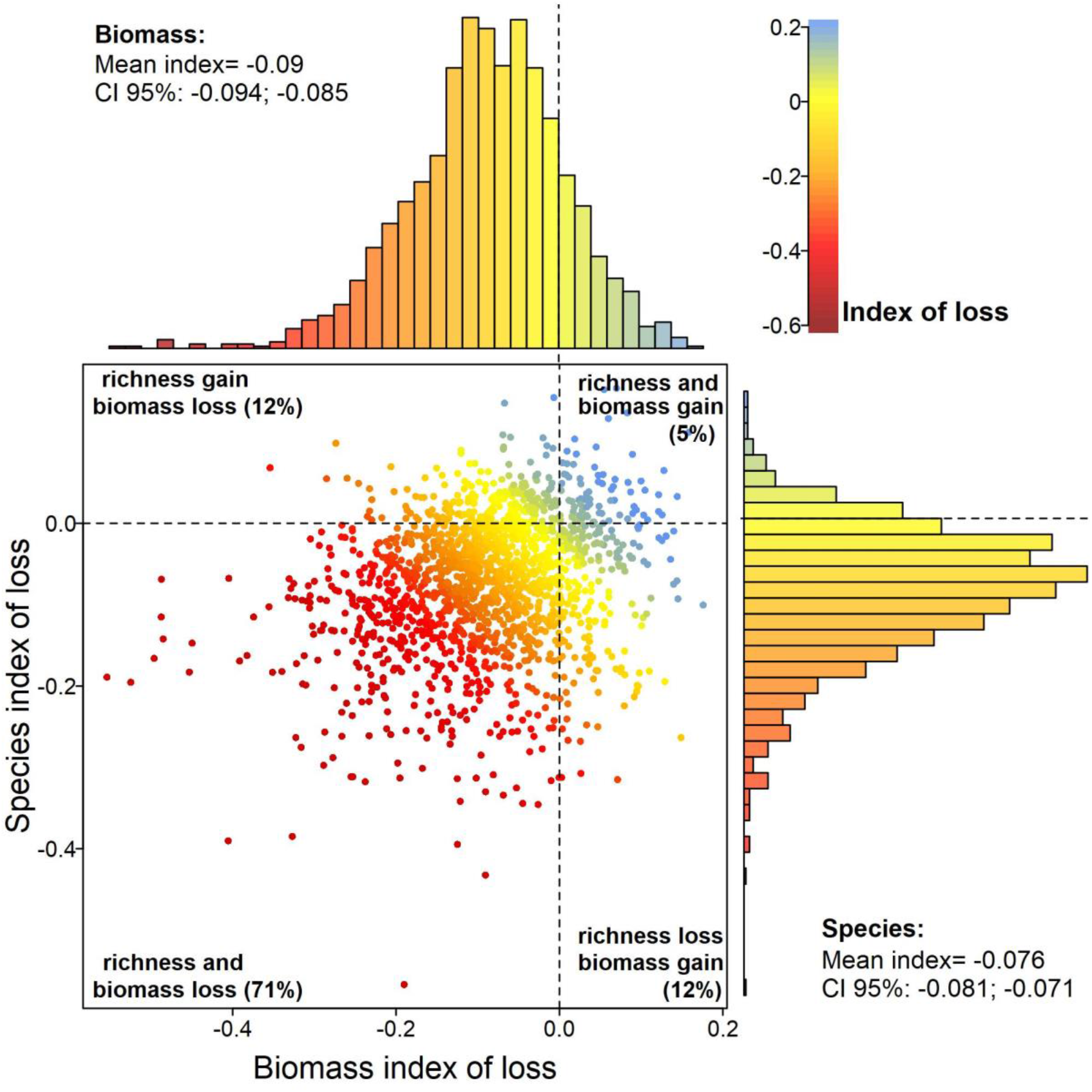
The relationship between losses of species richness and forest biomass due to human-induced impacts. The standardized indices of loss were estimated for each forest survey (points), with biomass on the *x*-axis and species on the *y*-axis. By the margin of each axis, the distribution of the index is presented with the mean and the corresponding 95% confidence interval (CI 95%). Dashed lines separate negative indices (losses) from positive ones (gains). The index of loss is dimensionless and is highlighted by different colours ranging from dark red (high losses) to blue (gains). Pearson’s correlation coefficient between the two indices was 0.22 (*p*< 0.0001).

**Table 1.** The magnitude of the loss of forest biomass, species richness and species properties due to human-induced impacts. The average proportion of losses were obtained from the averages of the absolute loss predicted (predicted - observed values) for each survey normalized by the reference values of low-disturbance Atlantic Forest fragments (Supplementary Table 5). The average of the proportional absolute loss was weighted by the probability of having a surveyed fragment of the same size in the entire pool of Atlantic Forest fragments, a probability obtained from a lognormal distribution fitted to the size distribution of the ~250,000 forest fragments. This procedure was conducted separately for each biogeographical region of the Atlantic Forest and then averaged across regions weighted by the area of each region.

| Forest descriptor | Dbh cut-off (cm) | Absolute loss (%) |
| --- | --- | --- |
| Forest biomass | ≥5.0 | 25.3 |
|  | ≥10.0 | 32.0 |
| Tree species richness | ≥5.0 | 30.9 |
|  | ≥10.0 | 22.9 |
| Species properties |  |  |
| Wood density | ≥5.0 | 2.2 |
| Max. adult height | ≥5.0 | -1.9 |
| Seed mass | ≥5.0 | 35.7 |
| Ecological group | ≥5.0 | 32.6 |
| Extinction level | ≥5.0 | 25.2 |
| Endemism level | ≥5.0 | 42.1 |

Human-induced impacts also caused a decline in the abundance of late successional, large-seeded, and endemic species (Supplementary Fig. 9, Supplementary Table 4), with reductions of 25-42% when compared to low-disturbance Atlantic Forests (Supplementary Table 5 and 6). Shifts in species composition caused by human impacts have been reported for tropical forests, including the Atlantic Forest^3,7,19^. Here we also found greater shifts in species properties in surveys with greater losses of species richness and biomass (Fig. 2, Fig. 10-12). This means that the erosion of richness and biomass is being accompanied by a parallel decline of species that can enhance the provision of ecosystem services^24,25^ and safeguard the conservation value of the Atlantic Forest. In the long run, these losses can reinforce each other^24^, leading to an even greater erosion of biodiversity and biomass.

**Figure 2.**
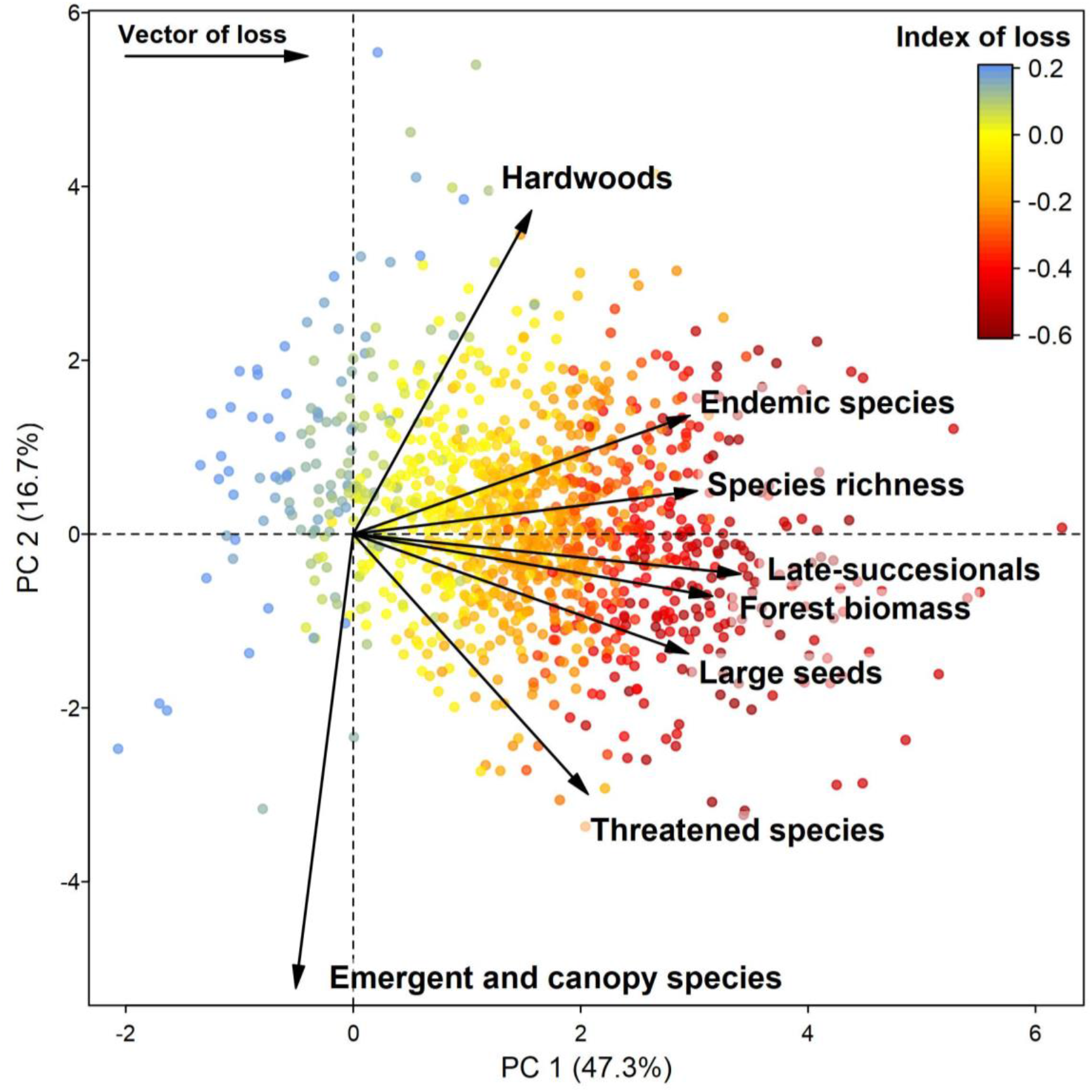
Human-induced impacts on tree species richness, multiple species properties and forest biomass. Two-dimensional representation of the standardized indices of loss for each forest survey (points) plotted on the two first principal component (PC) axes, which explain 64% of the variation in the indices of loss. Arrows and their length represent the vectors of loss, i.e. the direction and strength of each individual index of loss. The more aligned the arrows, the more correlated the indices are. The average of all indices of loss is highlighted by a colour scale for each survey ranging from dark red (high losses) to blue (gains).

**Figure 3.**
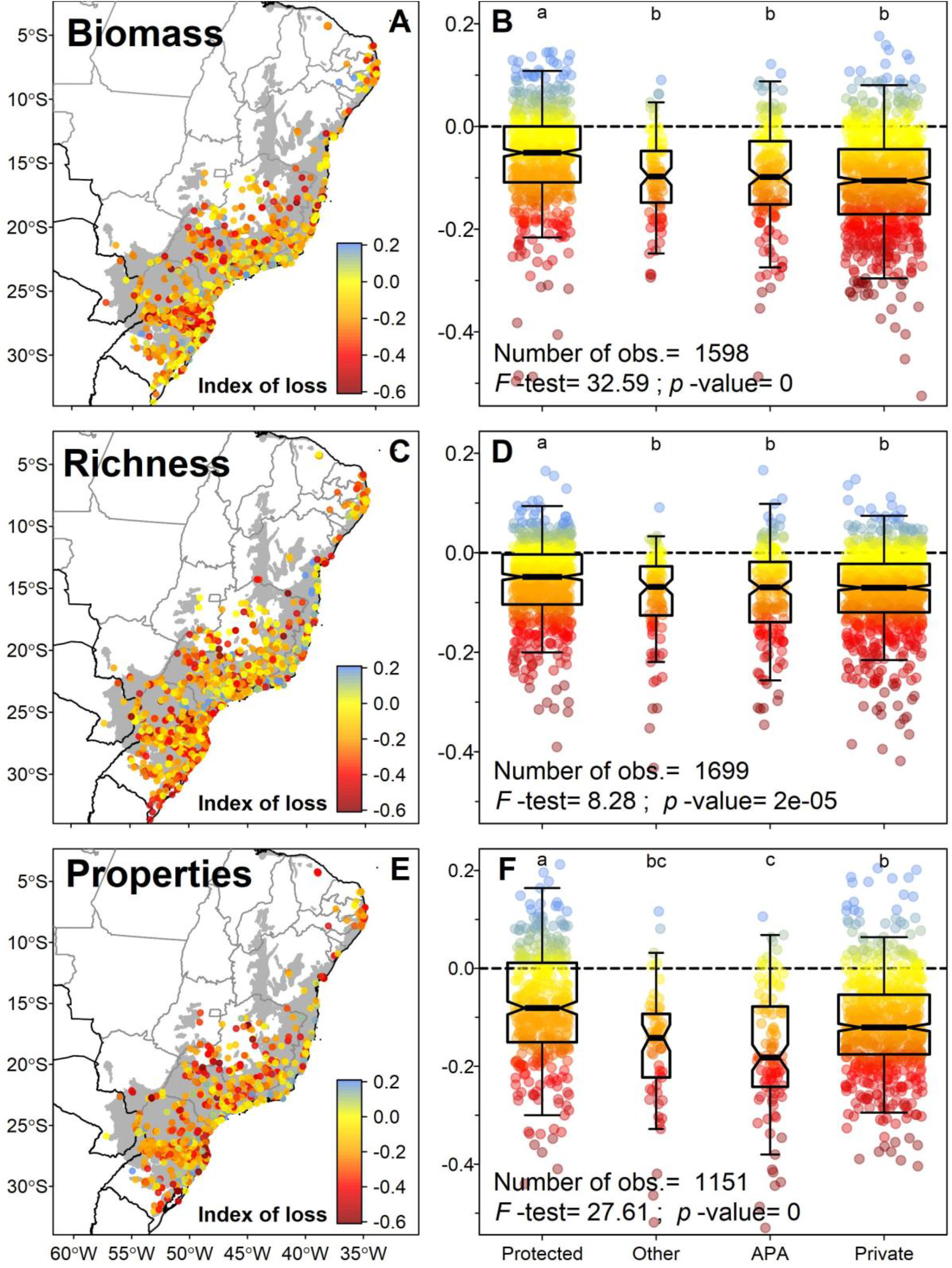
The distribution of the indices of loss across the Atlantic Forest and the effects of land conservation category. Each point represents one forest survey for which the index of loss was estimated for forest biomass (A and B), tree species richness (C and D) and species properties (E and F). In panels A, C and E, the shaded area is the original extent of the Atlantic Forest. In panels, B, D and F, the box-and-whisker diagrams summarize the distribution of the indices of loss for each land-use category (*i.e*., bold centre line, median; box limits, upper and lower quartiles; whiskers, 1x interquartile range), organized from left to right in decreasing order of protection. The *F*-statistics and number of observations of the models are presented, as well as the result of the Tukey test for difference among group means (lowercase letters). Degrees of freedom: 1579 (panel B), 1698 (panel D) and 1150 (panel F). Legend: Protect indicates Strict protection and Sustainable Use conservation units; Other indicates other public and protected lands (*e.g*., research centres, university campuses, botanical gardens, military and indigenous land); APA indicates private land inside areas with sustainable use of natural resources, locally known as “Environmental Protection Areas”; Private indicates private land.

The highest impact we found was related to the decline in the abundance of endemic species, meaning that endemics are being replaced by species with wider geographical ranges. Because species with wider ranges tend to occur over a wider variety of environmental conditions (*i.e*., generalists)^26^, widespread species can benefit at the expense of endemic and more specialist species in degraded fragments. Eventually, this process leads to the decrease of beta-diversity through time and thus to the biotic homogenization^19,27–29^. Thus, our results support the argument that human-induced impacts are driving the biotic homogenization of Atlantic Forest fragments^19,30^. Our approach based on CWMs of species properties, does not allow us to distinguish which groups of species are causing this homogenization, but we can draw some propositions. Widespread tree species in the Atlantic Forest are often small-seeded pioneers that proliferate at forest edges and small fragments^19,31,32^. However, we found only a weak correlation between the loses of endemism level and of seed mass and ecological groups (*r=* 0.14 and 0.17, respectively; Supplementary Fig. 12). This suggests that not all species proliferating in disturbed Atlantic Forests are small-seeded pioneers. In addition, because exotics represented only 0.3% of the trees in our data set, the proliferation of widespread native species is the most probable cause of the Atlantic Forest homogenization.

Wood density and maximum tree height, both related to carbon storage potential, presented the smallest changes. The latter even presented a positive mean index of loss, which may be explained by an increase in the abundance of late-successional, understorey species as human-impacts decrease. In the Atlantic Forest, species with those characteristics (common within Celastraceae, Erythroxylaceae, Myrtaceae, and Rutaceae) often have wood densities above 0.7 g cm^-3^, explaining the negative correlation found between maximum height and wood density losses (Fig. 2, Supplementary Fig. 12). Although taller species often have larger seed mass and higher wood density across vascular plants^33,34^, within trees there is still a wide spectrum of trait variation related to different regeneration strategies. Tree species that demand high irradiance for their development tend to have smaller seed sizes than species able to regenerate under mature canopies^35^, which was confirmed here by the relatively high correlation (r= 0.51) between the indices of loss of seed mass and ecological groups (Supplementary Fig. 12). Moreover, some canopy and emergent trees are long-lived pioneers, which have relatively light wood and small seeds^35,36^ (*e.g., Albizia, Ceiba, Ficus, Gallesia, Jacaratia, Parkia, Phytolacca, Piptadenia, Tachigali*). Altogether, these results suggest that declines in forest carbon stocks can be more easily explained by changes in forest structure than in its trait composition^37^.

### Influence of protected areas

Human-induced losses were lower inside than outside protected areas (Fig. 3, Supplementary Fig. 13) and they decreased as the size of the protected area increased (Supplementary Fig. 14). Thus, large protected areas are important to reduce human-induced degradation, besides preventing deforestation itself. However, even inside protected areas, we detected pervasive losses of species richness, species properties and forest biomass (Fig. 3), revealing the practical limits of conservation policies focused solely on the establishment of protected areas^16^. In addition, human-induced impacts in areas where human settlements and use of resources are allowed but regulated (*i.e*., the Brazilian “Environmental Protection Areas”) were as high as, or higher than, in other areas of private land (Fig. 3). Thus, not all types of protected areas are equally effective at conserving biomass and biodiversity^15^. This pervasiveness of human impacts, irrespective of land protection category, implies that strengthening the regulation within and surrounding existing protected areas is as important as the creation of new protected areas^16^. Although the land conservation category explained a small portion of the variation in all indices of loss, this factor explained more variation than other metrics of human pressure (Supplementary Fig. 13). This means that the magnitude of biodiversity and biomass loss is difficult to predict in space based on human presence indices (see Supplementary Discussion). Moreover, we found differences in average loss across the Atlantic Forest biogeographical regions for most indices of loss (Supplementary Fig. 15), reinforcing the idea that strategies to overcome biodiversity and carbon loss in the Atlantic Forest should be planned regionally^30^ (see discussion below).

### Total carbon losses in the Atlantic Forest

The erosion of biodiversity and biomass was pervasive across the entire Atlantic Forest hotspot. We projected the biomass loss across the remaining Atlantic Forest area and obtained losses of 451-525 Tg of carbon - equivalent to the deforestation of 55-70 thousand km^2^ (Table 2, Supplementary Fig. 16, Supplementary Table 7). This is about 1.4 times the carbon loss due to deforestation of the Atlantic Forest between 1985 and 2017, and about a quarter of its remaining forest area (26%). Assuming a value of US$5 per Mg C in international markets, this human-induced carbon loss translates into US$2.3–2.6 billion in carbon credits alone (see Supplementary Methods). Although carbon-related impacts can be priced, biodiversity loss is far more difficult to value. The loss of a species locally does not necessarily translate into regional extirpation. Moreover, impacts on tree diversity may be protracted over decades^5^ and the recovery of tree diversity and species composition occurs at a much slower rate than forest biomass^9^. Biodiversity losses can be valued indirectly from their negative impacts on ecosystem services^20,24,25^, but the underpinning of these relationships are currently not understood well-enough to provide accurate economical assessments, particularly regarding shifts in species properties within ecological communities.

**Table 2.** The average loss of forest carbon and its projection to the remaining Atlantic Forest area. The proportion of carbon loss was obtained per biogeographical region and then multiplied by the remaining forest area of each region to obtain the equivalent forest loss, *i.e*., the Atlantic Forest area that would match the carbon losses caused by post-deforestation human impacts. The equivalent carbon loss was computed based on the equivalent forest loss and the reference values of carbon storage per biogeographical region. We assumed a value of US$5 per Mg C paid for carbon credits obtained from projects of forestry and land use.

| **Dbh cutoff (cm)** | **Carbon loss (%)** | **Equivalent forest loss (km<sup>2</sup>)** | **Equivalent carbon loss (Tg C)** | **Carbon credits (Billion US\$)** |
| --- | --- | --- | --- | --- |
| ≥5 | 25.3 | 55,082 | 451.3 | 2.257 |
| ≥10 | 32.0 | 70,279 | 524.5 | 2.622 |

### Implications for the Atlantic Forest restoration

To explore strategies to mitigate biodiversity and carbon losses in Atlantic Forest remnants, we used our models to compare the outcomes of two contrasting restoration strategies: ‘fragment restoration’ that aimed at halving the within-fragment disturbance levels of Atlantic Forest remnants; and ‘landscape restoration’ that aimed at restoring 20% of landscape forest cover around them. The first scenario aimed at simulating restoration activities such as the reduction of forest-edge effects, control of invasive species and enrichment plantings, while the second aimed at increasing fragment size and thus landscape connectivity, *e.g*., forest corridors (see Supplementary Methods for details). We found that proportional gains were greater for the ‘fragment restoration’ scenario for all variables, with the exception of wood density, maximum height and endemism levels (Fig. 4). This means that the most effective strategy to restore the remaining Atlantic Forest fragments is to reverse forest degradation inside them. This statement was particularly true for forest biomass, seed mass, ecological groups and extinction level (Fig. 4). Despite being an expected result, it reinforces the role of forest disturbances as an important driver of biodiversity and biomass losses^13,14^.

**Figure 4.**
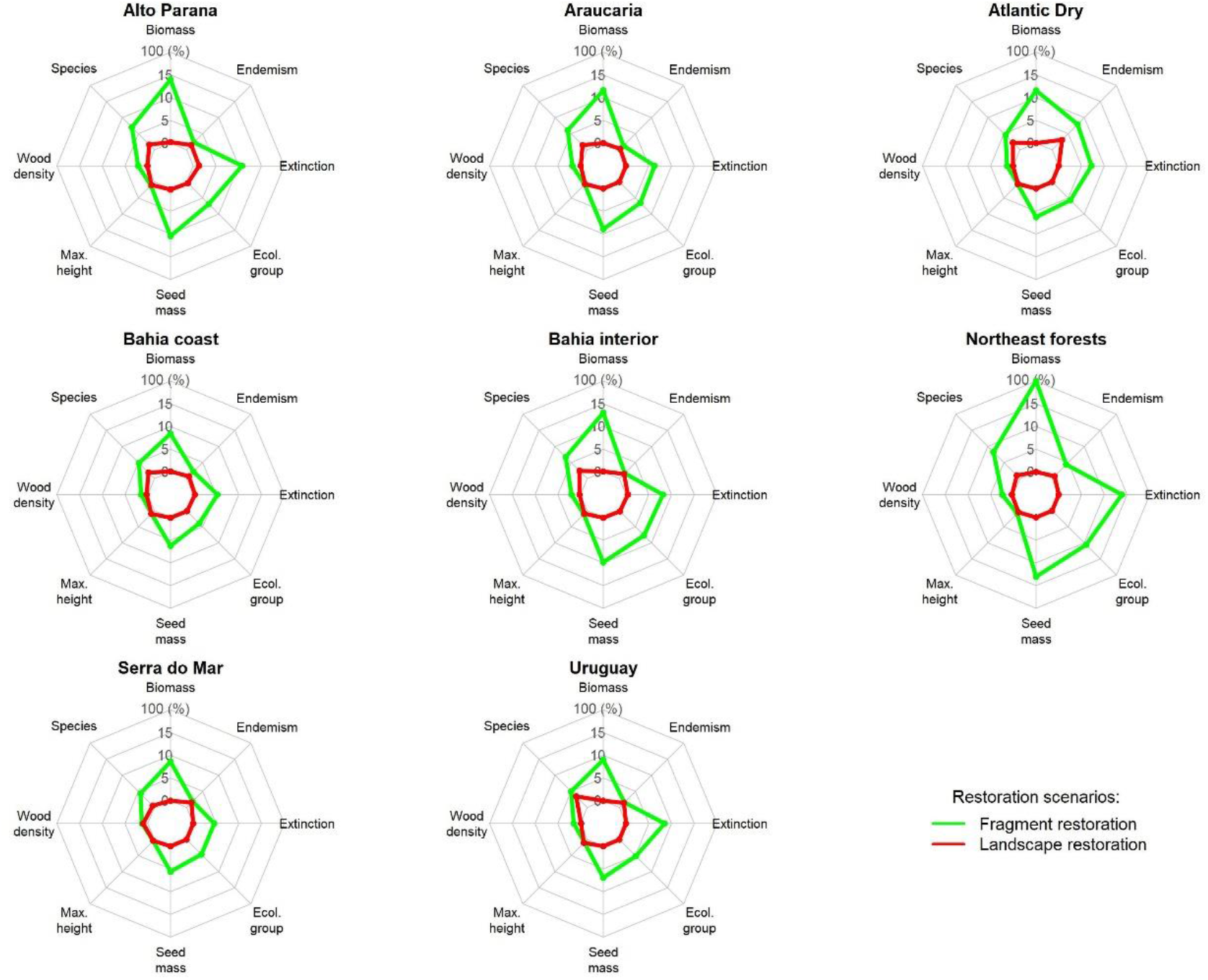
Multiple outcomes of two restoration scenarios for biogeographical region of the Atlantic Forest. In each radar plot, we provide the proportional gain of forest biomass, species richness, wood density, maximum adult height, seed mass, ecological groups, extinction, and endemism levels for the ‘Fragment restoration’ (green line) and ‘Landscape restoration’ scenarios (red line).

The ‘landscape restoration’ scenario, however, can recover much more carbon at the landscape-scale, since it includes the restoration of non-forest lands. Predicted carbon gains and restoration yield (*i.e*., carbon gains divided by total costs of restoration) for the ‘fragment restoration’ scenario were about 5-9% and 19-33% of the gains and yield for the ‘landscape restoration’ scenario (Supplementary Table 6), with restoration outcomes depending on the fragmentation and disturbance levels of the different Atlantic Forest regions (see Supplementary Results and Discussion). The ‘landscape restoration’ strategy can thus sequester carbon more efficiently, but it had little impact in reducing forest degradation within the remaining forest fragments (Fig. 4). Thus, restoration will require regionally-planned strategies that combine landscape with fragment restoration to efficiently attenuate biodiversity and carbon losses in human-modified tropical forests^30^.

The two restoration scenarios predicted small improvements for hardwood, tall and/or endemic tree species within fragments (Fig. 4). Greater improvements would probably require the restoration of fragments to their full pre-disturbance conditions and targeting landscape restoration to forest covers much larger than 20%. A 50% of landscape forest cover can decrease the uncertainty in restoration success^38^, but establishing such a high target for restoration of rural Atlantic Forest landscapes would probably not be economically feasible. In this context, selecting the appropriate species to maximize restoration outcomes becomes a more efficient strategy^39^. To help the species selection for restoration we provide a list of 242 common trees species (Supplementary Data 2) that have a high potential to increase the carbon storage and the conservation value of disturbed Atlantic Forest fragments (see Supplementary Methods for details). The re-introduction of these species could mitigate effects of local extirpation in the short term, while the connectivity of the landscape can be restored and allow the frugivore fauna to return in the long run. Seedlings from hardwood, tall or endemic species may be less available from nurseries and thus increase restoration costs^39,40^. So, local seed production cooperatives and nurseries able to propagate these species will be essential for the support of the Atlantic Forest restoration^39^, particularly regarding the re-introduction of large-seeded Atlantic Forest endemics.

### Implications for biodiversity and carbon conservation

The imprint of the human-induced degradation on the Atlantic Forest may be an indicator of the future condition of other tropical forests. Thus, the fate of tropical forests depends not only on avoiding the deforestation of intact forests or on promoting the reforestation of degraded lands^9,12^. It also depends on mitigating forest degradation in the remaining forest fragments^10,13,17,25^. The impacts of forest degradation are hard to quantify at regional scales and have therefore received less priority in the climate change and conservation agendas. Despite recent changes in the Brazilian government’s environmental priorities, the Brazilian pledges to mitigate greenhouse-gas emissions (*e.g*., Atlantic Forest pact, Paris Agreement or Bonn challenge) do not include actions to reverse forest degradation or to re-introduce species with certain properties (*e.g*. species with a high conservation value or carbon storage potential). In human-modified landscapes, the mitigation of forest degradation can be a more cost-effective strategy than reforestation depending on the outcomes desired. Moreover, it conflicts less with other land uses, such as agriculture^41^ and thus has a good potential for engaging stakeholders.

Therefore, besides safeguarding part of the biodiversity and catalysing the restoration of nearby areas^42^, restoring disturbed fragments is an opportunity to enhance tropical biodiversity and biomass. In regions such as the Atlantic Forest, where most of the forest remnants are on private land^21^, this opportunity has direct ramifications implications to the compensation mechanisms for climate change mitigation. In Brazil, funds to reduce carbon emissions from deforestation and forest degradation (REDD+) are mainly concentrated on Amazonia and focused on avoiding deforestation (*e.g*., the currently inactive Amazon Fund). Currently, only the State of Rio de Janeiro has a fund for the Atlantic Forest (http://www.fmarj.org). Our results indicate that the potential for a biome-wide compensation fund is in the order of billions of dollars. The creation of national policies to mitigate forest degradation, although highly dependent on the vision and will of politicians, could be the key to attracting funds to the Atlantic Forest and therefore be decisive for the future of this biodiversity hotspot^43^.

## Methods

### Forest survey and species data

We obtained 1,819 tree community surveys of natural Atlantic Forests available from the Neotropical Tree Community database (TreeCo)^23^. We considered all types of forest formations in Brazil, Paraguay and Argentina (Supplementary Fig. 1, Supplementary Data 1), with the exception of dry deciduous and early secondary forests. Moreover, we considered only surveys including trees with diameter at breast height (dbh) ≥3, ≥5 and ≥10 cm and using plots or the point-centred quarter method. Surveys represented a total effort of 1.45 million trees and 1,238 hectares (Supplementary Table 1) and they covered a wide spectrum of environmental conditions and patch/landscape metrics (Supplementary Table 2).

For each survey we extracted the tree density (trees ha^-1^), basal area (m^2^ ha^-1^), species richness, sampling method (plots and point-centred quarter method), arrangement of sample units (contiguous and systematic/random), sampled area (ha), dbh inclusion criteria (cm) and geographical coordinates. We verified the precision of the geographical coordinates provided to ensure that they corresponded to the forest fragment studied, otherwise the survey was discarded. Whenever needed, plot coordinates were corrected, based on maps or the site description provided in the study, including internet searches of any valuable information on the fragment, farm or park location.

We compiled data on species abundances and basal area for surveys with a sampling effort ≥0.1 ha and with species data presented in an extractable format (see Supplementary Methods). For all surveys providing information on voucher specimens associated with morpho-species, we checked for identification updates performed by taxonomists using the speciesLink network (http://splink.cria.org.br). Although only about one-third of the surveys provided vouchers, this effort improved the taxonomic resolution of around 10% of our records. We checked species names for typographical errors, synonyms, and orthographical variants, following the Brazilian Flora 2020^44^. Names marked as *confer* were assigned to the species suggested for confirmation, while those marked as *affinis* were considered at the genus level. We compiled 98,030 species records, totalling 1,171,935 trees measured and 3,124 valid species names.

We compiled information on the following species properties: wood density (g cm^-3^)^45^, maximum adult height (m)^46^, seed mass (grams)^47–49^, ecological group (or successional group, *i.e*., pioneer, early-secondary, late-secondary and climax), IUCN threat category^50^ and endemism level. Ecological groups were obtained from information provided in the original surveys and from the specialized literature. Endemism levels were defined based on species occurrences in different continents, countries and Brazilian states^44,46^. Species with Pantropical, Neotropical and South American distributions were classified as ‘not endemic’, while species restricted to one or two adjacent regions (*e.g*., São Paulo and Rio de Janeiro states) were classified as ‘local endemic’. Species restricted to South, South-eastern or North-eastern Brazil were classified as ‘regional endemic’. Over 300 different sources of information were consulted to complete these species properties (see Supplementary Notes). Records retrieved for numerical properties were averaged for each species. Maximum adult height was considered here as the 90% quantile of the distribution of maximum height records. For wood density and seed mass, we used genus-level instead of species-level averages if necessary^51,52^. We also completed missing information on ecological groups for typical pioneer Neotropical genera (*e.g*., *Cecropia, Trema, Vernonanthura)*.

We computed community weighted means (CWM) for the species properties in order to summarize the community composition *sensu lato* of each survey. For continuous species properties, the CWM was simply the average of each property weighted by the total number of individuals in the community. For wood density and maximum height, we used total basal area instead of the number of individuals of each species to calculate the CWM. For wood density, CWM was obtained after removing palms, palmoids, cacti, and tree ferns. For maximum height, we removed shrubs prior to the calculation of CWM. For seed mass, we removed tree ferns prior to the calculation of CWM. To compute the CWMs for the categorical properties, we treated them as ordinal categorical data: Pioneer < Early secondary < Late secondary < Climax for ecological groups; Not threatened/Not evaluated/Least concern < Data deficient/Near threatened < Vulnerable < Endangered < Critically endangered for extinction level; and Exotic/Naturalized < Not endemic < Northern/Eastern/Southern South-America < Regional endemic < Local endemic for the endemism level. We assigned scores to these ordered categories (see Supplementary Methods) and used them to compute the CWM. These scores were chosen so that the higher the CWM, the better the community is regarding species properties.

### Site descriptors

For each survey we obtained climatic (*e.g*., mean air temperature and rainfall) and topographic information (*i.e*., altitude, slope declivity and aspect) from different sources (see Supplementary Methods). We also obtained soil classes from the original surveys, which were checked for their consistency using soil maps at state and national scales. Missing data and inconsistencies between sources were double-checked to assure soil data quality and homogeneity. We used soil classes to infer average soil properties for plant growth by cross-referencing them with a database of physical and chemical soil properties (see Supplementary Methods). We used forest cover maps (30 m resolution)^1^ to extract 4×4 km landscapes centred on the survey coordinates and we used a 70% canopy closure threshold to classify maps into forest or non-forest pixels. Classified maps were used to calculate the proportion of forest cover and core forest cover, the median edge-to-edge distance between fragments and different landscape aggregation indices (see Supplementary Methods for details).

Fragment size was obtained from the original publications and was cross-validated using the 2002 and 2012 maps of the Atlantic Forest fragments^53^. We completed missing values of fragment size if there was consistency of the fragment size obtained from these maps with the one obtained from the classified 4×4 km landscapes. Inconsistencies between these sources were solved in Google Earth Pro (© Google Inc.), including the manual re-calculation of fragment size, which we conducted for ~400 surveys. The computation of the (mean) distance between surveys and forest edges was not possible for the majority of surveys for different reasons (see Supplementary Methods for details). Forest-edge effects are inextricably linked to fragment size and shape. Thus, their effects are indirectly accounted for in other landscape and patch metrics.

Forest disturbance level was assigned based on the information on the type, intensity and timing of human disturbances (*i.e*., selective logging, fire, hunting, thinning) provided by the authors of the surveys. We considered three levels of disturbance: high (*i.e*., highly/chronically disturbed forests, typically disturbed less than 50 years before the survey); medium (lightly/sporadically disturbed forests, and/or disturbed 50–80 years ago); and low (forests left undisturbed for at least 80 years). This classification is qualitative, with substantial variation in forest structure and diversity expected within classes. However, more objective and detailed information on disturbance histories, main disturbance types and their intensities was lacking (see Supplementary Methods). Thus, these coarse classes are the best information available to allow analysis across the Atlantic Forest.

We assigned each survey a biogeographical region^54^. To avoid an excessive subdivision of regions, we reassigned Atlantic Forest surveys mapped as *Cerrado, Caatinga, Campo Rupestre* enclaves, Atlantic Coast *Restingas* and Southern Atlantic Mangroves to the closest region (Supplementary Fig. 1). We included the Uruguayan biogeographical region, representing the forests in the transition to the pampas region of South Brazil as part of the Atlantic Forest^21,53^. The proportion of each region in our sample was similar to their contribution to the remaining Atlantic Forest area with few exceptions (Serra do Mar: 29% in the data set and 21% of remaining area, Araucaria: 24% and 13%, Alto Paraná: 21% and 31%, Bahia Inland: 8% and 11%, Bahia Coastal: 6% and 7%, Uruguay: 5% and 3%, Northeast: 5% and 4%, Atlantic Dry: 2% and 10%).

Categories of land conservation were obtained from the original study and/or from maps of conservation units, as follows: protected areas (*i.e*., strict protection and sustainable use conservation units); other public and protected land (*e.g*., research centres, university campuses, botanical gardens, military and indigenous lands); private land regulated by government laws, a conservation unit known in Brazil as Environmental Protection Areas (or APA for Portuguese: “Área de Proteção Ambiental”); and private lands outside conservation units (*e.g*., farms). We could not assign the land-use category for 5% of surveys. For surveys conducted inside protected areas, we also obtained the size and type of the protected area (*e.g*., strict protection, sustainable use).

We extracted the Human Influence Index (HII)^55^ based on the coordinates of each survey. The HII ranges from 0 to 64 (minimum and maximum human influence, respectively) and it accounts for human population density, land use (*e.g*., urban areas, agriculture) and ease of access (*i.e*., proximity to roads, railways, navigable rivers, and coastlines). The survey distance from main cities (>100,000 inhabitants) was significantly correlated with the HII (Pearson’s *r*= −0.38, *p*< 0.0001). Because results using this distance instead of the HII were qualitatively the same, we present only the results using the HII.

### Data analysis

We conducted separate analyses for each response variable, because not all surveys had data on forest biomass, species richness and CWM simultaneously. We also removed 50 surveys from the analyses of species properties, because in these surveys less than 80% of the sample individuals had information on species properties. Thus, analyses were conducted using 1676 surveys for biomass (92% of surveys), 1790 for species richness (98%) and 1213-1214 (68%) depending on the species property considered (Supplementary Data 1). Response and explanatory variables were transformed if necessary and candidate explanatory variables were pre-selected based on their co-linearity and on their impact on model performance (see Supplementary Methods).

We described forest biomass, species richness, and CWMs of species properties using linear mixed-effects regression models. To select which explanatory variables should be included in these models^56,57^, we first selected the co-variables composing the random structure of the models, which had survey methodology and biogeographical regions as candidate random effects. These two categorical variables divide the observations into groups, in order to account for correlated observations in the data^56,57^ (*e.g*., species richness is higher and more similar for some biogeographical regions when compared to other regions). Survey methodology refers to the sampling method, arrangement and dbh inclusion criteria, which were combined to create a methodological categorical variable (*e.g*., contiguous plots dbh ≥5 cm, systematic plots dbh ≥10 cm, etc). Exploratory data analyses suggested an interaction of the effects of sampling effort and the methodological variable on forest biomass and species richness. Indeed, the addition of a random term composed by the log-transformed sampling effort nested within the methodological categories significantly improved the model fit for both variables. To avoid the artificial inflation of model explanation^57^, we also included log-transformed sampling effort as a fixed effect in the final models of biomass and richness. The same was not true for the random structure of the models of species properties, which had fewer observations and were more prone to overfitting issues (*i.e*., model singularity).

We then selected the fixed effects to compose the optimum regression model. We kept to a minimum the interactions between fixed effects to avoid problems with the interpretation of individual coefficients, including only interactions with a clear biological meaning (*e.g*., temperature × rainfall seasonality). The optimum regression models contained only fixed effects that improved overall model fit (see Supplementary Methods) and they had the following general structure: *y* ~ *Environment* + *Human* + *Effort* + (*Effort*|*Method*) + (1|*Region*). The term *Environment* includes the climate, topography and soil variables (and their interactions). *Human* includes landscape metrics, fragment size, and disturbance level. *Effort* is the total survey effort (not included in species property models). As described in the previous paragraph, *Method* is the methodological categories (nine levels) and *Region* is the Atlantic Forest biogeographical regions (eight levels, Supplementary Fig. 1). Therefore, models had environmental, human, methodological and biogeographical (historical) components. The number of individuals sampled had, as expected, a strong influence on species richness. During explanatory analyses, we also found a positive relationship between biomass and tree density (individuals ha^-1^). So, we included number of individuals and tree density as fixed effects to model richness and biomass data, respectively.

### Evaluation of human impacts

We used the optimum models to quantify human-related impacts on species richness, species properties and forest biomass. We first obtained the model predictions in a scenario without major human-induced impacts, *i.e*., model predictions for human-related variables reset at values corresponding to large (>300,000 ha), low-disturbance patches in landscapes with 100% of (core) forest cover and maximum patch aggregation. We assumed current conditions for all other co-variables in the models (*i.e*., climate, soil conditions, sampling method and biogeographical region) and the deforestation caused by indigenous populations to be negligible. Then, we calculated the standardized difference between observed values and predicted values in the human-free scenario:

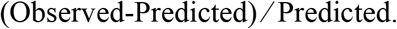

The difference between observed and predicted values was standardized to make them comparable across our response variables. We used this standardized difference as an index of loss related to human impacts (Supplementary Fig. 9). Negative indices mean that observed values of biomass, species richness or community weighted means (CWM) of species properties are smaller than the predictions in the human-free scenario at the same forest fragment.

Because nearly all Atlantic Forest surveys were conducted after the intensification of its deforestation (1940-1970s)^23,58^, it is impossible to obtain measurements of pristine forests, particularly at the scale of our study (~ 1.4 million km^2^). Instead, we assumed Atlantic Forests previous to human impacts to be large, fully-connected and undisturbed. This assumption has limitations but it is a valid yet conservative representation of past forest conditions. We compared the predictions for the human-free scenario against intervals obtained from 5000 samples of the estimated model coefficients (Supplementary Methods). The precision of these predictions was not strongly sensitive to variations in the parameter estimates (Supplementary Table 4). Predictions of carbon stocks for the human-free scenario ranged between 43-162 Mg C ha^-1^ for individual surveys, but they were typically between 88-110 Mg C ha^-1^ (mean of 99 Mg C ha^-1^), which is consistent with previous studies in the Atlantic Forest^58,59^. Finally, because we predict shifts in CWM of species properties and not in taxonomic composition itself, our approach did not provide any indication of which species increased or declined in abundance due to human impacts.

We inspected the relationship among indices of loss by plotting one against the other (Fig. 1, Supplementary Fig. 10-S11) and testing the strength of their relationship using linear regression models. We also inspected the correlation among the pairs of indices of loss for the six species properties (Supplementary Fig. 12). We averaged the indices of loss for the six species properties to generate a mean index of loss, using the conditional *R*^2^ of the mixed-effects models of each property as weights (Supplementary Table 3). We then explored the relationships between all indices of loss using the two first axes of a principal component analysis, produced using the scaled indices of loss for species richness, properties and biomass (Fig. 2). We also evaluated if the category of land use and of the Human Influence Index (HII)^55^ have an effect on these indices (Fig. 3, Supplementary Fig. 13). We plotted them over the Atlantic Forest map to reveal possible hotspots of human impacts. Finally, we evaluated the effect of the size (Supplementary Fig. 14) and type of the protected area on the indices of loss (see Supplementary Methods).

### Absolute losses and projections across the Atlantic Forest

For each biogeographical region, we estimated the mean values of species richness, species properties and forest biomass for low disturbance surveys, taken here as references to the Atlantic Forest species diversity, community composition and biomass storage. Here, we assumed modern, low-disturbance fragments as proxies of past forest conditions. We acknowledge that past forests may have had more biodiversity and biomass than modern-day forests, but these fragments represent the best information currently available for the Atlantic Forest, given that measurements prior to human impacts are unavailable. Thus, we took a conservative approach; if modern-day fragments underestimate past conditions, absolute losses would be even greater than reported here.

These reference values were obtained using a different subset of surveys for each forest descriptors and separately for each dbh inclusion criterion (Supplementary Table 5 and 6). Next, we calculated the absolute loss for each survey (*i.e*., Predicted - Observed), which was then averaged for each region. Because our sample is biased towards large forest fragments (Supplementary Fig. 2), these averages were weighted by the probability of having a fragment of the same size in each biogeographical region. These probabilities were obtained from log-normal distributions fitted to the 2016 size distribution of the remaining Atlantic Forest fragments^53^. We then calculated the proportion that these weighted average losses represent in respect to the region-specific reference values (Table 1, Supplementary Fig. 15).

To obtain the forest area that would match the carbon losses caused by post-deforestation human impacts (*i.e*., equivalent forest loss), the proportion of carbon loss was multiplied by the remaining Atlantic Forest area per biogeographical region^53^. Due to missing information on human-related variables for all Atlantic Forest fragments (e.g. within forest disturbance level), we assumed the same carbon loss for all fragments within each biogeographical region. Next, we obtained the total amount of carbon loss (*i.e*., the equivalent carbon loss), which was computed based on the equivalent forest loss and the reference values of carbon storage per region. The mean carbon storage, and proportional losses were averaged across the biogeographical regions, using the area of regions as weights. In contrast, values of equivalent forest and carbon loss were summed across regions. Thus, our estimate of carbon loss and its projection across the Atlantic Forest takes into account the bias towards larger fragments in our sample and the regional differences in Atlantic Forest carbon storage potential (Table 2).

We compared the equivalent forest loss to Atlantic Forest deforestation between 1985 and 2017^53^. In the Brazilian Atlantic Forest, deforestation was 1.93 million ha for the entire interval of 1985 and 2017^53^. For Paraguay, we found estimates for 1989-2000 and 2003-2013^60,61^, so we estimated a 1.96 million ha forest loss for 1985-2017 from the reported annual deforestation rates. For Argentina, we had estimates for 1998-2014^62^ and used the same procedure to obtain an estimate of 0.1 million ha for 1985-2017. It should be noted that overall deforestation in Brazil was larger than in other countries, but it occurred more intensely before 1985. Therefore, we estimated a total forest loss of 4.18 million ha between 1985-2017. Finally, we calculated how much money the equivalent carbon loss would represent if it had been traded as carbon credits in international markets (Table 2), assuming US$5 per Mg C paid for projects of forestry and land use^63^.

### Simulating strategies of forest restoration

We used the regression models fitted to biodiversity and biomass data to simulated two simple scenarios of forest restoration. The ‘fragment restoration’ scenario assumes no increase in landscape forest cover and aims at restoring forest disturbance levels to 50% of the lower level observed for the Atlantic Forest. This is realistic target, because restoring and maintaining disturbance levels below this threshold, although possible, would be too expensive. The ‘landscape restoration’ scenario mimics restoration activities around existing fragments, increasing fragment size and thus landscape integrity (*i.e*., greater forest cover) and connectivity (*i.e*., smaller fragment distance). This scenario implicitly assumes that restoration activities should focus on the restoration around existing forest fragments, which would increase the ‘area’ of restoration and thus maximize the success of the restoration project while minimizing its costs^64^. The ‘landscape restoration’ aimed at a minimum of 20% of landscape forest cover, a target that relates to the legal requirements for the Atlantic Forest fixed by the Brazilian Forest Code^65^.

The two scenarios were compared by generating model predictions for all the remaining fragments in 2016^53^. For each fragment, we compared the ‘current situation’ predictions to the predictions under each restoration scenario, by varying fragment disturbance level and patch/landscape metrics, respectively. Current situation predictions used fragment-specific coordinates to extract spatialized information (*e.g*., climate) or regional averages for information not available for all fragments (*e.g*., forest disturbance level). For the ‘landscape restoration’ scenario, the increase in forest cover to reach the 20% target and the corresponding changes in landscape metrics between the ‘current’ and ‘restored’ landscapes were derived from 100 iterations of simulated landscapes (see Supplementary Methods for more details). We compared the proportional gains of each restoration scenario for forest biomass, species richness, and CWMs of species properties (Fig. 4).

We also compared the costs and efficiency of the two scenarios in terms of carbon sequestration (Supplementary Table 6). We assumed an average value of US$750 per hectare for within-fragment restoration, which includes activities to reduce edge effects, control of invasive species, and enrichment plantings^66,67^. We assumed an average of US$1000 per hectare for land opportunity cost (10 years period for cattle-ranching activities)^68^ and an average of US$1500 per hectare for the restoration costs on degraded lands (from seedling plantation to assisted natural regeneration), which varies between US$350 and US$3000 per hectare for the Atlantic Forest^64,68–70^. However, we assumed different restoration costs for each biogeographical region (*i.e*., less disturbed and fragmented landscapes should cost less to be restored - Supplementary Table 6). As before, we monetarized the expected carbon gains in terms of carbon credits in international markets (US$5 per Mg C paid for projects of forestry and land use^63^). For the ‘landscape restoration’ scenario, we separated carbon gains into those expected inside the remaining fragments and in the restored areas *per se* (Supplementary Table 6). To calculate the potential of carbon recovery in the restored areas, we used the references of carbon density and the average carbon loss for each biogeographical region (Supplementary Fig. 15), assuming that the drivers of carbon loss in fragments would be similar in restored areas.

### Data availability

Survey, species abundances and trait data used in this study were extracted from the Neotropical Tree Communities database (TreeCo, version 4.0) and are available upon request at http://labtrop.ib.usp.br/doku.php?id=projetos:treeco:start. The list of surveys extracted from the TreeCo database is provided in Supplementary Data 1, together with the corresponding metadata. The sources of survey and species abundance data are referenced in Supplementary Data 1 and sources of species properties data referenced in the Methods or in the Supplementary Notes. Other relevant data are available from the corresponding author upon reasonable request.

### Code availability

Codes used to conduct the analysis is available from the corresponding author upon request.

## Acknowledgments

We thank Ary T. Oliveira-Filho, Danilo Mori, Markus Gastauer, Melina Melito, Natália Targhetta, Geison Castro, Luiz F.S. Magnago and Carolina Bello for their great help in constructing the database. We also thank André M. Amorim, Andréia A. Rezende, João A. Meira-Neto, João L.F. Batista, Márcia C.M. Marques, Maria T.Z. Toniato, Mariana C. Pardgurschi, Mario J. Marques-Azevedo, Ricardo R. Rodrigues, Robson L. Capretz and Victor P. Zwiener for providing published or unpublished data in digital format. Luiz F.S. Magnago provided functional trait information for several species, Mariano C. Cenamo and Mauricio A. Voivodic provided critical comments on the implications of our results, Luciana F. Alves and Simone Viera provided important sources on carbon stock estimates and Pedro, H.S. Brancalion, Ricardo R. Rodrigues, and Ricardo A.G. Viani provided useful advice on restoration costs in the Atlantic Forest. This research was funded by the grant 2013/08722-5, São Paulo Research Foundation (FAPESP). R.A.F.L. was supported by the European Union’s Horizon 2020 research and innovation program under the Marie Skłodowska-Curie grant agreement No 795114. A.C.V. and A.L.G. were supported by the Conselho Nacional de Desenvolvimento Científico e Tecnológico (CNPq), grant 312075/2013-8 and the Fundação de Amparo à Pesquisa e Inovação de Santa Catarina (FAPESC), grant 2017TR1922. J.C. acknowledges funding by “Investissement d’Avenir” grants managed by Agence Nationale de la Recherche (CEBA, ref. ANR-10-LABX-25-01 and TULIP, ref. ANR-10-LABX-0041). We gratefully acknowledge the hundreds of researchers, students and technicians that performed the field work, species identification and data analysis that resulted in the surveys included in this study. These surveys were funded by many different agencies (*e.g*., CAPES, CNPq, FAPESP, FAPEMIG, FAPERJ, FAPESC, etc.), but space prohibits publishing the complete list of grants. Some major plot initiatives used in this study were supported by grants 99/08515-0, 99/09635-0 and 04/04820-3 (FAPESP) and grant DE-FC26-01NT411151 (US Department of Energy).

## Author information

### Contributions

This study was conceived and designed by R.A.F.L., with the help of J.C. and P.I.P. Data were compiled and/or validated by R.A.F.L., G.R.P., A.L.G., A.C.V. Data analysis was conducted by R.A.F.L. and P.I.P, with the support of A.A.O. R.A.F.L. drafted the paper and all authors have revised the subsequent drafts. All authors have contributed substantially to the interpretation and discussion of the results.

## Supplementary information

Supplementary information is available for this paper.

