## Supplementary Information for "The erosion of biodiversity and biomass in the Atlantic Forest biodiversity hotspot"

[1] Departamento de Ecologia, Instituto de Biociências, Universidade de São Paulo. Rua do Matão, trav. 14, 321, 05508-090, São Paulo, Brazil.

[2] Naturalis Biodiversity Center, Darwinweg 2, 2333 CR Leiden, The Netherlands.

[3] Departamento de Ciências Naturais, Universidade Regional de Blumenau. Rua Antônio da Veiga, 140, 89030-903, Blumenau, Brazil.

[4] Departamento de Engenharia Florestal, Universidade Regional de Blumenau. Rua São Paulo, 3250, 89030-000, Blumenau, Brazil.

[5] Laboratoire Evolution et Diversité Biologique, UMR 5174 CNRS, Université Paul Sabatier, IRD. 118, route de Narbonne, 31062, Toulouse, France.

[6] Systems Ecology, Vrije Universiteit, De Boelelaan 1087, Amsterdam, 1081 HV, Netherlands

*

**This PDF file includes:**

### 1. Supplementary Methods

#### Study site

The Atlantic Forest is a global biodiversity hotspot that once covered 1.63 million km$^2$ mostly in Brazil (92% of the total area), but also in Paraguay (6%) and Argentina (2% - Extended Data Fig. 1). It covers a wide range of climatic and edaphic conditions, with forest types ranging from rainforests to seasonal forests, including cloud, swamp, and white-sand forests[71]. The Atlantic Forest has been suffering from deforestation and degradation for over 500 years. Today, it includes some of the largest cities in South America, with over 148 million people currently living within the Atlantic Forest limits[62]. Less than 20% of the original Atlantic Forest remains and the remnants are characterized by small (<50 ha), isolated and altered fragments[21]. Although deforestation started centuries ago, the main period of fragmentation history occurred in the beginning of the 20th century, and deforestation rates have declined in the past two decades[53]. Regional differences in human population density, land conversion patterns, and cultural aspects have created a great diversity of landscapes in terms of forest cover, fragment size, and disturbance levels[21,72].

#### Forest surveys

We assembled a database of 1,819 surveys by compiling a large number of published and unpublished references. The surveys ranged from 0.03 to 26 ha in sampling effort (mean ± standard deviation: 0.68 ± 1.46 ha), with the majority being conducted in evergreen and semi-deciduous forests (73%), the two main Atlantic Forest formations[71]. Sampling design and methods varied greatly across surveys, but this methodological variation was taken into account during the analyses (see below). We compared the environmental conditions of the 1,819 forest surveys in our sample with the same conditions available for all Atlantic Forest fragments mapped in 2016 by ref. 53. Our sample was representative of the range of climate variation within the Atlantic Forest domain, with a slight sampling bias towards forest fragments in colder and wetter climates, which is partly due to the exclusion of dry deciduous forests surveys (Supplementary Fig. 1, panels A-C). Our sample, however, was strongly biased toward larger forest fragments (Supplementary Fig. 1, panel D), a bias which was taken into consideration before the projection of our results to the entire Atlantic Forest area.

#### Survey methods

Surveys in the Atlantic Forest typically include tall shrubs, treelets, trees, palms, and tree ferns and they exclude lianas and hemi-epiphytes. Forest descriptors obtained for each survey were tree density, total basal area, and observed species richness. These descriptors were cross-checked for 70% of the surveys for which abundance and biomass per species were available. If descriptors were missing, or their estimation was in doubt, we recomputed them based on phytosociological tables (that provided total tree abundance and richness) or on tree diameter distributions (biomass), as reported in the original studies. We assumed that randomly and systematically distributed sample units have equivalent methodological effects on the estimation of forest biomass and richness[73].

#### Species data

To evaluate human-related impacts on species responses, we extracted data on species composition and abundances for surveys with a total sampling area of at least 0.1 ha, and complete phytosociological tables. The sample size cut-off ensures a minimum representativeness of the composition of the forests surveyed. The second filter is related to the fact that many quantitative surveys present the phytosociological table partially (*e.g.*, only the most abundant species are given) or not at all. Also, the table was sometimes given in full but without the information needed to run analyses (*e.g.*, tables presenting only the species name and its importance value). In those cases, we tried to contact the original authors of the publications asking for complete phytosociological tables, but the rate of success of our e-mails was low. Surveys including species data represented 72% of the 1819 surveys, but they contained 84% and 81% of the sampling cover and number of trees, respectively.

#### Species properties

For each of the 3124 species included in the database, we compiled three functional traits: wood density, maximum height, and seed mass. We also documented information about the successional status of the species, according to the following categories and scores: Pioneer=1, Early secondary= 2, Late secondary=3, Climax= 4. Next, we quantified the conservation status for all species, including the extinction threat status at national and global scales. In case of inconsistencies, we used the category provided in the national red list, and we scored this status as follows: Not threatened/Not evaluated/Least concern= 0, Data deficient= 0.5, Near threatened= 1, Vulnerable= 2, Endangered= 3, Critically endangered= 4 for extinction risk. Finally, we classified species into different endemism levels according to their occurrences in different continents, countries and Brazilian states. We scored species endemism level as follows: exotic/naturalized= -1, not endemic= 0, Northern/Eastern/Southern South-America= 1, regional endemic= 2, local endemic= 3. For the latter, we assumed that the presence of exotic species has a negative score of endemic species.

The proportion of species with information available (at species or genus level) was: 99.5% for wood density, 93.4% for maximum height, 95.9% for seed mass, 65.4% for ecological groups, 100% for extinction threat status and 96.9% for endemism level. These species descriptions were compiled from over 300 different sources of information (see Supplementary Notes) and they are related to their role in carbon storage (*i.e.*, provision of ecosystem services), ecological interactions and strategies (*i.e.*, ecological functioning) and biodiversity conservation (*i.e.*, species conservation value). Besides being relatively easy to obtain from the literature, these six characteristics had good coverage at the species level for Atlantic Forest species.

#### Site descriptors

Based on the verified survey coordinates we extracted the mean annual air temperature, relative humidity, and rainfall, as well as monthly temperature and rainfall at ~100 m resolution[74,75], which were used to compute temperature and rainfall seasonality[76]. We also obtained the climatic water deficit, environmental stress factor (both at ~4.5 km resolution)[77] and the 19 Bioclimatic variables (BIO1, BIO2, …, BIO19) from WorldClim version 2.0 (~1 km resolution)[78]. For 23 surveys from Paraguay and Argentina, sites not covered by refs. [74,75], temperature and rainfall were obtained from ref. 78. We also extracted mean values of percentage cloud cover (%) and frost day frequency (days, both at ~50 km resolution)[79]. We gathered elevation data from SRTM (~30 m resolution)[80] and used it to calculate slope declivity (range: 0–90 degrees) and slope aspect (1–360 degrees). We combined slope declivity and aspect with latitude to calculate the potential direct incident radiation (MJ $cm^{-2}$ $y^{-1}$)[81].

Based on the same verified coordinates, we obtained missing soil classes for 37% of the surveys from studies conducted at the same sites or from state- or locality-level soil maps, if there was no disagreement across state and national soil maps[82]. For forest types over specific soil classes (*i.e.*, white-sand and swamp forests), missing soil types were assigned irrespective of possible source conflicts. For each survey, we assigned a soil type classification at the finest taxonomic resolution possible, following the Brazilian classification[83]. To obtain soil properties from soil classes, we used a database with physical and chemical soil properties for ~6000 soil profiles[84]. For each soil type in the database, we averaged the sum of bases ($cmol_c$/kg), cation retention in clay ($cmol_c$/kg) and organic matter content (%) across soil profiles. Moreover, profiles were ranked from 0 (very restrictive) to 4 (very suitable for plant growth) based on soil drainage, depth, chemical fertility and aluminium toxicity[85]. The sum of these ranks was used as a soil quality measure varying from 0 (very unfit) to 16 (very suitable). Thus, because soil classes in our surveys had the same classification of the soil profile database[83], soil classes were associated with mean values of soil quality and fertility obtained from the soil profile database.

#### Landscape metrics

We used forest cover maps for the years of 2000 and 2010 (30 m resolution, https://earthenginepartners.appspot.com)[1] to extract 4×4 km landscapes centred on the verified coordinates of each survey. The 2000 map was used for studies between 1995 and 2005 (24% of the surveys) and the 2010 map for studies after 2005 (76% of the surveys). We assumed a scale of effect[86] of 2 km, given that the extent of processes affecting tree populations, such as species dispersal and forest edge effects, generally do not exceed 1–3 km[4,87–91].

We used a threshold of 70% of canopy closure to classify the 30×30 m pixels of the vegetation maps into forest and non-forest. The 70% threshold was selected based on the evidence of a gap between the 95% confidence intervals of the canopy closure found for open-canopy savannas and Atlantic Forest formations (R.A.F. Lima, unpublished data). Classified maps were used to calculate the proportion of forest cover and core forest cover, the median edge-to-edge distance between fragments and different landscape aggregation indices (*e.g.*, effective mesh size and division, aggregation and splitting indices)[92,93]. Regarding the landscape metrics chosen, the proportion of forest cover and the proportion of core forest cover of the landscape are related to landscape integrity, while the median distance between fragments and landscape aggregation indices are more related to landscape connectivity. The proportion of core forest cover in each classified 4×4 km window was derived using a 90 m threshold (3 times the vegetation map resolution) to delimit fragment edges and cores. The 90 m threshold was chosen because most of the forest edge-effects are concentrated in distances up to ~100 m[4,8,11,94].

Because of problems related to the distinction between natural and planted forests in the maps provided by ref. 1[95], we used a Tree Plantation map[96] to delineate forest plantations. To minimize differences in landscape metrics between the year of map and that of the forest survey, we only considered surveys conducted after the decline and levelling off in the Atlantic Forest deforestation rates[53]. This decline occurred after 1990 for Rio de Janeiro and Rio Grande do Sul states. For the states of Bahia, Goiás, and Mato Grosso do Sul deforestation rates remained high until 2000[53]. For the other countries and Brazilian states, we set 1995 as the cut-off year. Surveys conducted prior to the cut-off year but inside large (>1000 ha) conservation units or university campuses were kept in the dataset. We excluded surveys in fragments that were suppressed afterwards. Landscape metrics were extracted from maps using R[97] and the contributed packages *raster*[98] and *SDMTools*[93].

#### Forest fragment size

We assigned a forest fragment to each survey and obtained their size based on information from the original publications. Fragment size was cross-validated using other sources of information. We used vegetation maps with 30 m resolution[19], so forest fragments should be detectable down to a resolution of ca. 0.1 ha. Less than 1% of our surveys were in fragments <0.9 ha (10 times the resolution of the maps). Therefore, cross-validation, as well as obtaining missing fragment sizes, was performed with good confidence and accuracy. The smallest fragment was 0.2 ha (Supplementary Table 1), while the few large fragments (>50,000 ha) were concentrated in the Serra do Mar and

*Araucaria* eco-regions of São Paulo, Paraná, and Santa Catarina states. Fragment size is a good proxy for the habitat availability for species establishment and proportion of the fragment core area susceptible to landscape/edge effects[99]. We could not obtain an accurate (mean) distance between the sampling unit and forest edges, because this information was generally missing from the original publication. Moreover, about 60% of the surveys are aggregates of multiple plots scattered across the fragment, but the coordinates of each plot were not provided. For older surveys, in particular, we had no precise coordinates within the sampled fragment, and the centre of the fragment was then taken as the most likely coordinate (18% of the surveys). Finally, typographical errors in reporting the geographical coordinates or conversion errors also caused inconsistencies that could not be solved below the fragment scale.

#### Forest disturbance level

We classified the surveys into three levels of forest disturbance: high, medium and low. We assumed the following scores to each level, which can be interpreted as the time in years since the last disturbance: high= 40, medium= 70 and low=100. This ordered classification has support in the legal classification of the Atlantic Forest[100] and it implicitly assumes that highly disturbed forests can redevelop into medium and low disturbance forests through natural regeneration or restoration. It also assumes that low-disturbance forests can regress into medium or high disturbance levels depending on the type and intensity of disturbances[30,101,102]. Natural events can alter the structure and dynamics of forest fragments[4,103], but we assumed that these events were rare[104] and that the major drivers of forest disturbance are related to direct (*e.g*., clear-cutting, logging, and fire) and indirect human activities (*e.g*., invasive exotic species). If there were doubts about the classification of the forest disturbance level based on the information provided by the original author of the publication (about 10% of the cases), the density of typical pioneer Neotropical genera was used to assign the surveys to one of the three forest disturbance levels. We discarded a total of 136 surveys for which a forest disturbance level could not be unambiguously assigned. We were unable to refine this classification any further due to a lack of more objective and detailed information in the original publications. For most surveys, the only information provided was a simple mention of the forest successional stage (*e.g*., initial, intermediary or advanced).

#### Processing and selection of variables

Some of the response variables had left-skewed distributions, and they were transformed accordingly to meet the normality of the regression residuals. We used logarithmic and power transformations, using a Box-Cox procedure to choose between both transformations and to select the best exponent in the case of power transformations. We eventually transformed biomass, richness, and seed mass. We log-transformed some of the dependent variables, namely tree abundance, fragment size, slope declivity, soil fertility, which also had left-skewed distributions. We power-transformed frost frequency, air relative humidity, and the climatic water deficit. Before analysis, we standardized all numerical explanatory variables [(observed − mean)/standard deviation], so that all variables had a similar range of variation, making their estimated effects comparable.

All categorical variables, namely ecological groups, extinction and endemism levels, and forest disturbance level, were treated as ordinal categorical data (also known as rank-ordered data) since they represent ordered and directional changes in the forest. For these variables, we compared the impact of different scoring methods: naïve (described above for each variable), ridit, normal and log-normal scores[105,106]. We compared the performance of models fitted using these different scoring methods based on the assumptions of normality of the studentized residuals (Shapiro-Wilk test), homoscedasticity (Breusch-Pagan test) and the number of outliers (Bonferroni test). For disturbance level, the only explanatory categorical variable we had, we also compared the difference in the Akaike information criterion (AIC) of the models fitted including the different scoring methods. The use of the ridit scores resulted in models that had marginally or significantly better performance than other scoring methods. Therefore, ridit scores were used throughout the analysis for all categorical variables.

The selection of candidate explanatory variables to be included in the regression analysis was based on their performance in previous similar studies and on the absence of strong correlation (Pearson $r > 0.75$) or co-linearity with other candidate explanatory variables. We included mean annual temperature and rainfall, regarded here as the two basic climate variables, and different measures of their seasonality. We also included the climatic water deficit, environmental stress factor and the 19 bioclimatic variables. The measure of temperature seasonality with the smallest correlation and no co-linearity with annual temperature was mean diurnal range (BIO2), which presented a strong correlation with temperature annual range (BIO7 - Supplementary Fig. 2). The climatic water deficit was always chosen as the best rainfall seasonality variable and it presented a strong correlation with all other rainfall seasonality measures available, including the Precipitation of Driest Quarter (BIO17), the number of dry months (rainfall below 60 mm) and the rainfall seasonality index[76] (Supplementary Fig. 3). With few exceptions, the correlations among mean soil properties tended to be smaller than for other groups of environmental variables (Supplementary Fig. 4; see Methods section for details on soil properties).

Although we estimated several landscape metrics, the metrics with the lowest co-linearity with forest cover were landscape shape index and median fragment distance (Supplementary Fig. 5), which provide simple measures of patch aggregation in the landscape[92]. Values of the landscape shape index close to one represent high aggregation, while higher values representing disaggregation of patches. Similarly, low values of median fragment distance represent a higher aggregation of patches. Altitude, latitude, and distance from the ocean (our proxy of site continentality) explained almost all variation in annual temperature (adjusted $R^2$= 92.3%; $F$[d.o.f. =1817] = 5472), so they were not considered as candidate variables during analysis (Supplementary Fig. 6). The regression models containing all the pre-selected candidate natural and human-related variables were tested for co-linearity and all condition indices were below 4 – considered to be quite low[107]. There was some co-linearity between temperature and climatic water deficit and between forest cover and fragment size, but the correlation between these variables in the final models was acceptable (Pearson's $r \leq 0.75$).

#### Model selection

To obtain the best linear mixed-effects regression models, we used a model selection procedure based on the AIC of the candidate models to select the best structure of fixed and random effects. Differences in AIC values greater than log(8) between candidate models were regarded as an indicator of differences in model fit[108]. For the final regression model obtained for forest biomass, species richness and species properties, we tested the significance of the full model based on the comparison of a "null" model containing only random effects and sampling effort, using Chi-squared statistics (Extended Data Fig. 2). We also obtained the conditional (pseudo)$R^2$, which may be seen as the variance explained by the full model (fixed + random effects[109]). All statistical analysis was performed in R[97] using packages *lme4*[110], *MuMIn*[111], and *r2glmm*[112]. Models describing the CWM of species properties had lower explanatory powers than forest biomass and tree species richness (Supplementary Table 2), particularly for carbon-related traits. Compared with biomass and richness, species composition is more dependent on site history, random dispersal events, and species interactions (negative or positive) and thus less predictable[113].

#### Evaluation of human impacts

We obtained an index of loss related to human-induced impacts as the standardized difference between values observed in the surveys and baseline predictions for scenarios in the absence of major human impacts. The distribution of this standardized index of loss was symmetric for all variables (*i.e.*, well approximated by a normal distribution – Extended Data Fig. 3). The only exception was species richness that presented a left-skewed distribution and was better described by a Weibull distribution. We used the fit of these two distributions to estimate 95% confidence intervals around the mean index of loss, used here to assess the difference between the mean losses across our response variables.

In the main text, we used univariate models to assess whether the land conservation category (*i.e.*, conservation units, private lands, etc.) and the Human Influence Index (HII)[55] could explain the standardized index of loss for forest biomass, species richness and trait composition. Because the indices of biomass and richness loss were significantly correlated ($r^2$= 0.22), we tested if the use of multivariate regression models would improve our interpretation of the effects of land use and HII. A similar procedure was used for the indices of trait loss, which were often correlated as well (Extended Data Fig. 4–6). These models accounted for spatial autocorrelation between response variables and their parameters were estimated using Markov chain Monte Carlo (MCMC) methods[114]. The results from the bivariate models were qualitatively the same as those obtained using univariate models presented in the main text. And since the HII did not have a strong effect on the indices of loss (Supplementary Fig. 7), we finally presented only the results of the univariate model including the land-use category (Fig. 3). This model is equivalent to a standard Analysis of Variance (ANOVA), so we computed the *F*-statistics of the model and the Tukey Honest Significant Differences between the indices of loss for each pair of land-use categories. This test was performed using a 95% confidence level.

#### Effect of size and type of conservation units

We tested the shape and strength of the relationship between the indices of loss and the log-transformed size of the protected area by comparing the fit of linear, quadratic and piecewise regression models. Piecewise regression was used to detect the existence of a critical size of protected area that would minimize biomass, diversity, and trait losses. As described above, the fit of models was compared based on AIC values and only the model with the best fit to data is reported (Extended Data Fig. 7). However, we found evidence of a critical size of protected areas only for extinction level and mean trait loss.

We also compared differences in losses between conservation units classified as strict protection, on the one hand, and sustainable use of natural resources, on the other. We found differences neither in biomass and richness losses, nor for most of the traits. The two traits that presented evidence of differences were maximum height (smaller losses in lands under strict protection: adjusted $R^2$= 1.2%; $F$[d.o.f. =363] = 4.67; $p$-value= 0.031) and endemism level (smaller losses in lands under sustainable use of resources: adjusted $R^2$= 1.3%; $F$[d.o.f. =363] = 4.87; $p$-value= 0.028), but the strength of evidence was small.

#### Assessing the precision of model predictions

To assess the precision of our model predictions under the human-free scenario, we generated 5,000 simulations by sampling values of the coefficients estimated for the fixed and random effects of the model. These values were sampled from a Gaussian distribution with the estimated mean and standard errors of the coefficients. For each run, we re-calculated the model predictions but using draws of model coefficients. We then re-calculated two metrics presented in the main text, namely the standardized index of loss and the proportion of surveys with negative standardized indices of loss. This analysis was performed separately for each variable (*i.e.*, forest biomass, species richness, and species properties) using R[97] and the contributed package *merTools*[115], using the function ‘predictInterval’.

We found for all variables that the bootstrapped prediction intervals included the observed standardized indices of loss and these intervals generally did not include zero (Supplementary Table 3). The estimates of the bootstrap standard deviation were significantly higher than the observed ones for all variables (results not shown). Bias in the variance estimation is common for this type of simulation[116] and is a probable explanation for the tendency of simulated indices to be slightly more conservative than the observed ones.

For the CWM of species extinction level, the prediction interval for the index of loss included zero, meaning that the predictions for this variable were the least precise. But, the proportion of surveys with negative proportions was within the simulated intervals, meaning that the lower precision of the predictions for the index of loss of this trait is related to extreme positive values of the index. Further inspection of the index of loss for the extinction level revealed that these extreme values were commonly related to surveys with a high relative abundance of the vulnerable *Euterpe edulis* (24 to 51%) and of the endangered *Tabebuia cassinoides* (57 to 81%), two species that are red-listed due to overexploitation. For these sites, the models generated predictions 79 to 89% lower than the observed ones, generating such extreme positive indices. But those cases were rare (~15 surveys). A similar situation was found for maximum height, but in this case, the inclusion of zero within the prediction interval is probably more related to a lack of evidence for human-related impacts on this specific trait than a lack of certainty around the observed means. The high density of some species also caused some extreme values for seed mass (*e.g.*, *Araucaria angustifolia* which has a mean seed mass of 6.3 g), however, such cases had a lower impact on the predictions for this trait.

#### Reference values for undisturbed Atlantic Forests

To express in absolute terms the losses due to human-related impacts, we calculated mean values of basal area ($m^2$ $ha^{-1}$), tree richness (species $ha^{-1}$) and trait composition (CWM) to be used as references to old-growth Atlantic Forests (Supplementary Table 4). We assumed that surveys conducted in low-disturbance forests are good approximations to forests with low human impacts. These values were expected to vary among regions, so they were estimated separately for each biogeographical region of the Atlantic Forest. Reference values were estimated using a different subset of low-disturbance surveys. For forest biomass, we included surveys of a total sampling effort of ≥0.2 ha. We used the equation provided below to convert basal area into mean above-ground biomass (Mg $ha^{-1}$) and we assumed 47% of carbon concentration in the dry mass to convert above-ground biomass into above-ground carbon (Mg $ha^{-1}$). To estimate species richness, we used only surveys of ~1 ha. For trait composition and forest conservation value, averages were obtained using only surveys with 500 or more individuals sampled.

Because the dbh inclusion criterion influences the forest description[117], the reference values and consequently the average proportion of loss were calculated separately for each dbh inclusion criterion. We retrieved fewer surveys using dbh ≥3.0–3.2 cm (Extended Data Table 1), and even fewer surveys of low-disturbance forests with total sampling effort of ≥0.2 ha or ~1 ha (Supplementary Table 4). Therefore, we conducted no projections of human-induced impacts based on forests surveyed using dbh ≥3.0–3.2 cm. For species traits, ecological groups and conservation value, dbh ≥5.0 was the most frequent cutoff criterion. Surveys using dbh ≥5.0 cm represent better the composition of the lower tree layer of the forest, which contains the individuals that will compose the future canopy of the forest. Considering the long lifecycle of tree species, we assume that trees between dbh 5.0 and 10.0 cm should reflect better the changes in forest composition. So, the reference values and average losses of species properties are reported only for dbh ≥5.0 cm (Extended Data Fig. 8, Supplementary Table 4).

#### Projections across the Atlantic Forest

In the main text, we present the projections of forest carbon loss for the entire Atlantic Forest. We extended the average region-specific losses of carbon for the remaining Atlantic Forest fragments for which no data was available. We assumed the same average carbon loss for each remaining fragment at each biogeographic region and we used the remaining forest are in each biogeographical region to project carbon losses. Although the average proportion of species loss and changes in species properties were obtained, the projection of these proportions for the entire Atlantic Forest is more complex. Observed species richness is non-linearly related to the sampling area[117], so it cannot be directly extrapolated from smaller sample sizes to the entire Atlantic Forest area. Moreover, the loss of species from a local community does not necessarily mean that these species are locally or regionally extinct. Thus, the equivalent forest area lost due to losses in species richness cannot be calculated. Also, there is no market for "biodiversity credits", so pricing species losses is very difficult. Trait composition is contingent on community species composition, which is highly variable across the Atlantic Forest hotspot. This also hinders the possibility of projecting and valuing trait composition losses across the entire study area.

#### Implications for the Atlantic Forest restoration

As explained in the main text, the fragment and landscape restoration scenarios were simulated for the Brazilian Atlantic Forest using the 289 thousand fragments remaining in 2016 mapped by ref. 53. In practice, we used our models to predict values of biomass, species richness and CWM of species characteristics for each fragment. These predictions were used to set the ‘current situation’ of the Atlantic Forest fragments and they were obtained by using the climatic, topographic and biogeographic variables available for the central coordinate of each fragment. Fragment size is provided by ref. 53. For the model co-variables not spatially available or not available in the same spatial resolution used for analysis (*i.e.*, soil type, forest disturbance level, landscape metrics), we used our 1,819 surveys to assign average values for each biogeographical region of the Atlantic Forest[54] (Extended Data Fig. 1). All predictions were generated for the same sampling effort (*i.e.*, one hectare), sampling design (*i.e.*, systematic plots dbh ≥5 cm) and using the average number of trees per hectare for each region (Supplementary Table 5).

To simulate the two restoration scenarios, we used the same co-variable values used to predict the ‘current situation’ of biomass, species richness and CWMs with exception to the fragment disturbance level and patch/landscape metrics. Because values of fragment disturbance level and landscape metrics were region-specific, the changes in disturbance level and landscape metrics between the current and restoration predictions also were region-specific. For instance, the decrease in disturbance level and the increase in patch/landscape metrics were smaller for fragments in the Serra do Mar region (lower fragmentation) than in the Alto Paraná region (highest fragmentation, Supplementary Table 5). While generating the predictions for the ‘landscape restoration’ scenario around forest fragments ≥5,000 ha, we assumed no increase in fragment size and only half of the increase of forest cover.

The changes in patch/landscape metrics necessary to meet the 20% forest cover target were obtained from 4×4 km simulated landscapes. These landscapes were constructed using averages of patch density and size distribution of forest patches for each biogeographical region. The distribution of patch size for each simulation was obtained using the region-specific lognormal distributions fitted to the 2016 Atlantic Forest fragments. For each iteration, we calculated the landscape metrics for the current landscape, and if the landscape forest cover was smaller than 20%, we obtained the amount of forest cover necessary to reach the 20% target. This ‘to-be-restored’ forest area was distributed among the forest patches proportionally to their original sizes, simulating restoration efforts around the existing fragments. We re-calculated the landscape metrics for this ‘restored landscape’ and then calculate the changes in these metrics between the ‘current’ and ‘restored’ landscapes. For each biogeographical region, we average these changes for 100 iterations, generated using contributed R package *landscapeR*[118]. As above, landscape metrics were obtained using the R package *SDMTools*[93].

#### The tree species list for the Atlantic Forest restoration

Aiming to assist the selection of species that could maximize restoration efforts (*e.g.*, enriching forest fragments), we provide a list of frequent Atlantic Forest species with higher-than-average potential for carbon storage (*i.e.*, high wood density and maximum adult height), ecological interactions (*i.e.*, large seed mass and late-successional species) and biodiversity conservation (*i.e.*, threat and endemic status). To produce this list, we first selected the species within the 100 most frequent in each of the eight biogeographical regions used in this study (Extended Data Fig. 1). We then ranked species regarding their wood density, maximum adult height, seed mass, ecological group, IUCN threat category, and endemism level. Once again, the three last species categories were treated as ordinal categorical data. We standardized each rank and then average them to obtain a mean rank for each species. In the final list we included only species with a mean rank higher than the average for all species (frequent or not frequent). We also present the average rank for each group of species properties (*i.e.*, carbon storage potential, ecological interactions, and conservation status) and the frequency of each species in each biogeographical region, as percentages (Supplementary Data 2). Therefore, this list represents a compromise between species that are easier to find for seedling production and have the greatest potential to improve ecosystem provision, taxonomic and functional diversity and/or conservation value of the Atlantic Forest. We removed *Dicksonia sellowiana* from the list because the seedling production from spores for this threatened tree-fern is incipient.

#### Basal area as a measure of forest biomass

Forest above ground biomass (AGB; live oven-dry matter of trees, in Mg $ha^{-1}$) is a key determinant of ecosystem integrity. However, it is more difficult to retrieve estimates of AGB than basal area (BA; summed cross-section area of the tree trunks, in $m^2$ $ha^{-1}$) from the literature. Estimates of AGB[77] in forest plots are expected to be closely related to BA because both depend on tree diameter. Indeed, strong statistical relationship between BA and AGB has been found in Mexican forests at stand level and in the Atlantic Forest at species level[25]. But the strength of such a relationship at stand level has never been tested for the Atlantic Forest.

We compiled 503 surveys for which both AGB and BA were available from two Brazilian state inventories[119–122]. AGB estimates represent different Atlantic Forest formations, using allometric equations specific to each forest type, based on tree diameter and height (Supplementary Table 6). We fitted simple linear regression models to the log-transformed values of AGB and BA for each set of data separately. These models were compared against non-linear models (*i.e.*, power and exponential functions) to assess the assumption of linearity in this relationship. We also tested the impact of survey sampling area in the relationship between the two variables by adding it as weights in the regression model and inspecting if there was any improvement on model fit.

We found a strong, linear and positive relationship of the log-transformed values of AGB and BA for each of the main Atlantic Forest types. Basal area explained 89–97% of the variation in AGB (Supplementary Fig. 9). For Seasonal forests (Supplementary Fig. 9, panel C), a power-function fitted values slightly better (results not shown) due to a better fit of very extreme values of BA (below 8 and above 40 $m^2$ $ha^{-1}$ of BA). But the linear function was a very good description of the relationship for all forest types.

We also fitted linear regressions for forest types altogether. The linear mixed effect regression model assuming different random intercepts and slopes for each forest type (conditional $R^2$= 96.5%) yielded a much better fit than the model considering only BA as a predictor of AGB (adjusted-$R^2$= 88.2%). So, the general expression averaging parameter estimates across forest types is: AGB = exp[1.129 + 1.178 *ln*(BA)]. The 95% confident interval estimates of the regression parameters were 0.598–1.688 for the intercept and 1.031–1.327 for the slope parameter. We validated this equation with another set of 53 unpublished Atlantic Forest surveys using the global allometric equations provided by ref. 123, and found an excellent match, demonstrating that AGB can be confidently inferred from BA. We converted aboveground biomass (Mg $ha^{-1}$) into carbon storage (Mg C $ha^{-1}$) by assuming 47% of carbon concentration[124].

#### 2. Supplementary Results and Discussion

##### Evaluation of human impacts

The majority of the analyzed variables (species richness, species properties and forest biomass) had mean standardized indices of loss that were significantly negative (Extended Data Fig. 3). Moreover, the frequency of sites presenting negative indices of loss was between 60–88% (Supplementary Table 3). These results suggest that post-deforestation, human-induced impacts are pervasive across the Atlantic Forest. About 17% of the sites presented positive indices of loss (*i.e.*, gains) of species richness or forest biomass and 5% presented gains in richness and biomass simultaneously (Fig. 1). Increases in species diversity in disturbed forests can occur depending on the frequency and intensity of the disturbances, by a balance of the contribution of early successional and of late sucessional species[125,126]. Increases in carbon storage potential can also follow the increase in the availability of resources due to global changes, such as atmospheric $CO_2$, solar radiation and rainfall[127]; although these gains are small compared to losses caused by disturbances[128]. Even though the goodness of fit measures of our models were fairly high for species richness and forest biomass (pseudo-$R^2$= 71 and 53%, respectively), model fits were not perfect and they will not generate accurate predictions for all surveys. If a given survey was conducted in a forest under a combination of climate, soil, and human-related conditions which is underrepresented in our data set, predictions in the human-free scenario may be smaller than the observed values. Thus, caution must be taken when interpreting the index of loss for individual surveys.

As mentioned in the Supplementary Methods, the Human Influence Index (HII)[55] around the surveyed fragments had small or no effects on the magnitude of human-related impacts (Supplementary Fig. 7). Exchanging the HII with the distance from main cities (>100,000 inhabitants), did not increase the explanatory power of this analysis (not shown). This is unexpected since the HII combines different vectors of human pressure and accessibility, which should be an important vector of human-induced impacts in forest fragments. This result may be explained by different factors: (*i*) the highly fragmented nature of the Atlantic Forest: there are few fragments that are very large, undisturbed and really "far from" human presence, (*ii*) and the nature of the human impact itself: disturbed fragments are the combination of different and idiosyncratic histories of land use changes and degradation. For instance, urban forests can be in better conditions than suburban ones because they represent a vital resource for society (both aesthetic and monetary). On the other hand, small and isolated fragments in private lands can be well-conserved and diversified, depending on the understandings of the land owner regarding the natural environment or whether the surrounding community have access and can use the forest resources. Other authors found that the effects of human-impacts in tropical forests can be quite unpredictable at regional scales[129–131]. Therefore, human impacts in the Atlantic Forest are the result of a complex history which makes current patterns relatively independent from human presence indices. So, although we found differences in average losses across biogeographical regions (Extended Data Fig. 8), biomass and biodiversity losses remained difficult to predict from human pressure/presence indices.

#### Implications for the Atlantic Forest restoration

As mentioned in the main text, we used our regression models to predict the costs and gains of two different restoration scenarios: one that focused only on reducing the within-fragment forest disturbance level (*i.e.*, 'fragment restoration' scenario) and the other the focused on increasing fragment size, landscape integrity and connectivity (see 'Implications for restoration' in Supplementary Methods).

In addition to the results already presented in the main text, we found a tendency for gains to be proportionally higher in the Alto Paraná and Northeast forests than in other biogeographical regions. These areas are within the more fragmented and disturbed regions of the Atlantic Forest, meaning that restoration efforts in these regions would provide better outcomes regarding carbon and biodiversity recovery within existing fragments. Consequently, these regions also presented fragment restoration yields that were comparatively higher when compared to landscape restoration yields (Supplementary Table 5). Thus, in these regions the fragment restoration strategy is more cost-effective than in other Atlantic Forest regions, adding additional knowledge on how to prioritize restoration efforts in the Atlantic Forest[64]. The Atlantic Dry region was also among the regions with the highest proportional carbon gains in the fragment restoration scenario, but in this case due to its higher forest cover and smaller carbon storage potential (less restored area combined with less carbon gain per hectare to attain the 20% forest cover target of the landscape restoration scenario). In contrast, in the Serra do Mar and Bahia Coastal regions, which combine higher carbon storage potentials with lower disturbance levels, the expected outcomes of restoration within fragments were among the lowest when compared to restoration of the surrounding landscape. On average, these areas have higher landscape resilience and thus lower implementation costs of restoration, making landscape restoration proportionally more efficient.

This is a simple exercise of the possible outcomes of contrasting restoration strategies since it assumes that all fragments/landscapes will have high restoration success and will follow the same restoration trajectories[132]. A more complete appraisal would require spatialized, fragment-specific information on the disturbance level, landscape conditions, restoration success/unpredictability, and land opportunity costs, as well as a more complete and detailed assessment of the impact of the different restoration scenarios and restoration targets on species survival and on ecosystem services other than carbon storage (*e.g.*, refs. 102, 134, 138). However, these two simple scenarios provided an approximation of costs and how outcomes may vary across the Atlantic Forest. In addition, they allowed us to indirectly explore the role of within-fragment disturbances and patch/landscape metrics as drivers of biomass and biodiversity loss.

#### 3. Supplementary Tables and Figures

**Supplementary Table 1. Summary of the Atlantic Forest surveys studied.**

Description of the tree community surveys used to estimate human-related impacts on forest fragments, separated by diameter at breast height (dbh) inclusion criteria. Number of species per dbh inclusion criterion refers only to the tree community surveys included in the analyses of multiple species properties (*n*= 1213).

| Dbh inclusion criteria (cm) | Number of surveys | Total effort (ha) | Number of trees included | Number of tree species |
|---|---|---|---|---|
| ≥3.0-3.2 | 130 | 35.9 | 87,019 | 907 |
| ≥4.8-5.0 | 1,063 | 703.5 | 1,059,410 | 2,987 |
| ≥10.0 | 626 | 498.4 | 301,933 | 1,291 |
| All criteria | 1,819 | 1,237.8 | 1,448,362 | 3,124 |

**Supplementary Table 2. Summary of the environmental and human-related conditions of the forest fragments studied.**

Mean, standard deviation (s.d.), minimum (Min.), maximum (Max.), first and third quartiles, and the coefficient of variation (%) of t continuous descriptors of the surveyed sites (*n*= 1819). Temperature, rainfall, climatic water deficit, mean frost, and potential direct incident radiation are annual means, while soil quality and mean shape index are dimensionless. For climatic water deficit, mean frost frequency, slope declivity, fragment size and core forest cover, all with skewed distributions, values are actually medians and the coefficient of variation was calculated for the transformed variables, as used in the data analysis.

| **Variable** | **Mean ± s.d.** | **Min.-Max.** | **1st - 3rd quartiles** | **Coefficient of variation** |
|---|---|---|---|---|
| Annual temperature ($^{o}$C) | 19.3 ± 2.8 | 11.3−25.7 | 17.2−21.3 | 14.5 |
| Diurnal range ($^{o}$C) | 10.4 ± 1.65 | 5.7−14.5 | 9.5−11.7 | 15.8 |
| Frost frequency (days) | 0.020 ± 0.18 | 0−1.032 | 0−0.126 | 71.9 |
| Annual rainfall (mm) | 1637 ± 301 | 501−3062 | 1406−1811 | 18.4 |
| Climatic water deficit (mm) | -136 ± 173 | -1091−0 | -256−0 | 95.9 |
| Relative humidity (%) | 79.4 ± 3.3 | 63.8−84.1 | 77.1−81.0 | 34.5 |
| Cloud cover (%) | 69.2 ± 5.4 | 47.3−78.9 | 65.5−73.8 | 7.8 |
| Direct radiation (MJ cm$^{-2}$ yr$^{-1}$) | 0.97 ± 0.12 | 0.44−1.17 | 0.91−1.05 | 12.2 |
| Slope declivity ($^{o}$) | 10.0 ± 8.9 | 0−47 | 4−18 | 35.4 |
| Soil quality | 6.99 ± 2.19 | 1.9−14.5 | 6−8 | 31.7 |
| Cation retention (cmol$_c$ kg$^{-1}$) | 14.6 ± 14.2 | 2.9−81.0 | 9.6−22.6 | 20.4 |
| Fragment size (ha) | 155 ± 42473 | 0.2−303,000 | 29−1784 | 53.5 |
| Forest cover (%) | 47.6 ± 29.5 | 0.04−100 | 23.1−71.7 | 62.0 |
| Core forest cover (%) | 13.9 ± 30.2 | 0−100 | 3.87−46.6 | 43.1 |
| Mean shape index | 1.56 ± 0.29 | 1−4.41 | 1.42−1.63 | 18.8 |
| Med. fragment distance (m) | 1889 ± 736 | 0−3047 | 1364−2121 | 45.2 |

**Supplementary Table 3. Description of the linear mixed-effects regression models with the best fit to forest biomass, species richness, and species properties data.**

Each model is the result of a model selection procedure of different random and fixed effects structures. Candidate fixed effects included different environmental and human explanatory variables (and their interactions), while random effects included the survey methodology and the Atlantic Forest biogeographical regions. For each model, we present the transformation applied to each variable, the number of observation (*n*), marginal/conditional $R^2$, full model Chi-squared statistics and number of model parameters (Par.), along with the selected fixed effects and interactions (separated by a vertical slash). Full variables names are given in Suppl. Table 1 (CWD= Climatic water deficit).

| **Variable** | **Transf.** | ***n*** | **$R^2$ (%)** | **Chi$^2$** | **Par.** | **Fixed effects** | **Interactions** |
|---|---|---|---|---|---|---|---|
| Forest biomass | log | 1676 | 38.5/52.7 | 362.9 | 16 | Temperature\|Slope\|Soil fertility\|For. disturbance\|Core for. cover\|Tree density\|Effort | Temperature:Slope\|For. disturbance:Tree density\|For. disturbance:Core forest cover |
| Species richness | log | 1790 | 58.2/70.7 | 390.2 | 20 | Temperature\|Diurnal range\|Slope\|Env. stress\|Soil fertility\|Organic matter\|For. disturbance\|Fragment size\|Forest cover\|Shape index\|log(N) | Env. stress:Soil fertility |
| Wood density | log | 1213 | 18.9/25.8 | 126.6 | 15 | Temp.seasonality\|Slope\|Cloud cover\|Soil quality\|Radiation\|For. disturbance\|Forest cover\|Fragment size\|Frag. distance\|Effort | Radiation:For. disturbance |
| Adult height | – | 1213 | 9.4/27.0 | 82.2 | 14 | Temperature\|Diurnal range\|CWD\|Cloud cover\|Soil fertility\|Slope\|Fragment size\|Forest cover\|Effort | Fragment size:Forest cover |
| Seed mass | log | 1214 | 28.3/35.2 | 203.2 | 14 | Temperature\|Isothermality\|Radiation\|CWD\|Humidity\|Soil quality\|For. disturbance\|Core for. cover\|Effort | Humidity:Soil quality |
| Ecological group | Ridit | 1214 | 23.5/35.7 | 211.2 | 15 | Frost\|Diurnal range\|Humidity\|Soil fertility\|For. disturbance\|Core for. cover\|Tree density\|Effort | Frost:Humidity\|Humidity:Soil fertility\|For.disturbance:Core for. cover |
| Extinction level | Ridit | 1214 | 26.2/41.6 | 186.3 | 13 | Frost\|Diurnal range\|CWD\|Cloud cover\|Organic matter\|For. disturbance\|Core for. cover\|Effort | Frost:Seasonality |
| Endemism level | Ridit | 1213 | 37.1/43.6 | 290.9 | 16 | Temperature\|Slope\|Humidity\|Cloud cover\|Soil quality\|For. disturbance\|Fragment size\|Shape index\|Effort | Temperature:Humidity\|Humidity:Soil quality\|For. disturbance:Fragment size |

**Supplementary Table 4. Precision of the model predictions in the human-free scenario.**

For each variable included in our study we present the observed, the bootstrap estimates and 95% confidence intervals (inside brackets) of the average standardized index of loss and the frequency of Atlantic Forest surveys presenting losses due to human-related impacts (*i.e.*, negative standardized index of loss). The bootstrap means and intervals of the predictions were generated using 5000 samples of the model coefficients taken from normal distributions around the parameters estimated by the models fitted to the data.

| **Variable** | **Standardized Index of loss** | | **Frequency (%)** | |
|---|---|---|---|---|
| | Observed | Simulated [CI 95%] | Observed | Simulated [CI 95%] |
| Forest biomass | -0.089 | -0.082 [-0.115; -0.047] | 83.5 | 74.9 [65.3; 83.4] |
| Species richness | -0.072 | -0.065 [-0.103; -0.022] | 82.7 | 71.0 [57.7; 82.3] |
| Wood density | -0.066 | -0.099 [-0.163; -0.042] | 59.9 | 61.3 [52.7; 69.5] |
| Adult height | 0.037 | 0.050 [-0.003; 0.106] | 38.0 | 40.5 [28.3; 53.9] |
| Seed mass | -0.101 | -0.079 [-0.126; -0.028] | 75.8 | 68.0 [59.1; 76.3] |
| Ecological group | -0.182 | -0.152 [-0.213; -0.085] | 84.9 | 74.6 [65.7; 82.3] |
| Extinction level | -0.098 | -0.022 [-0.172; 0.248] | 66.9 | 59.5 [48.4; 69.9] |
| Endemism level | -0.267 | -0.251 [-0.321; -0.075] | 86.9 | 79.5 [71.1; 86.6] |

**Supplementary Table 5. Mean reference values of forest biomass, tree species richness and species properties for the Atlantic Forest.**

Average estimates were obtained using only surveys conducted in low-disturbance forest fragments. For biomass, we considered only surveys with total sampling area >0.2 ha. For species richness, we considered only surveys of about one hectare of total effort. References for the community-weighted means for species properties were obtained only for surveys using dbh ≥5 cm and with a minimum sample size of 500 individuals. Values of average carbon were estimated based on the relationship between above ground biomass and basal area for Atlantic Forest sites (see 'Supplementary Methods') and assuming 47% of carbon content in dry biomass. Mean references and confidence intervals were estimated separately for each biogeographical region, but here we present the weighted average of these estimates across these regions, using the region areas as weights.

| **Forest descriptor** | **Dbh cutoff (cm)** | **Number of surveys** | **Mean reference** | **Confidence interval (95%)** |
|---|---|---|---|---|
| Basal area ($m^2$ $ha^{-1}$) | ≥5.0 | 239 | 33.6 | 29.3−37.9 |
| | ≥10.0 | 82 | 30.9 | 22.8−39.1 |
| Carbon storage (Mg $ha^{-1}$) | ≥5.0 | 239 | 93.0 | 79.3−107.2 |
| | ≥10.0 | 82 | 84.3 | 59.5−111.3 |
| Species richness ($ha^{-1}$) | ≥5.0 | 121 | 104 | 84−130 |
| | ≥10.0 | 37 | 75 | 51−115 |
| Species properties | | | | |
| Wood density (g $cm^{-3}$) | ≥5.0 | 326 | 0.632 | 0.617–0.645 |
| Max. adult height (m) | ≥5.0 | 326 | 22.5 | 21.7–23.3 |
| Seed mass (g) | ≥5.0 | 326 | 0.172 | 0.132–0.198 |
| Ecological group | ≥5.0 | 326 | 0.912 | 0.848–0.975 |
| Extinction level | ≥5.0 | 326 | 0.214 | 0.158–0.306 |
| Endemism level | ≥5.0 | 326 | 0.646 | 0.575–0.712 |

**Supplementary Table 6. Reference values used to simulate the two restoration scenarios, 'fragment restoration' (Scenario 1) and 'landscape restoration' (Scenario 2), and the description of the gains and costs considered to calculate their outcomes.**

For each biogeographical region of the Atlantic Forests, we provide the average values used to predicted the differences between the 'current' and the two restoration scenarios. For Scenario 1, there is no increase in forest cover (FC). For Scenario 2, the increase in FC represents the amount of forest cover that should be restored to reach 20% of average landscape forest cover. Costs of restoration for the Scenarios 1 and 2 vary according to the average disturbance level of the fragments and the average forest cover, respectively.

| | **Alto Paraná** | **Araucaria forests** | **Atlantic Dry** | **Bahia Coast** | **Bahia Interior** | **Northeast forests** | **Serra do Mar** | **Uruguay forests** |
|---|---|---|---|---|---|---|---|---|
| Tree density (dbh≥ 5 cm $ha^{-1}$) | 1400 | 1718 | 891 | 1533 | 1505 | 1263.8 | 1663.8 | 1681.1 |
| Ref. carbon storage (Mg C $ha^{-1}$) | 80.1 | 115.6 | 45.2 | 89.1 | 67.6 | 77.4 | 96.4 | 90 |
| Fragment disturbance level[a] | -0.19 | -0.06 | 0.16 | 0.16 | -0.14 | -0.4 | 0.22 | 0.11 |
| Average forest cover (FC) (%) | 8.6 | 17.8 | 23.6 | 14.4 | 11.2 | 17.0 | 41.6 | 16.5 |
| Average forest size (ha) | 52 | 67 | 217 | 51 | 43 | 61 | 163 | 68 |
| Mean patch density (16 $km^{-2}$) | 5.7 | 6.8 | 4.8 | 5.5 | 7.5 | 7.1 | 5.2 | 5.3 |
| Med. fragment distance (km) | 1.79 | 1.78 | 2.00 | 1.47 | 2.00 | 2.05 | 1.23 | 1.73 |
| Remaining FC (million ha) | 3.46 | 3.84 | 2.81 | 1.63 | 2.77 | 1.02 | 4.69 | 0.93 |
| FC increase (Scenario 2; million ha) | 1.80 | 1.54 | 1.43 | 1.88 | 1.75 | 1.34 | 1.17 | 1.86 |
| Carbon Gain Scenario 1 (Mg $ha^{-1}$) | 5.1 | 5.5 | 2.6 | 3.5 | 4.2 | 6.5 | 3.3 | 3.7 |
| Carbon Gain Scenario 2 (Mg $ha^{-1}$) | 59.7 | 95.1 | 33.6 | 63.5 | 48.7 | 48.0 | 74.1 | 69.6 |
| Costs Scenario 1 (US$ $ha^{-1}$)[b] | 876 | 783 | 618 | 617 | 841 | 1034 | 573 | 658 |
| Costs Scenario 2 (US$ $ha^{-1}$)[c] | 2654 | 2516 | 2430 | 2568 | 2615 | 2529 | 2154 | 2535 |
| Yield Scen. 1 (Mg C $ha^{-1}$ per million US$)[d] | 5,838 | 7,048 | 4,269 | 5,631 | 4,935 | 6,234 | 5,752 | 5,615 |
| Yield Scen. 2 (Mg C $ha^{-1}$ per million US$)[d] | 22,508 | 37,797 | 13,812 | 24,735 | 18,627 | 18,966 | 34,409 | 27,454 |

[a] Average disturbance level of fragments are scaled around zero and vary from high (negative values) to low (positive values).

[b] Costs for scenario 1 ('fragment restoration') vary around 750 US$ $ha^{-1}$ according to the average disturbance level of biogeographical regions.

[c] Costs for scenario 2 ('landscape restoration') vary around 2500 US$ $ha^{-1}$ according to the average forest cover of biogeographical regions.

[d] Yield is calculated by the amount of carbon increase in each scenario (Mg C $ha^{-1}$) divided by the total restoration costs (million US$).

**Supplementary Table 7. Studies used to evaluate the relationship between above ground biomass and basal area in the Atlantic Forest.**

For each study containing surveys with estimates of above ground biomass and basal area simultaneously, we present the Atlantic Forest formation studied, the number of surveys used in the analysis, their combined sampled area and dbh inclusion criteria.

| Source | Forest formation | Number of surveys | Total effort (ha) | Dbh inclusion criteria (cm) |
|---|---|---|---|---|
| Ref. 120 | Seasonal forests | 78 | 28.7 | ≥10 |
| Ref. 121 | *Araucaria* forests | 151 | 59.1 | ≥10 |
| Ref. 122 | Rainforests | 201 | 72.1 | ≥10 |
| Ref. 119 | Seasonal and Rainforests | 73 | 109.6 | ≥5 |

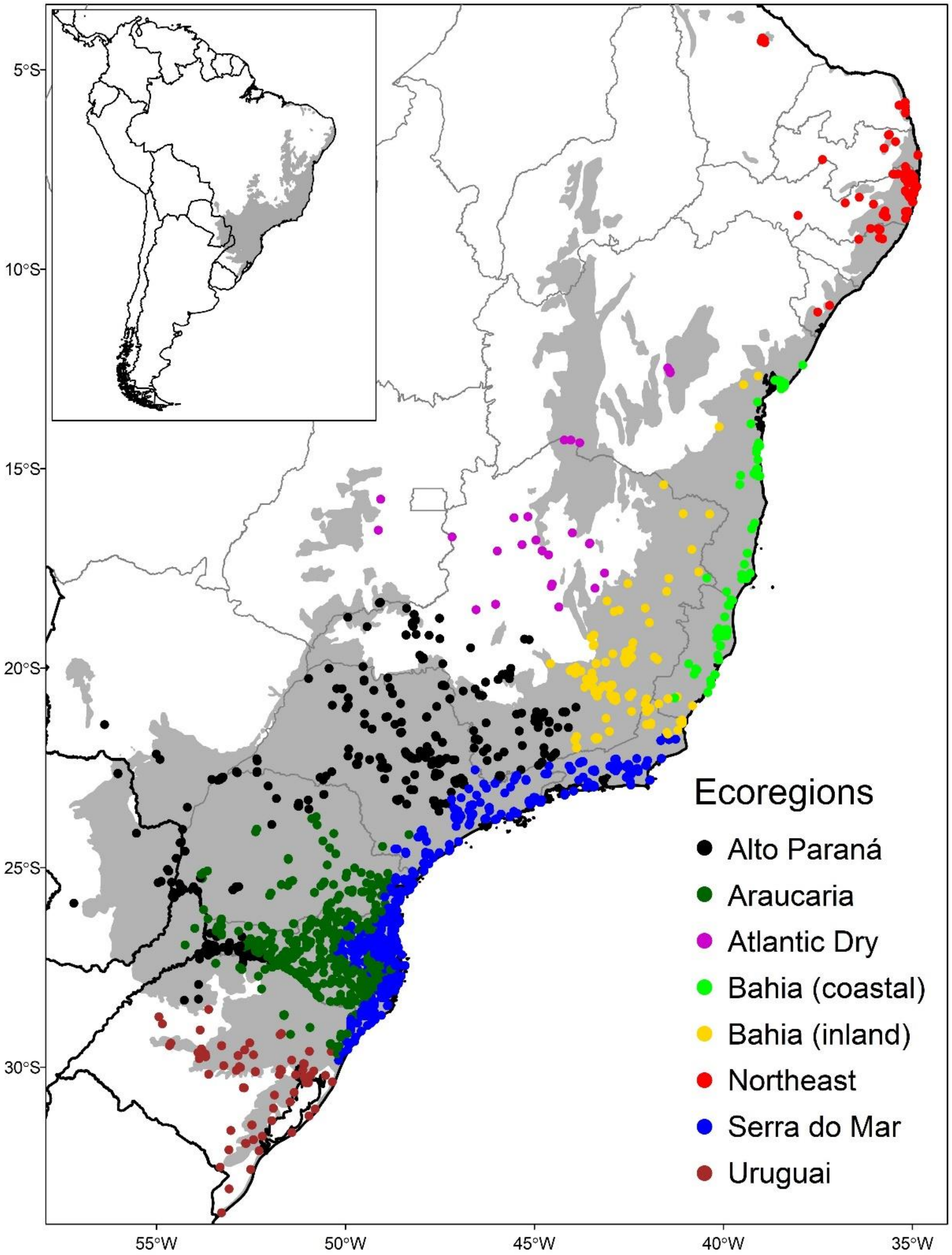

**Supplementary Figure 1. The Atlantic Forest and the distribution of the tree community surveys used in this study.**

The shaded grey corresponds to the Atlantic Forest limits and each circle represents one forest survey. Colours of circles correspond to the eight eco-regions used as a biogeographical control during data analyses. We considered surveys of forests classified by the original authors as Atlantic Forests, even if they were outside the Atlantic Forest limits.

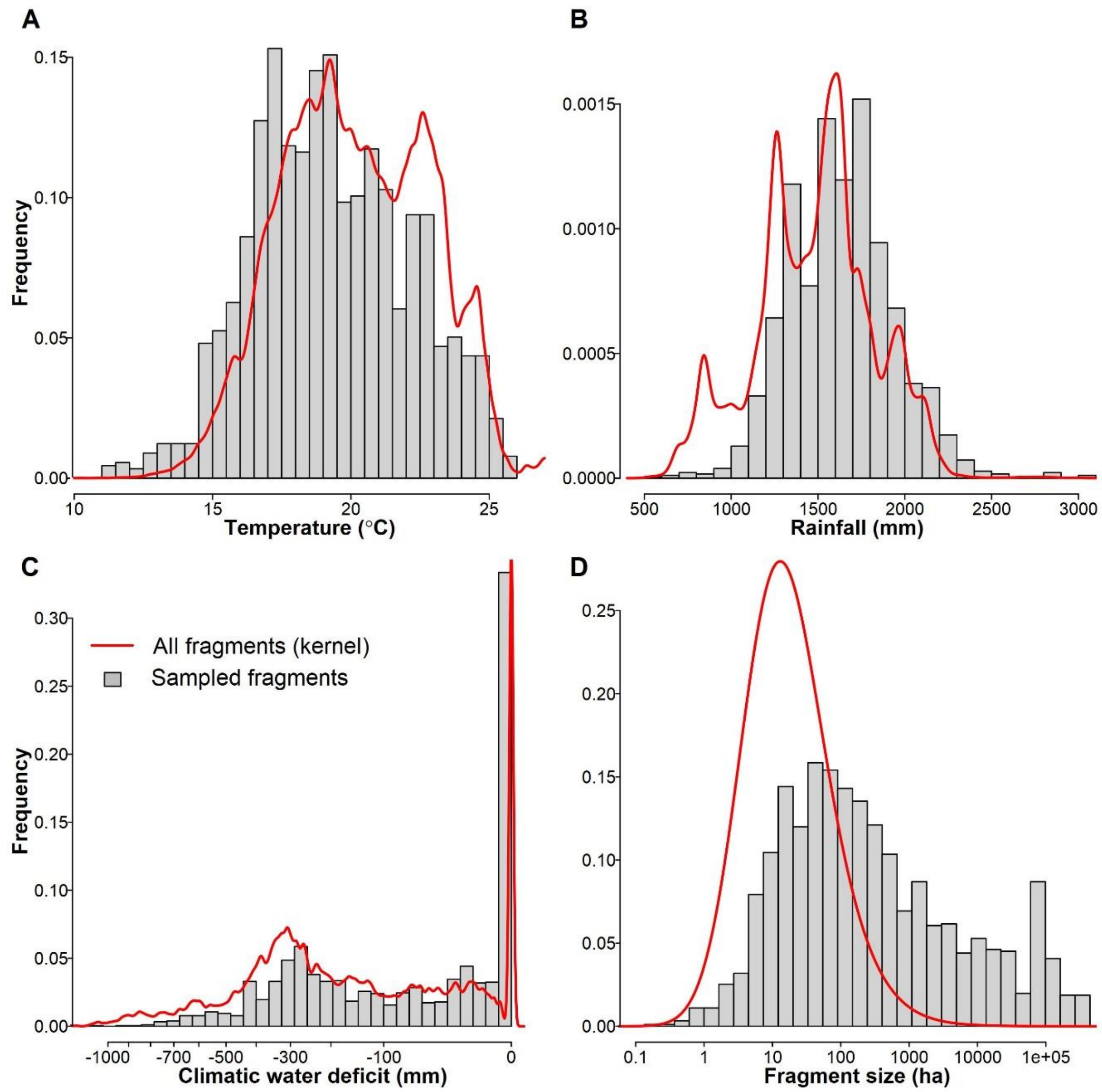

**Supplementary Figure 2. Comparison of climate and fragment size between all fragments in the Atlantic Forest and the fragments included in our sample.**

Panels represent the distribution of conditions in the surveys (grey; n=1,819) compared with a sample of fragments across the Atlantic Forest (red line; n=250,000) for

Panels represent (**A**) the distribution of mean annual air temperature; (**B**) mean annual rainfall; (**C**) climatic water deficit; and (**D**) forest fragment size for all Atlantic Forest fragments (red thick lines; n= ~250,000). These are compared with their distribution in the fragments included in the analysis (grey bars; n= 1,819). Note that the *x*-axes for panels C and D are transformed.

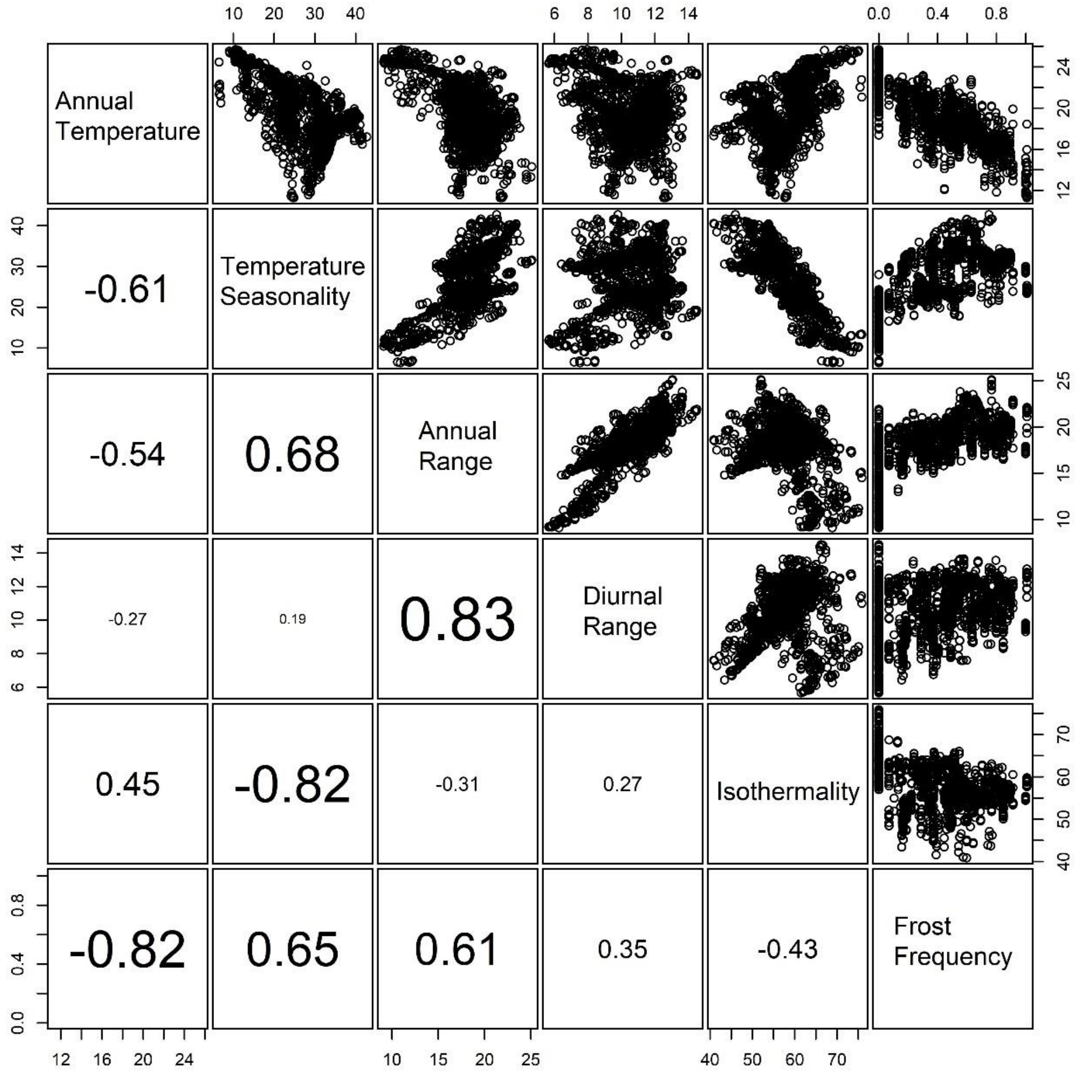

**Supplementary Figure 3. Correlations between pairs of candidate explanatory variables related to air temperature and its seasonality.**

The six temperature-related candidate explanatory variables (legends in the diagonal) that were considered during the construction of the linear mixed-effect models (see 'Methods' for definition and sources of each variable). Above the diagonal, the scatter plots between each pair of variables (each point represents a forest survey) are presented. The value of Pearson's correlation index for the corresponding pair of variables is given below the diagonal. The size of the font of the correlation index is proportional to the strength of the correlation.

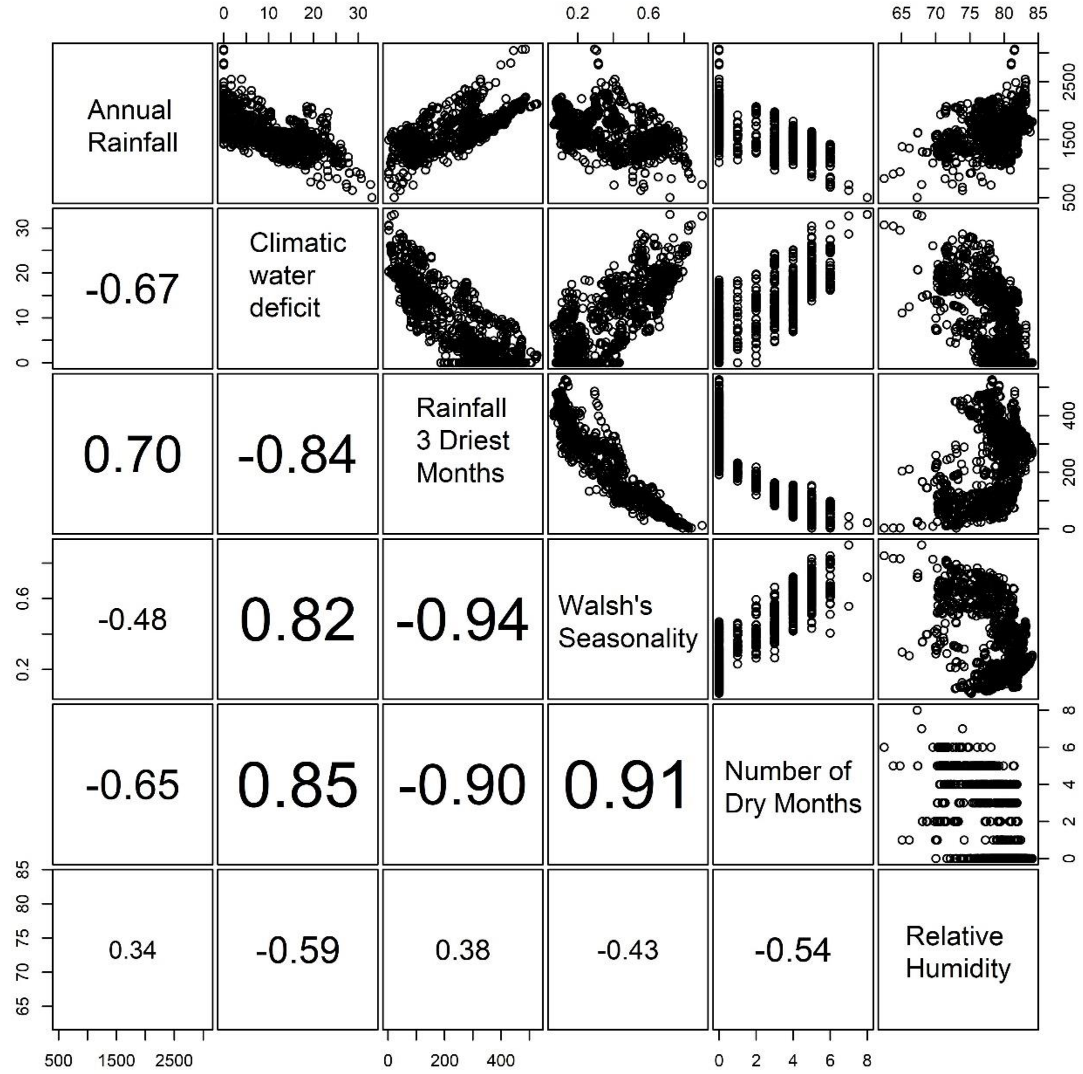

**Supplementary Figure 4. Correlations between pairs of candidate explanatory variables related to rainfall and its variation in time.**

The six rainfall-related candidate explanatory variables (legends in the diagonal) that were considered during the construction of the linear mixed-effect models (see 'Methods' for definition and sources of each variable). Above the diagonal, the scatter plots between each pair of variables (each point represents a forest survey) are presented. The value of Pearson's correlation index for the corresponding pair of variables is given below the diagonal. The size of the font of the correlation index is proportional to the strength of the correlation.

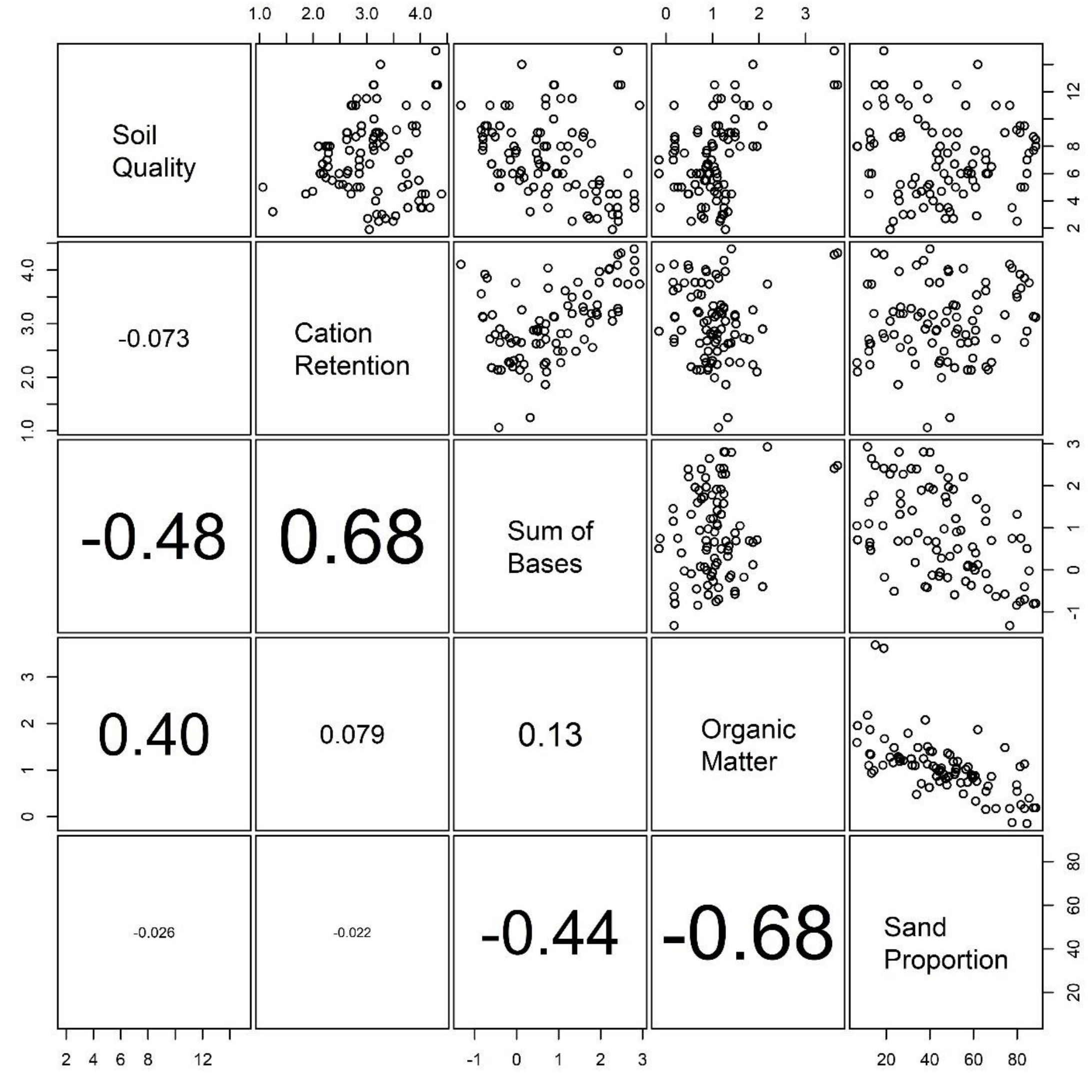

**Supplementary Figure 5. Correlations between pairs of candidate explanatory variables related to soil properties.**

The five soil-related candidate explanatory variables (legends in the diagonal) that were considered during the construction of the linear mixed-effect models (see 'Methods' for definition and sources of each variable). Above the diagonal, the scatter plots between each pair of variables (each point represents a forest survey) are presented. The value of Pearson's correlation index for the corresponding pair of variables is given below the diagonal. The size of the font of the correlation index is proportional to the strength of the correlation.

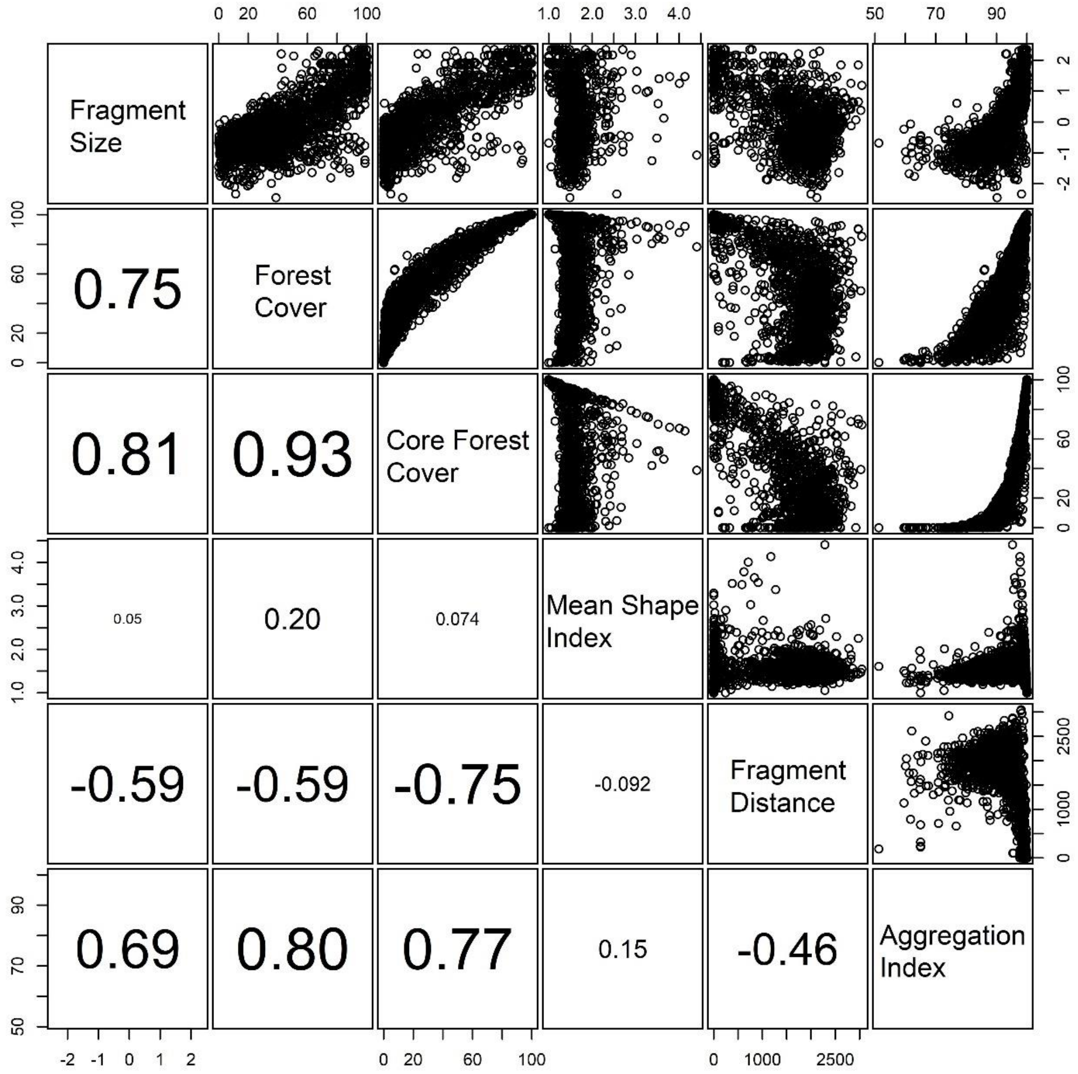

**Supplementary Figure 6. Correlations between pairs of candidate explanatory variables related to fragment and landscape metrics.**

The six patch and landscape metrics used as candidate explanatory variables (legends in the diagonal) during the construction of the linear mixed-effect models (see 'Methods' for definition and sources of each variable). Above the diagonal, the scatter plots between each pair of variables (each point represents a survey) are presented. The value of Pearson's correlation index for the corresponding pair of variables is given below the diagonal. The size of the font of the correlation index is proportional to the strength of the correlation.

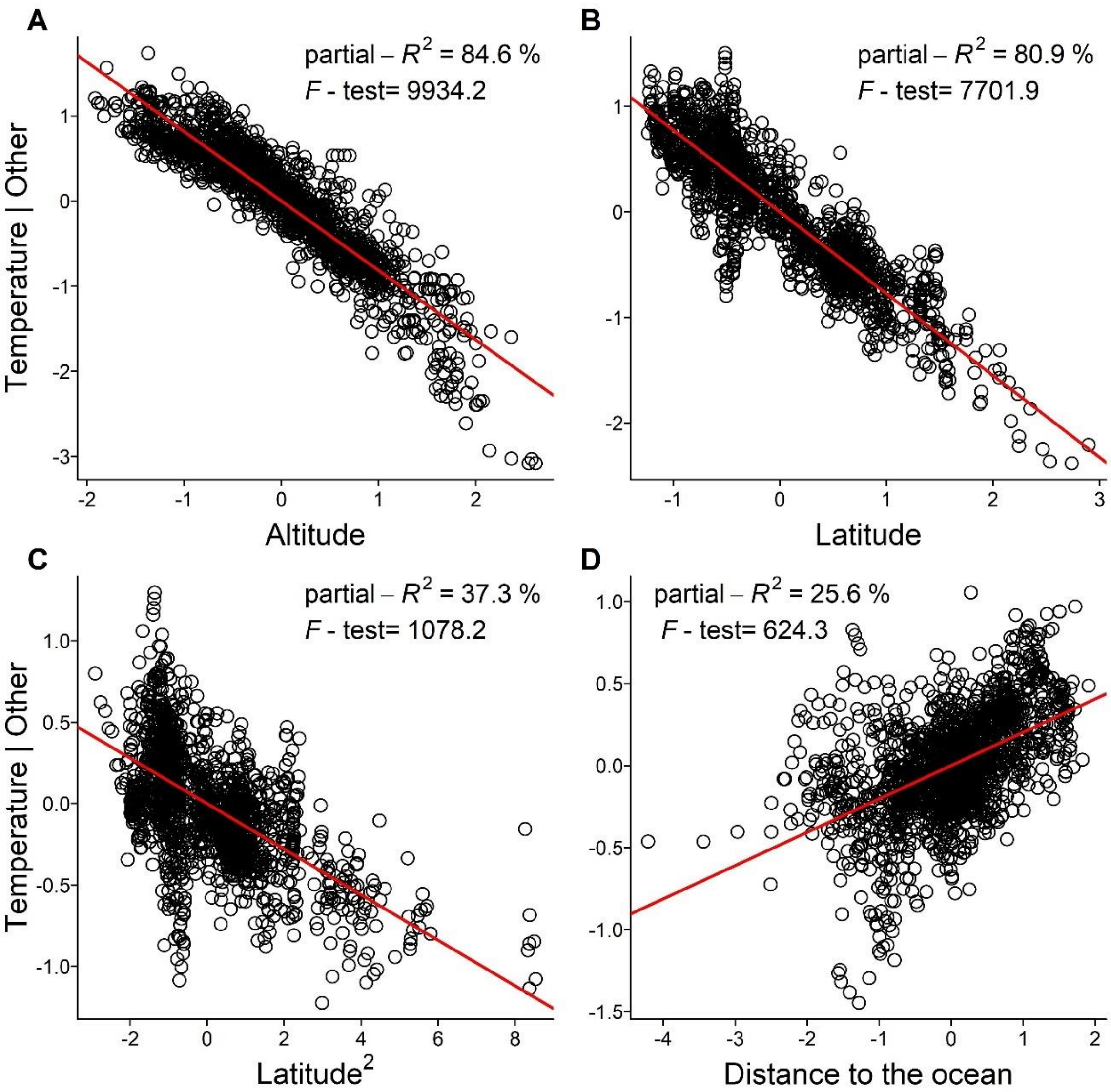

**Supplementary Figure 7. Correlation between temperature and other temperature-related variables for the studied surveys.**

Partial-regression plots presenting the results of the multiple linear regression model relating mean air temperature to (**A**) the altitude above sea level; (**B**) latitude South; (**C**) the quadratic term for latitude; and (**D**) distance to the ocean. For each variable, the partial-$R^2$ and the $F$-test evaluating the effect of each variable is also given. For both axes, values are the residuals of each variable given the presence of all other variables in the model. The addition of the quadratic term for latitude significantly improved model fit with respect to the model containing only linear terms (ΔAIC= 855).

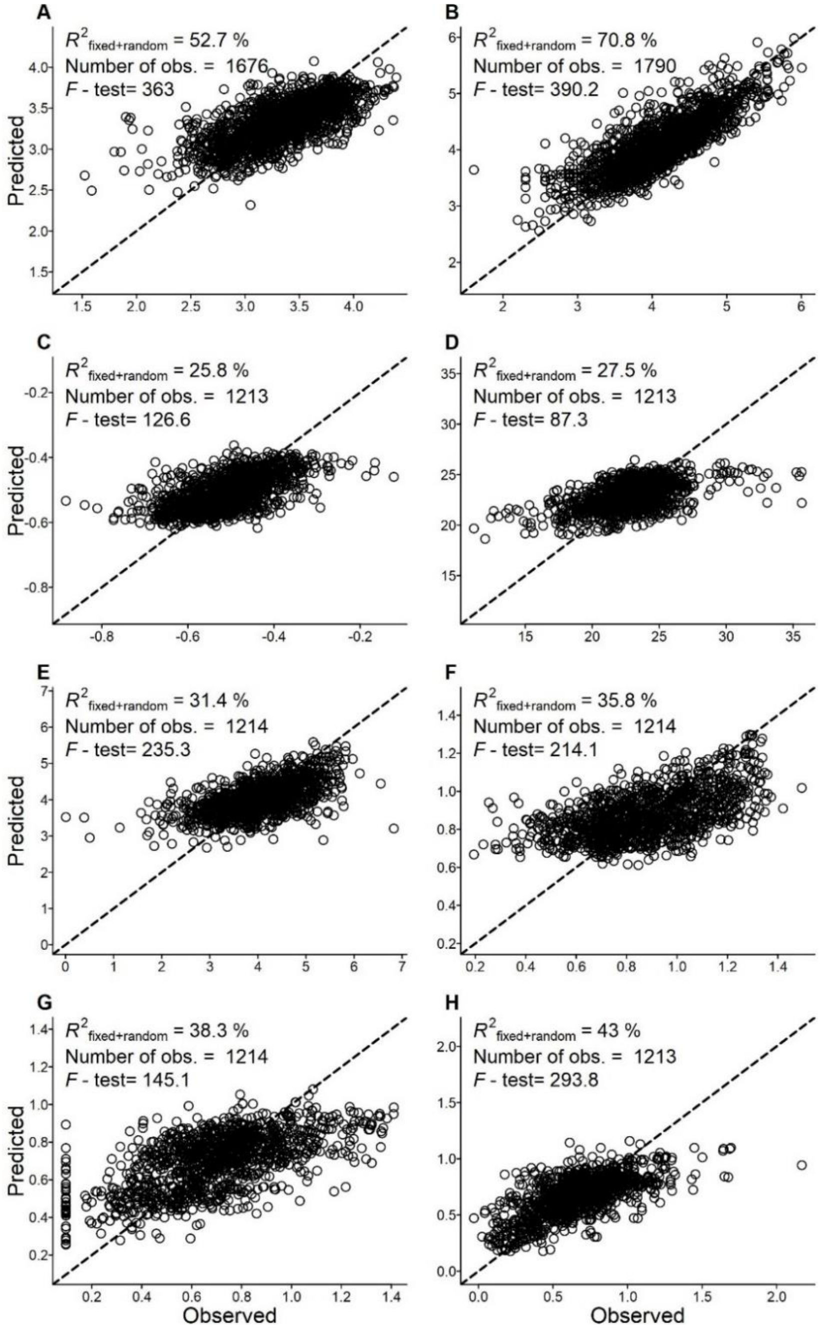

**Supplementary Figure 8. Relationship between the observed and predicted values of the models fitted to forest biomass, species richness and species properties data.** Each circle represents one survey regarding (**A**) forest biomass; (**B**) tree richness; (**C**) wood density; (**D**) maximum height; (**E**) seed mass; (**F**) ecological groups; (**G**) extinction threat; and (**H**) endemism level. The summary of the linear mixed-effects regression models is given in the top of each panel, where $R^2_{\text{fixed+random}}$ is the variation explained by the full model, *i.e*. the conditional $R^2$. Degrees of freedom: A= 1660, B= 1770, C= 1198, D–F= 1199, G= 1201, H= 1197. The dashed line represents the 1:1 ratio between observed and predicted values. Note that some of the axes were log- or power-transformed.

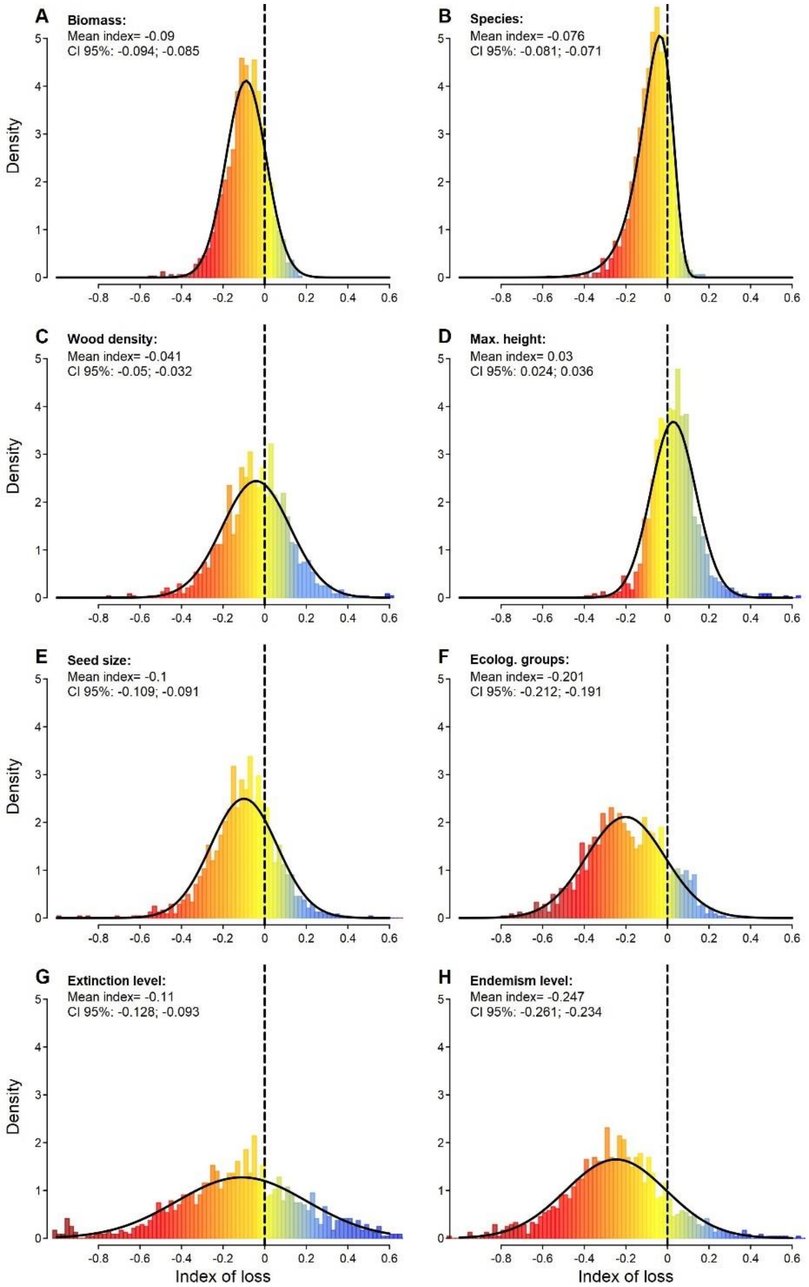

**Supplementary Figure 9. The distribution of the standardized indices of biomass, species richness and properties loss in the Atlantic Forest hotspot.** Frequency distribution (coloured bars) of the index of loss for (**A**) forest biomass; (**B**) tree richness; (**C**) wood density; (**D**) maximum height; (**E**) seed mass; (**F**) ecological groups; (**G**) extinction threat; and (**H**) endemism level, with their fits by the Normal or Weibull distributions (solid bold lines), the estimated mean and its 95% confidence interval (CI). Dashed lines separate negative indices (losses due to human-related impacts) from positive ones (gains due to human-related impacts). Bars are highlighted by colours ranging from dark red (high losses) to blue (gains). The standardized index of loss is dimensionless and is highlighted by different colours ranging from dark red (high losses) to blue (gains).

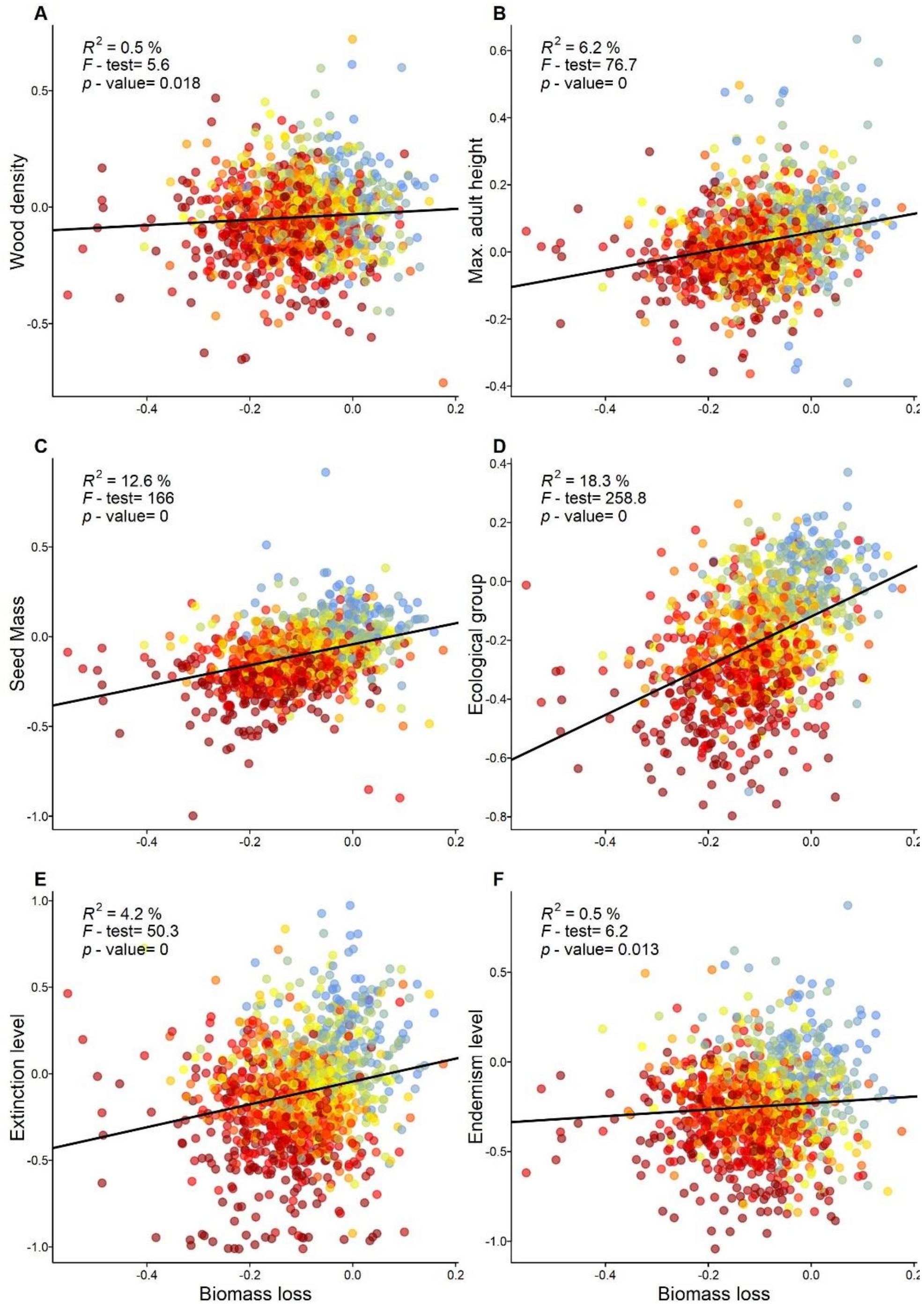

**Supplementary Figure 10. The relationship between indices of loss of forest biomass and of the community-weighted mean of each species property.**

Each point is a survey for which the loss of (**A**) wood density; (**B**) maximum height; (**C**) seed mass; (**D**) ecological groups; (**E**) threat of extinction; and (**F**) endemism level is available (*n*= 1153). The summary of the linear regression models is given in the top of each panel, where $R^2$ is the variation explained by the full model (all modes have 1,1152 degrees of freedom). The standardized index of loss is dimensionless and is highlighted by different colours ranging from dark red (high losses) to blue (gains).

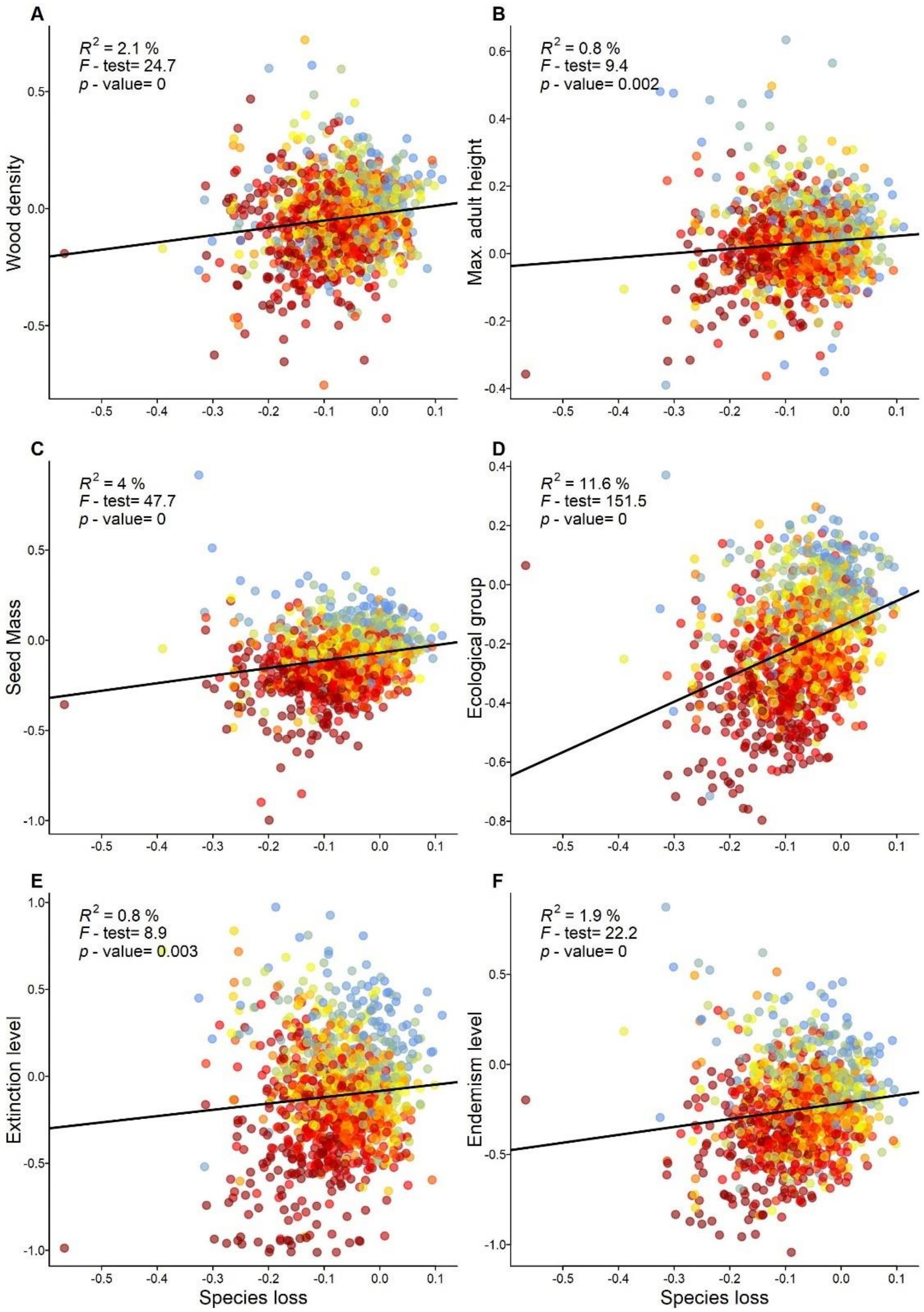

**Supplementary Figure 11. The relationship between indices of loss of species richness and of the community-weighted mean of each species property.** Each point is a survey for which the loss of (**A**) wood density; (**B**) maximum height; (**C**) seed mass; (**D**) ecological groups, (**E**) threat of extinction; and (**F**) endemism level is available ($n$= 1153). The summary of the linear regression models is given in the top of each panel, where $R^2$ is the variation explained by the full model (all modes have 1152 degrees of freedom). The standardized index of loss is dimensionless and is highlighted by different colours ranging from dark red (high losses) to blue (gains).

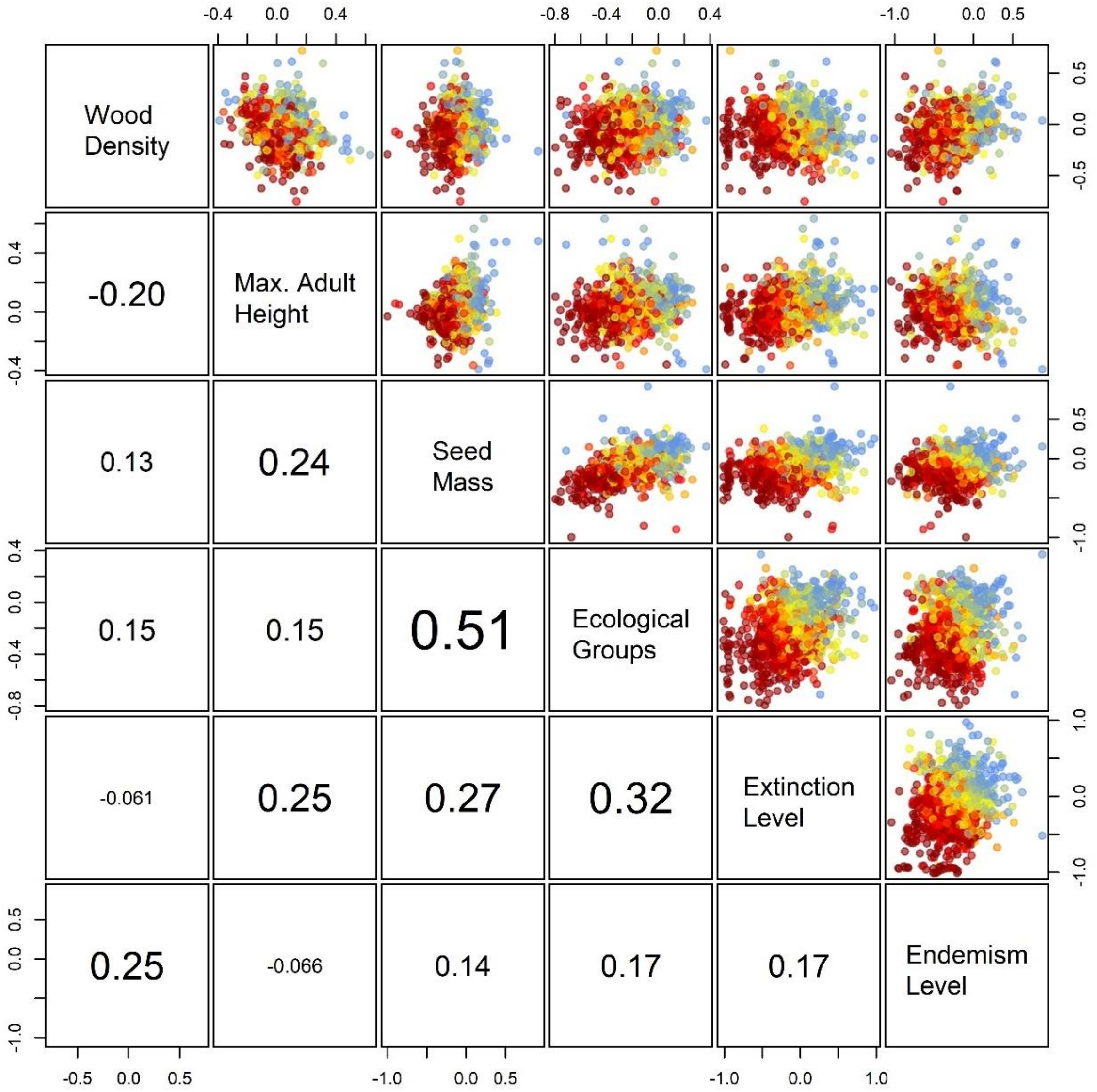

**Supplementary Figure 12. The relationship among the standardized indices of loss for multiple species properties of the Atlantic Forest hotspot.**

Above the diagonal, we present the scatterplots relating each pair of indices of loss for the six properties evaluated in this study (panels in the diagonal), in which every point represents a survey (*n*= 1213). Below the diagonal, we present the value of the Pearson's correlation index for the corresponding pair of properties. The size of the font of the correlation is proportional to the strength of the correlation between indices of loss. The standardized index of loss is dimensionless and is highlighted by different colours ranging from dark red (high losses) to blue (gains).

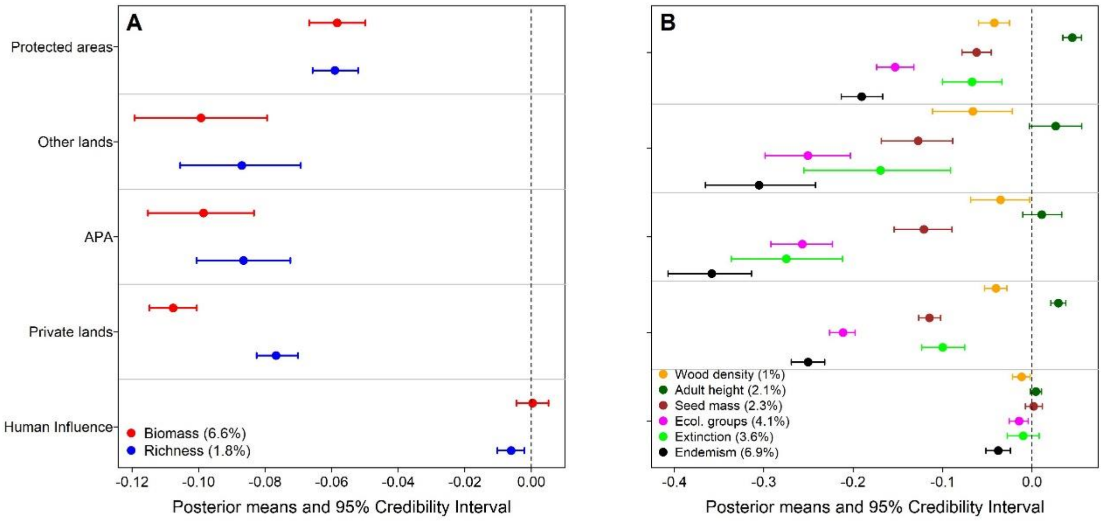

**Supplementary Figure 13. The effect of land conservation category and of the Human Influence Index on the carbon and biodiversity losses in the Atlantic Forest.**

Each point represents the posterior mean of the standardized effect sizes of the multivariate regression models containing the joint losses of (**A**) forest biomass and species richness and (**B**) of community-weighted species properties. For the categorical variable 'land conservation', posterior means refer to the estimates of the model intercept, while for the continuous variable 'human influence' the means are the estimates of the model slope. Credibility intervals (brackets) touching the dashed line reflect the lack of support of a significant effect of the co-variables. The coefficient of correlation of each model ($R^2$) is reported in parentheses. Legend: Protected areas indicate Strict protection and Sustainable use conservation units; Other lands indicate research centres, university campuses, botanical gardens, and military and indigenous lands; APA indicate private lands inside areas with sustainable use of natural resources, locally known as "Environmental Protection Areas" (APA is the acronym in Portuguese).

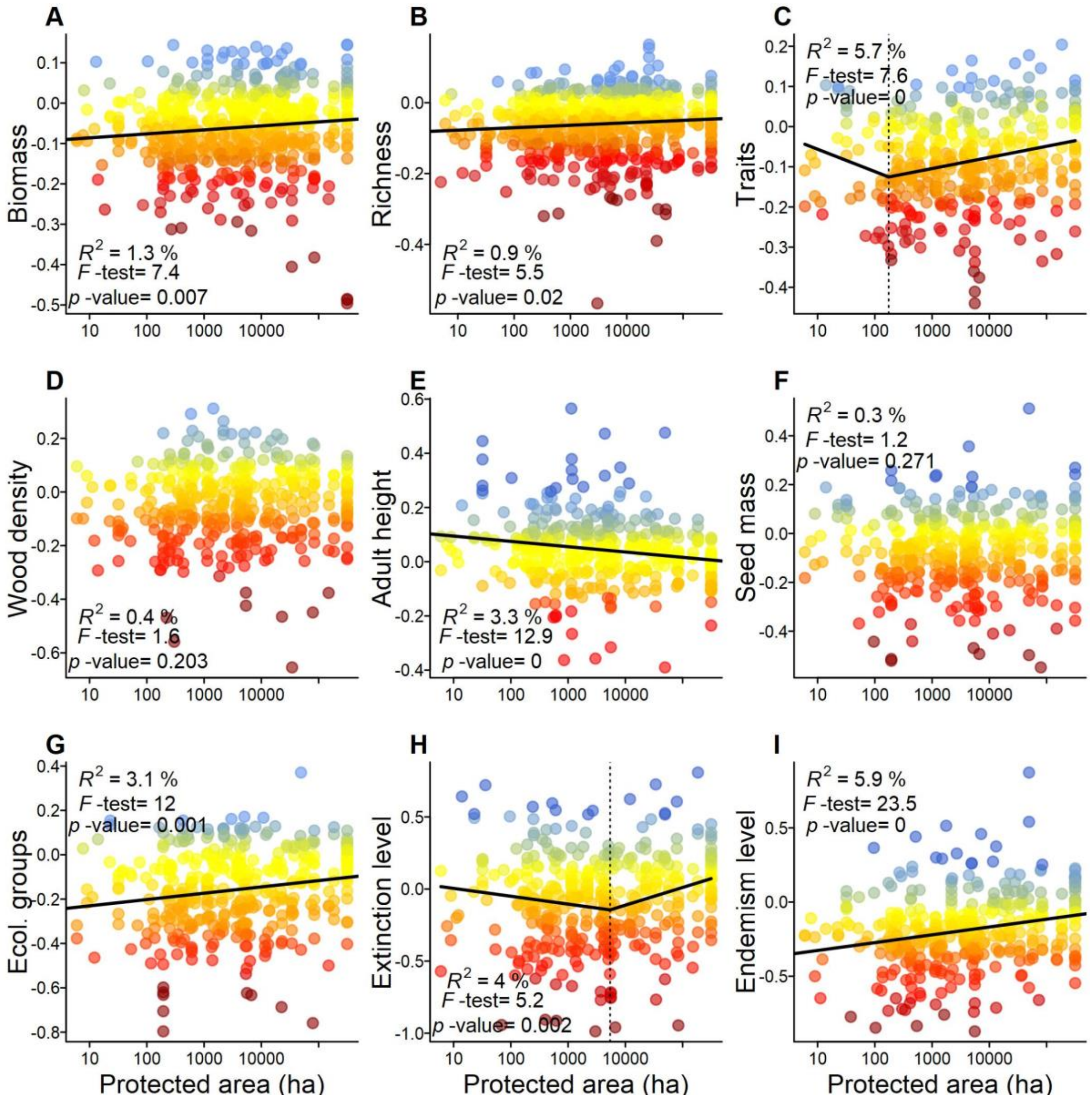

**Supplementary Figure 14. The effect of the size of the protected area on the loss of forest biomass, species richness and species properties.**

For each survey (points), panels present the indices of loss of (**A**) forest biomass (*n*= 555); (**B**) tree species richness (*n*= 617); (**C**) weighted average species property loss (*n*= 364 for all species properties); (**D**) wood density; (**E**) maximum height; (**F**) seed mass; (**G**) ecological groups; (**H**) threat of extinction; and (**I**) endemism level. The summary of the linear or of the piecewise regression models is given in the top of each panel, along with the adjusted $R^2$ of the model and its summary F-statistics. Degrees of freedom for the regression models are: A= 554, B= 616, C–H= 364.The vertical dashed line is the estimated break-point of the piecewise model, which is plotted only for the variables were this model had a better performance than the linear model. The standardized index of loss is dimensionless and is highlighted by different colours ranging from dark red (high losses) to blue (gains).

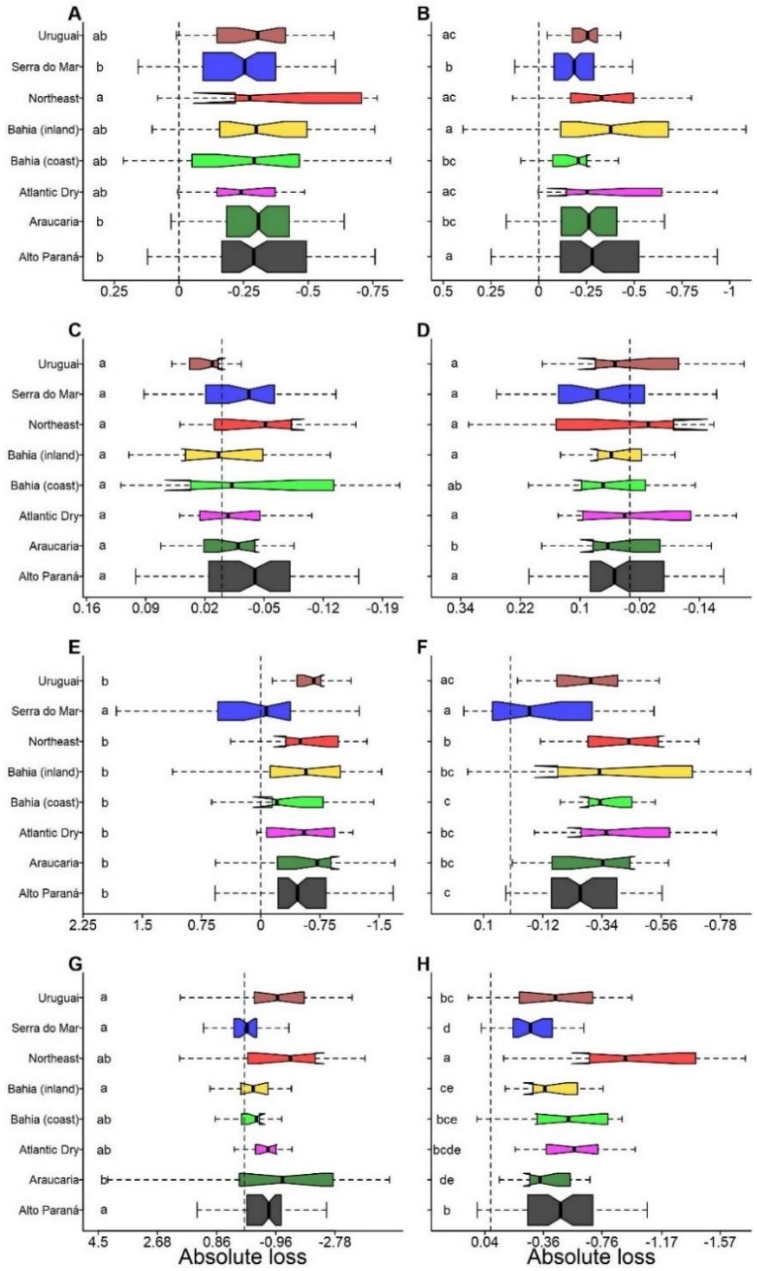

**Supplementary Figure 15. The percentage of absolute loss of biomass, species richness and species properties by biogeographical region of the Atlantic Forest.** Weighted box-and-whisker plots of (**A**) forest biomass; (**B**) species richness; (**C**) wood density; (**D**) maximum height; (**E**) seed mass; (**F**) ecological groups; (**G**) threat of extinction; and (**H**) endemism level. These plots summarize the distribution of losses for each region (*i.e*. vertical bold line, median; box limits, upper and lower quartiles; whiskers, 5 and 95% quantiles). Outliers of the distributions not presented for clarity. Dashed lines separate gains from losses due to human-related impacts. The result of the Tukey test for difference among group means (lowercase letters) is also presented. Colours represent the different regions as in Fig. S1.

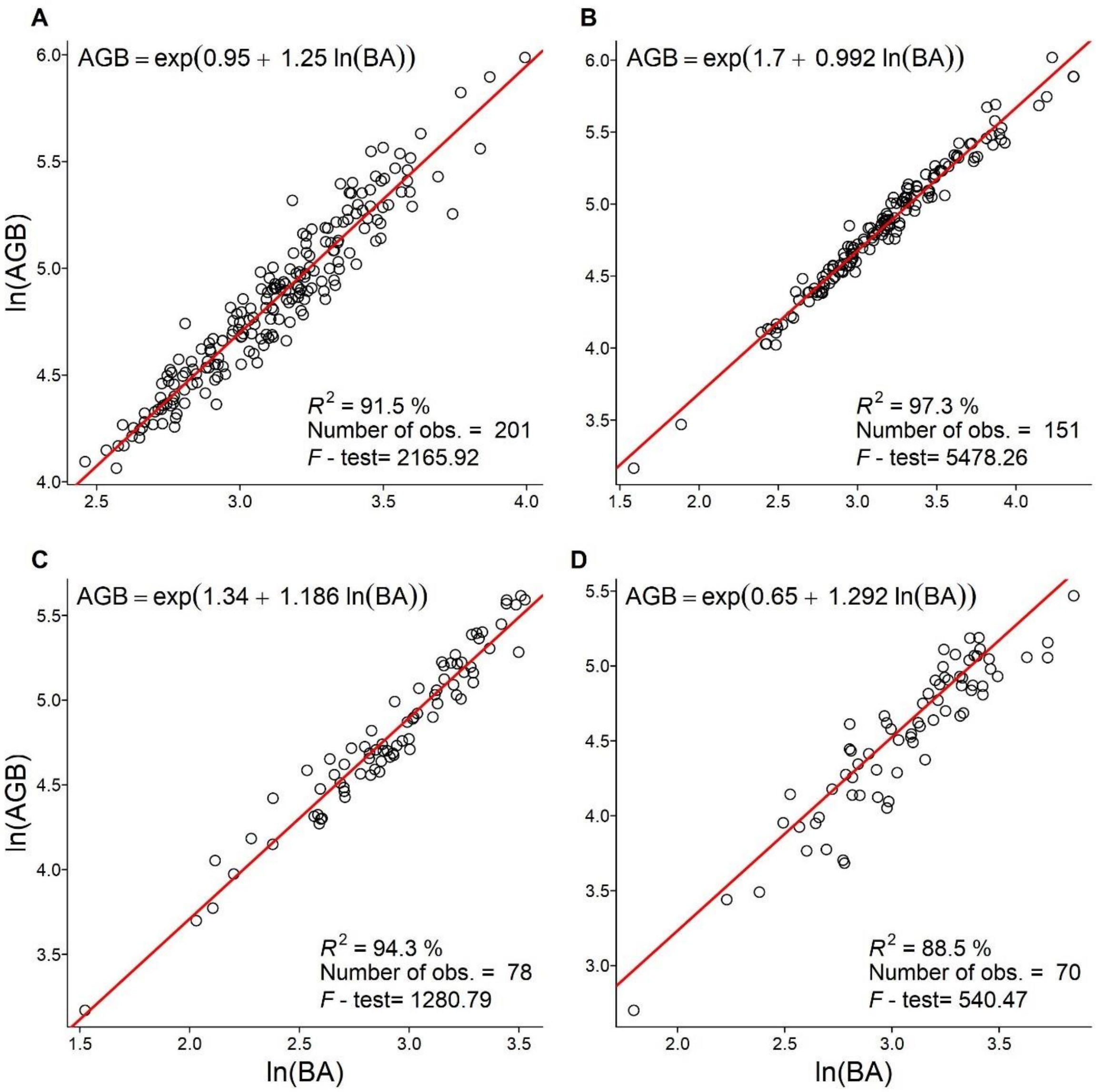

**Supplementary Figure 16. Correlation between tree basal area and above ground biomass in different Atlantic Forest formations.**

Mean prediction of the linear regression model relating basal area (in $m^2$ $ha^{-1}$) to above-ground biomass (in Mg $ha^{-1}$) for (**A**) seasonal forests; (**B**) *Araucaria* forests; (**C**) rain forests of Santa Catarina state; and (**D**) for seasonal plus rain forests of Minas Gerais state. The statistical model is given in the top of each panel and the model statistics are provided below each panel. For panel D, the legend corresponds to the result of the weighted regression model, which contained the total sampling effort (in ha) of the forest surveys as weights.

#### 4. Supplementary Notes

##### Sources of species trait information and their TreeCo reference ID

ABREU, K.M.P.; SILVA, G.F. & SILVA, A.G. 2013. Análise fitossociológica da Floresta Nacional de Pacotuba, Cachoeiro de Itapemirim, ES – Brasil. Cerne 19(1): 157-168. [TreeCo refID: 7]

ALMEIDA, S.D.; PROENÇA, C.E.; SANO, S.M. & RIBEIRO, J.F. 1998. Cerrado: espécies vegetais úteis. Embrapa-CPAC, Planaltina. 464p. [TreeCo refID: 10121]

ALMEIDA, V.C. 1996. Composição florística e estrutura do estrato arbóreo de uma floresta situada na Zona da Mata Mineira, município de Lima Duarte, MG. Dissertação (Mestrado). Universidade Federal do Rio de Janeiro, Rio de Janeiro. 89p. [TreeCo refID: 136]

AMARO, M.A. 2010. Quantificação do estoque de volume, biomassa e carbono em uma Floresta Estacional Semidecidual Montana em Viçosa, MG. Tese (doutorado). Universidade Federal de Viçosa, Viçosa. 168p. [TreeCo refID: 10003]

AMORIM, A.M.; CARVALHO, G.M. & MATOS, F. 2009. Relatório Técnico dos Diagnósticos Temáticos do Grupo FLORA - PARNA Descobrimento, Bahia, Brasil. Fundação Biodiversitas/SAVE Brasil/IESB/CEPLAC/UFMG. 81 p. [TreeCo refID: 2747]

AMORIM, A.M.; JARDIM, J.G. & FIASCHI, P. 2007. Relatório Técnico dos Diagnósticos Temáticos do Grupo FLORA - Ampliação e Estabelecimentos de novas RPPNs no sul da Bahia. Instituto de Estudos Sócio Ambientais do Sul da Bahia (IESB). [TreeCo refID: 2750]

AMORIM, I.L.; SAMPAIO, E.V.S.B. & ARAÚJO, E.L. 2005. Flora e estrutura da vegetação arbustivo-arbórea de uma área de caatinga do Seridó, RN, Brasil. Acta Botanica Brasilica 19(3): 615-623. [TreeCo refID: 2176]

ANTONIO, F. & GIULIETTI, A. 2014. A tribo Pisonieae Meisner (Nyctaginaceae) no Brasil. Boletim De Botânica 32(2): 145-268. [TreeCo refID: 10189]

ARCHANJO, K.M.P.A.; SILVA, G.F.; CHICHORRO, J.F. & SOARES, C.P.B. 2012. Estrutura do componente arbóreo da Reserva Particular do Patrimônio Natural Cafundó, Cachoeiro de Itapemirim, Espírito Santo, Brasil. Floresta 42 (1): 145-160. [TreeCo refID: 6]

ARRUDA, D.M.; BRANDÃO, D.O.; COSTA, F.V.; TOLENTINO, G.S.; BRASIL, R.D.; D´ÂNGELO NETO, S. & NUNES, Y.R.F. 2011. Structural aspects and floristic similarity among tropical dry forest fragments with different management histories in northern Minas Gerais, Brazil. Revista Árvore 35(1): 131-142. [TreeCo refID: 62]

ASSIS, L.C.S. 2009. Sistematica e Filosofia: filogenia do complexo Ocotea e revisao do grupo Ocotea indecora (Lauraceae). Tese (doutorado). Universidade de São Paulo, São Paulo. 238 p. [TreeCo refID: 10004]

BARBÉRIO, M. 2013. Maturação de sementes de Andira fraxinifolia Benth. (Fabaceae) em uma área de restinga. Dissertação (Mestrado). Instituto de Botânica, São Paulo. 53p. [TreeCo refID: 10409]

BELLO, C.; GALETTI, M.; MONTAN, D.; PIZO, M.A.; MARIGUELA, T.C.; CULOT, L.; BUFALO F.S.; LABECCA, F.M.; PEDROSA, F.R.V.; CONSTANTINI, R.; EMER, C.; SILVA, W.R.; SILVA, F.R.; OVASKAINEN, O. & JORDANO, P. 2017. Atlantic frugivory: a plant–frugivore interaction data set for the Atlantic Forest. Ecology 98(6): 1729. [TreeCo refID: 10418]

BENVENUTI-FERREIRA, G.; COELHO, G.C. 2009. Floristics and structure of the tree component in a Seasonal Forest remnant, Chiapetta, Rio Grande do Sul State, Brazil. Revista Brasileira de Biociências 7(4): 344-353. [TreeCo refID: 754]

BERG, C. 2001. Moreae, Artocarpeae, and Dorstenia (Moraceae), with Introductions to the Family and Ficus and with Additions and Corrections to Flora Neotropica Monograph 7. Flora Neotropica 83: iii-346. [TreeCo refID: 10138]

BERG, C.; ROSSELLI, P.F. & DAVIDSON, D. 2005. Cecropia. Flora Neotropica 94: 1-230. [TreeCo refID: 10139]

BERNACCI, L.C.; FRANCO, G.A.D.C. & ÁRBOCZ, G.F. 2006. O efeito da fragmentação florestal na composição e riqueza de árvores na região da Reserva Morro Grande (Planalto de Ibiúna, SP). Revista do Instituto Florestal 18: 121-166. [TreeCo refID: 2690]

BIANCHINI, E.B.; POPOLO, R.S.; DIAS, M.C. & PIMENTA, J.A. 2003. Diversidade e estrutura de espécies arbóreas em uma área alagável do município de Londrina, sul do Brasil. Acta Botanica Brasilica 17(3): 405-419. [TreeCo refID: 449]

BIGIO, N.C. & SECCO, R.S. 2012. As espécies de Pera (Euphorbiaceae s.s) na Amazônia brasileira. Rodriguésia 63(1): 163-207. [TreeCo refID: 10157]

BORGHI, W.A.; MARTINS, S.S.; DEL QUIQUI, E.M. & NANNI, M.R. 2004. Caracterização e avaliação da mata ciliar à montante da Hidrelétrica de Rosana, na Estação Ecológica do Caiuá, Diamante do Norte, PR. Cadernos de Biodiversidade 4(2): 9-18. [TreeCo refID: 454]

BOSA, D.M. 2011. Composição florística e estrutural de comunidade arbórea de floresta ombrófila densa montana no município de Morro Grande, Santa Catarina. Dissertação (Mestrado). Universidade do Extremo Sul Catarinense, Criciúma. 89p. [TreeCo refID: 833]

BOTREL, R.T.; OLIVEIRA-FILHO, A.T.; RODRIGUES, L.A. & CURI, N. 2002. Composição florística e estrutura da comunidade arbórea de um fragmento de floresta estacional semidecidual em Ingaí, MG, e a influência de variáveis ambientais na distribuição das espécies. Revista Brasileira de Botânica 25(2): 195-213. [TreeCo refID: 66]

BRANDÃO, C.F.L.S.; MARANGON, L.C.; FERREIRA, R.L.C. & LINS-E-SILVA, A.C.B. 2009. Phytosociological structure and successional classification of arboreus component in a fragment of a dense ombrophylous forest, Igarassu – Pernambuco. Revista Brasileira de Ciências Agrárias 4(1): 55-61. [TreeCo refID: 1515]

BROTTO, M.L.; CERVI, A.C. & SANTOS, E.P. 2013. O gênero Ocotea (Lauraceae) no estado do Paraná, Brasil. Rodriguésia 64(3): 495-525. [TreeCo refID: 10008]

BROWN, S. 1997. Appendix 1 - List of wood densities for tree species from tropical America, Africa, and Asia. In: Estimating Biomass and Biomass Change of Tropical Forests: a Primer. UN FAO Forestry Paper, 134. <http://www.fao.org/docrep/w4095e/w4095e00.htm#Contents > Acessado em: 23 fev 2016. [TreeCo refID: 10009]

BUDKE, J.C. 2007. Pulsos de inundação, padrões de diversidade e distribuição de espécies arbóreas em uma floresta ribeirinha no sul do Brasil, Tese (Doutorado). Universidade Federal do Rio Grande do Sul, Porto Alegre, RS. 195p. [TreeCo refID: 663]

BUDKE, J.C.; JARENKOW, J. A. & OLIVEIRA-FILHO, A.T. 2008. Tree community features of two stands of riverine under different flooding regimes in Southern Brazil. Flora 203(2): 162-174. [TreeCo refID: 664]

CADDAH, M.K. 2013. Estudos taxônomicos e filogenéticos em Miconia sect. Discolor (Melastomataceae, Miconieae). Tese (Doutorado). Universidade Estadual de Campinas, Campinas. 261 p. [TreeCo refID: 10186]

CALIARI, C.P. 2013. Estudos em Myrtaceae do Estado de São Paulo: Myrcia seção Gomidesia. Dissertação (Mestrado). Escola Superior de Agricultura "Luiz de Queiroz", Piracicaba. 129p. [TreeCo refID: 10010]

CAMPOE, O.C. 2008. Efeito de práticas silviculturais sobre a produtividade líquida de madeira, o índice de área foliar e a eficiência do uso da luz em de restauração da Mata Atlântica. Dissertação (Mestrado). Escola Superior de Agricultura "Luiz de Queiroz", Piracicaba. 120p. [TreeCo refID: 10011]

CAMPOS, J.B. & SOUZA, M.C. 2003. Potential for natural Forest regeneration from seed bank in an Upper Paraná River Floodplain, Brazil. Brazilian Archives of Biology and Technology 46(4): 625-639. [TreeCo refID: 2686]

CAMPOS, J.B.; ROMAGNOLO, M.B. & SOUZA, M.C. 2000. Structure, composition and spatial distribution of tree species in a remnant of the semideciduous seasonal alluvial forest of the upper Paraná River floodplain. Brazilian Archives of Biology and Technology 43(2): 185-194. [TreeCo refID: 509]

CANALE, G.R.; SUSCKE, P.; ROCHA-SANTOS, L.; SÃO BERNARDO, C.S.; KIERULFF, M.C.M. & CHIVERS, D.J. 2016. Seed Dispersal of Threatened Tree Species by a Critically Endangered Primate in a Brazilian Hotspot. Folia Primatologica 87(3): 123-140. [TreeCo refID: 10012]

CAPRETZ, R.L.; BRITEZ, R.M.; BEBBER, D.P.; REGINATO, M. & ZWIENER, V.P. (Unpublished data). Floristic and structural patterns along a successional chronosequence in the Atlantic Forest of Southern Brazil. UFPR. Curitiba. [TreeCo refID: 2714]

CARAUTA, J.P.P. & DIAZ, B.E. 2002. Figueiras no Brasil. Editora UFRJ, Rio de Janeiro. 212p. [TreeCo refID: 10013]

CARMELLO-GUERREIRO, S.M. & PAOLI, A.A.S. 2005. Anatomy of the pericarp and seed-coat of Lithraea molleoides (Vell.) Engl. (Anacardiaceae) with taxonomic notes. Brazilian Archives of Biology and Technology, 48(4): 599-610. [TreeCo refID: 10014]

CARUZO, M.B.R. 2010. Sistematica de Croton sect. Cleodora (Euphorbiaceae s.s.). Tese (Doutorado). Universidade de Sao Paulo, São Paulo. 273p. [TreeCo refID: 10015]

CARVALHO-OKANO, R.M. 1992. Estudos taxonômicos do gênero Maytenus Mol. emend. Mol. (Celastraceae) do Brasil extra-amazônico. Tese (Doutorado). Universidade Estadual de Campinas, Campinas. 261p. [TreeCo refID: 10184]

CARVALHO, A.F. 2013. Caracterização da madeira e do carvão vegetal produzido a partir de cinco espécies florestais utilizadas na região de Biguaçu, SC. Dissertação (Mestrado). Universidade do Estado de Santa Catarina, Lages. 142p. [TreeCo refID: 10016]

CARVALHO, A.M. & AMORIM, A.M. 1996. Composição Florística e Estrutura da Mata da Esperança no município de Ilhéus, BA. In: XLVII Congresso Nacional de Botânica. [TreeCo refID: 2749]

CARVALHO, D.A.; OLIVEIRA-FILHO, A.T. & VILELA, E.A. 1999. Floristics and phytosociology of the arboreal-shrubby vegetattion of a deciduous riparian forest of the low Paranaiba (Santa Vitoria, Minas Gerais). Revista Arvore 23(3): 311-320. [TreeCo refID: 151]

CARVALHO, D.A.; OLIVEIRA-FILHO, A.T.; VAN DEN BERG, E.; FONTES, M.A.L.; VILELA, E.A.; MARQUES, J.J.S.G.S. M. & CARVALHO, W.A.C. 2005. Variações florísticas e estruturais do componente arbóreo de uma floresta ombrófila alto-montana às margens do Rio Grande, Bocaina de Minas, MG, Brasil. Acta Botanica Brasilica 19 (1): 91-109. [TreeCo refID: 69]

CARVALHO, D.A.; OLIVEIRA-FILHO, A.T.; VILELA, E.A. & CURI, N. 2000. Florística e estrutura da vegetação arbórea de um fragmento de floresta semidecidual às margens do reservatório da Usina Hidrelétrica Dona Rita (Itambé do Mato Dentro, MG). Acta Botanica Brasilica 14(1): 37-55. [TreeCo refID: 68]

CARVALHO, D.A.; OLIVEIRA-FILHO, A.T.; VILELA, E.A.; CURI, N.; VAN DEN BERG, E.; FONTES M.A.L. & BOTEZELLI, L. 2005. Distribuição de espécies arbóreo-arbustivas ao longo de um gradiente de solos e topografia em um trecho de floresta ripária do rio São Francisco em Três Marias, MG, Brasil. Revista Brasileira de Botânica 28(2): 329-345. [TreeCo refID: 150]

CARVALHO, E.S.; SANTOS, J.P.A. SILVA, R.R.; MIGUEZ, L.S.; SOUZA, M.O. & MENDONÇA, A.V.R. 2018. Biometria de frutos e sementes de Lecythis lurida (Miers) S.A. Mori. Anais do VII Congresso Florestal Latino-americano, Vitória. https://even3.blob.core.windows.net/anais/95401.pdf [TreeCo refID: 10416]

CARVALHO, G.M. 2011. Influência de processos estocásticos sobre a estruturação de comunidades em Floresta de Tabuleiros Bahia, Brasil. Dissertação (Mestrado), Universidade Estadual de Santa Cruz, Ilhéus. 61p. [TreeCo refID: 2702]

CARVALHO, L.D'A.F. 1988. Revisão Taxonômica das Especies de Solanum das Seções Cernuum e Lepidotum-Solanaceae. Doctoral Thesis, Universidade Estadual de Campinas, Campinas. 228p. [TreeCo refID: 10167]

CARVALHO, P.E.R. 2004. Pau-Marfim – Balfourodendron riedelianum. Embrapa Florestas. Colombo, PR. Circular Técnica n° 93. <http://www.cnpf.embrapa.br/publica/circtec/edicoes/circtec93.pdf > [TreeCo refID: 10017]

CARVALHO, W.A.C. 2002. Variações da composição e estrutura do compartimento arbóreo da vegetação de oito fragmentos de floresta semidecídua do vale do Alto Rio Grande, MG. Dissertação (Mestrado). Universidade Federal de Lavras, Lavras. 168p. [TreeCo refID: 154]

CARVALHO, W.A.C. 2010. Diversidade do estrato arbóreo-arbustivo de sete comunidades de floresta ombrófila altomontanas da APA Fernão Dias, MG, Brasil. Tese (Doutorado). Universidade Federal de Minas Gerais, Belo Horizonte. 72p. [TreeCo refID: 2556]

CATHARINO, E.L.M.; BERNACCI, L.C.; FRANCO, G.A.D.C. & DURIGAN, G.; METZGER, J.P. 2006. Aspectos da composição e diversidade do componente arbóreo das florestas da Reserva Florestal do Morro Grande, Cotia, SP. Biota Neotropica 6(2): ISSN 1806-129X. [TreeCo refID: 965]

CERQUEIRA, R.M. 2005. Florística e estrutura de um fragmento de floresta estacional semidecídua montana no município de Itatiba, SP. Dissertação (Mestrado). Universidade de Campinas, Campinas. 106p. [TreeCo refID: 2810]

CÉSAR, O. & LEITÃO–FILHO, H.F. 1990b. Estudo fitossociológico de mata mesófila semidecídua na Fazenda Barreiro Rico, município de Anhembi, SP. Revista Brasileira de Biologia 50: 443-452. [TreeCo refID: 970]

CHAGAS, A.P. 2014. Ingeae Benth. (Leguminosae-Mimosoideae) no Espírito Santo, Brasil. Dissertação (Mestrado). Universidade Federal de Viçosa, Viçosa. 109p. [TreeCo refID: 10181]

CHAVE, J.; COOMES, D.A.; JANSEN, S.; LEWIS, S.L.; SWENSON, N.G. & ZANNE, A.E. 2009. Towards a worldwide wood economics spectrum. Ecology Letters 12(4): 351-366. (ZANNE, A.E. et al. 2009. Global Wood Density Database. Dryad Digital Repository. https://doi.org/10.5061/dryad.234) [TreeCo refID: 10254]

CHIMELO, J.P.; MAINIERI, C.; HAHUZ, M.A.R.; PESSOA, A.L. 1976.Madeiras do município de Aripuanã, Estado de Mato Grosso. I - Caracterização anatômica e aplicações. Acta Amazonica, 6(4): 95-106 apud PAULA, J.E. & ALVES, J.L.H. 2007. 897 Madeiras nativas do Brasil: anatomia, dendrologia, dendrometria, produção, uso. Editora Cinco Continentes, Porto Alegre. 438p. [TreeCo refID: 10124]

CINTRA, T.C. 2009. Avaliações energéticas de espécies florestais nativas plantadas na região do Médio Paranapanema, SP. Dissertação (Mestrado). Escola Superior de Agricultura "Luiz de Queiroz", Piracicaba. 84p. [TreeCo refID: 10018]

COELHO, R.L.G. 2008. Estudos Taxonómicos em Matayba Aubl. sect. Matayba (Sapindaceae). Dissertação (Mestrado). Universidade Estadual de Campinas, Campinas. 172p. [TreeCo refID: 10019]

COELHO, R.L.G. 2014. CoeEstudos sistemáticos das espécies neotropicais de Allophylus L. (Sapindaceae). Tese (Doutorado). Universidade Estadual de Campinas, Campinas. 479p. [TreeCo refID: 10403]

COELHO, S.; CARDOSO-LEITE, E. & CASTELLO, A.C.D. 2016. Composição florística e caracterização sucessional como subsídio para conservação e manejo do PNMCBio, Sorocaba – SP. Ciência Florestal 26(1): 331-344. [TreeCo refID: 3824]

COLONETTI, S.; CITADINI-ZANETTE, V.; MARTINS, R.; SANTOS, R.D.; ROCHA, E.; & JARENKOW, J.A. 2009. Floristic composition and phytosociological structure in a submontane ombrophilous dense forest at São Bento river dam, Siderópolis, Santa Catarina State. Acta Scientiarum-Biological Sciences 31(4): 397-405. [TreeCo refID: 813]

COSMO, N.L.; NOGUEIRA, A.C.; LIMA, J.G. & KUNIYOSHI, Y.S. 2010. Morfologia de fruto, semente e plântula de Sebastiania commersoniana, Euphorbiaceae. Floresta 40(2): 419-428. [TreeCo refID: 10411]

COSTA-FILHO, S.V.S. 2016. Estrutura, composição e diversidade florística dos estratos verticais de um fragmento de floresta ombrófila mista em Campo do Tenente, Paraná. Monografia (Graduação). Universidade Federal do Paraná, Curitiba. 13 p. [TreeCo refID: 3834]

COSTA, J.C.A. 2004. Fixação de carbono e produção de biomassa pela cupiúva (Tapirira guianensis Aubl.), em um fragmento manejado de mata atlântica, município de Goiana-PE. Dissertação (Mestrado). Universidade Federal Rural do Pernambuco, Recife. 109p. [TreeCo refID: 10020]

COSTA, T.G.; BIANCHI, M.L.; PROTÁSIO, T.P.; TRUGILHO, P.F. & PEREIRA, A.J. 2014. Qualidade da madeira de cinco espécies de ocorrência no cerrado para produção de carvão vegetal. CERNE 20(1): 37-46. [TreeCo refID: 10021]

COWAN, R. 1967. Swartzia (Leguminosae, Caesalpinioideae Swartzieae). Flora Neotropica 1: 1-228. [TreeCo refID: 10435]

CUNHA, M.P.S.C.; PONTES, C.L.F.; CRUZ, I. A.; CABRAL, M.T F.D.; CUNHA NETO, Z.B. & BARBOSA, A.P.R. 1989. Estudo químico de 55 espécies lenhosas para geração de energia em caldeiras. In: 3° encontro Brasileiro em madeiras e em estruturas de madeira: Anais, São Carlos: 2: 93-121. [TreeCo refID: 10022]

DALANESI, P.E.; OLIVEIRA-FILHO, A.T. & FONTES, M.A.L. 2004. Flora e estrutura do componente arbóreo da floresta do Parque Ecológico Quedas do Rio Bonito, Lavras – MG, e correlações entre a distribuição das espécies e variáveis ambientais. Acta Botanica Brasilica 18(4): 737-757. [TreeCo refID: 70]

DALY, D.C. 1990. The genus Tetragastris and the forests of Eastern Brazil: studies in Neotropical Burseraceae III. Kew Bulletin 45(1): 179-194. [TreeCo refID: 10023]

DALY, D.C. 1999. Notes on Trattinnickia, including a Synopsis in Eastern Brazil's Atlantic Forest Complex. Studies in Neotropical Burseraceae IX. Kew Bulletin 54(1): 129-137. [TreeCo refID: 10024]

DAMÁSIO, R.A.P.; PEREIRA, B.L.C.; OLIVEIRA, A.C.; CARDOSO, M.T.; VITAL, B.R. & CARVALHO, A.M.L.M. 2013. Caracterização anatômica e qualidade do carvão vegetal da madeira de pau-jacaré (Piptadenia gonoacantha). Pesquisa Florestal Brasileira 33(75): 261-267. [TreeCo refID: 10025]

DAVIDE, A.C.; TONETTI, O.A.O. & SILVA, E.A.A.D. 2011. Improvement to the physical quality and imbibition pattern in seeds of candeia (Eremanthus incanus (Less.) Less.). CERNE 17(3): 321-326. [TreeCo refID: 10026]

DIAS-NETO, O.C.; SCHIAVINI, I.; LOPES, S.F.; VALE, V.S.; GUSSON, A.E. & OLIVEIRA, A. P. 2009. Estrutura fitossociológica e grupos ecológicos em fragmento de floresta estacional semidecidual, Uberaba, Minas Gerais, Brasil. Rodriguésia 60(4): 1087-1100. [TreeCo refID: 160]

DIAS, M.C. 1988. Estudos taxonomicos do genero Xylopia L. (Annonaceae) no Brasil extra-amazonico. Dissertação (Mestrado). Universidade Estadual de Campinas, Campinas. 183p. [TreeCo refID: 10170]

DUARTE, M.C. 2010. Análise filogenética de Eriotheca Schott & Endl. e gêneros afins (Bombacoideae, Malvaceae) e estudo taxonômico de Eriotheca no Brasil. Tese (Doutorado). Instituto de Botânica da Secretaria de Estado do Meio Ambiente, São Paulo. 190p. [TreeCo refID: 10129]

DUARTE, T.G. 2007. Florística, fitossociologia e relações solo-vegetação em Floresta Estacional Decidual em Barão de Melgaço, Pantanal de Mato Grosso. Tese (Doutorado). Univeridade Federal de Viçosa, Viçosa. 162p. [TreeCo refID: 1959]

DURIGAN, G.; BAITELLO, J.B.; FRANCO, G.A.D.C. & SIQUEIRA, M.F. 2004. Plantas do Cerrado Paulista: imagens de uma paisagem ameaçada. Ed. Páginas e Letras, São Paulo. 475p. [TreeCo refID: 10158]

ÉDER-SILVA, E. & ARAÚJO, D.R. 2014. Physiological quality, morphometric aspects and chromosome number of the species Talisia esculenta Radlk. Revista Verde de Agroecologia e Desenvolvimento Sustentável 9(3):275-82. [TreeCo refID: 10413]

ESPÍRITO-SANTO, F.D.B.; OLIVEIRA-FILHO, A.T.; MACHADO, E.L.M.; SOUZA, J.S.; FONTES, M.A.L. & MARQUES, J.J.G.S.M. 2002. Variáveis ambientais e a distribuição de espécies arbóreas em um remanescente de floresta estacional semidecídua montana no campus da Universidade Federal de Lavras, MG. Acta Botanica Brasilica 16(3): 331-356. [TreeCo refID: 161]

FAGUNDES, L.M.; CARVALHO, D.A.; VAN DEN BERG, E.; MARQUES, J.J.G.S.M. & MACHADO, E.L.M. 2007. Florística e estrutura do estrato arbóreo de dois fragmentos de florestas decíduas às margens do rio Grande, em Alpinópolis e Passos, MG. Acta Botanica Brasilica 21(1): 65-78. [TreeCo refID: 108]

FARAH, F.T. 2009. Vinte anos de dinâmica em um hectare de Floresta Estacional Semidecidual. Tese (Doutorado). Universidade Estadual de Campinas, Campinas. 130p. [TreeCo refID: 986]

FARIA JUNIOR, J.E.Q. 2010. O gênero Eugenia L. (Myrtaceae) nos estados de Goiás e Tocantins, Brasil. Dissertação (Mestrado). Universidade de Brasília, Brasília. 266p. [TreeCo refID: 10031]

FARIA JUNIOR, J.E.Q. 2014. Revisão Taxonômica e Filogenia de Eugenia sect. Pilothecium (Kiaersk) D.Legrand (Myrtaceae). Tese (Doutorado). Universidade de Brasília, Brasília. 215p. [TreeCo refID: 10032]

FERNANDES, I. 1997. Taxonomia e fitogeografia de Cyatheaceae e Dicksoniaceae nas Regiões Sul e Sudeste do Brasil. Tese (Doutorado). Universidade de São Paulo, São Paulo. 435p. [TreeCo refID: 10033]

FERREIRA JUNIOR, M. & VIEIRA, A.O.S. 2015. Tree and shrub Rubiaceae Juss. family species along Tibagi river basin, Paraná State, Brazil. Hoehnea 42(2): 289-336. [TreeCo refID: 10034]

FERREIRA, P.I.; GOMES, J.P.; BATISTA, F.; BERNARDI, A.P.; COSTA, N.C.F.D.; BORTOLUZZI, R.L.D.C. & MANTOVANI, A. 2013. Potential species for recovery of permanent preservation areas in the highlands of Santa Catarina state, Brazil. Floresta e Ambiente 20(2): 173-182. [TreeCo refID: 814]

FIASCHI, P. & PIRANI, J.R. 2007. Estudo taxonômico do gênero Schefflera J.R. Forst. & G. Forst (Araliaceae) na região sudeste do Brasil. Boletim de Botânica da Universidade de São Paulo 25: 95-142. [TreeCo refID: 10163]

FIGLIOLA, M.B. & PINA-RODRIGUES, F.C.M. 1995. Manejo de sementes de espécies arbóreas. IF. Serie Registros 15: 1-59. [TreeCo refID: 10035]

FIGUEIREDO FILHO, A.; ORELLANA, E.; NASCIMENTO, F.; DIAS, A.N. & INOUE, M.T. 2011. Produção de sementes de Araucaria angustifolia em plantio e em floresta natural no Centro-sul do Estado do Paraná. Floresta 41(1): 155-162. [TreeCo refID: 10405]

FONSECA, C.R. & CARVALHO, F.A. 2012. Aspectos florísticos e fitossociológicos da comunidade arbórea de um fragmento urbano de floresta Atlântica (Juiz de Fora, MG, Brasil). Bioscience Journal 28(5): 820-832. [TreeCo refID: 116]

FONTES, M.A.L. 2008. Dinâmica de comunidades arbóreas de florestas alti-montanhas de Minas Gerais. Tese (Doutorado). Universidade Federal de Minas Gerais, Belo Horizonte. 85p. [TreeCo refID: 3597]

FORERO, E. 1983. Connaraceae. Flora Neotropica 36: 1-207. [TreeCo refID: 10135]

FORGIARINI, C.; SOUZA, A.F.; LONGHI, S.J. & OLIVEIRA, J.M. 2014. In the lack of extreme pioneers: trait relationships and ecological strategies of 66 subtropical tree species. Journal of Plant Ecology 8(4): 359-367. [TreeCo refID: 10037]

FRANÇA, G.S. & STEHMANN, J.R. 2004. Composição florística e estrutura de componente arbóreo de uma floresta altimontana no município de Camanducaia, Minas Gerais, Brasil. Revista Brasileira de Botânica 27 (1): 19-30. [TreeCo refID: 71]

FRANÇA, M.B. 2002. Modelagem de biomassa florestal através do padrão espectral no Sudoeste da Amazônia. Dissertação (Mestrado). Instituto Nacional de Pesquisas da Amazônia/Fundaçãoo Universidade Federal do Amazonas, Manaus, Amazonas. 106p. [TreeCo refID: 10038]

FREITAS, M.F. & KINOSHITA, L.S. 2015. Myrsine (Myrsinoideae-Primulaceae) no sudeste e sul do Brasil. Rodriguésia 66(1): 167-189. [TreeCo refID: 10187]

FURLAN, A. & GIULIETTI, A.N. 2014. A tribo Pisonieae Meisner (Nyctaginaceae) no Brasil. Boletim de Botânica 32(2): 145-268. [TreeCo refID: 10039]

GALETTI, M.; PIZO, M.A. & MORELLATO, L.P.C. 2011. Diversity of functional traits of fleshy fruits in a species-rich Atlantic rain forest. Biota Neotropica 11(1): 181-193. [TreeCo refID: 10040]

GANDOLFI, S. 2000. História Natural de uma floresta estacional semidecidual no município de Campinas (São Paulo, Brasil). Tese (Doutorado). Universidade de Campinas, Campinas. 520p. [TreeCo refID: 2631]

GENTRY, A. 1992. Bignoniaceae: Part II (Tribe Tecomeae). Flora Neotropica 25(2): 1-370. [TreeCo refID: 10436]

GERMANO FILHO, P. 1999. Estudos Taxonômicos do gênero Bathysa C.Presl (Rubiaceae, Rondeletieae), no Brasil.Rodriguésia 50(76/77): 49-75. [TreeCo refID: 10171]

GIBBS, P.E. & SEMIR, J. 2003. A taxonomic revision of the genus Ceiba Mill. (Bombacaceae). Anales Jardín Botánico de Madrid 60: 259-300. [TreeCo refID: 10172]

GOLDENBERG, R. 2004. O gênero Miconia (Melastomataceae) no Estado do Paraná, Brasil. Acta bot. Bras. 18(4): 927-947. [TreeCo refID: 10401]

GOMES, E.P.C.; FISCH, S.T.V. & MANTOVANI, W. 2005. Estrutura e composição do componente arbóreo na Reserva Ecológica do Trabiju, Pindamonhangaba, SP, Brasil. Acta Botanica Brasilica 19(3): 451-464. [TreeCo refID: 997]

GONÇALVES, C.A., LELIS, R.C.C. & ABREU, H.S. 2010. Caracterização físico-química da madeira de sabiá (Mimosa caesalpiniaefolia Benth.).Revista Caatinga 23(1): 54-62. [TreeCo refID: 10041]

GOODMAN, R.C.; PHILLIPS, O.L.; CASTILLO TORRES, D. del.; FREITAS, L.; TAPIA CORTESE, S.; MONTEAGUDO, A. & BAKER, T.R. 2013. Amazon palm biomass and allometry. Forest Ecology and Management 310: 994 - 1004. [TreeCo refID: 10093]

GREEN, P.S. 1994. A Revision of Chionanthus (Oleaceae) in S. America and the Description of Priogymnanthus, gen. nov. Kew Bulletin 49(2): 261-286. [TreeCo refID: 10042]

GROMBONE-GUARATINI, M.T.; GOMES, E.P.C.; TAMASHIRO, J.Y & RODRIGUES, R.R. 2008. Composição florística da Reserva Municipal de Santa Genebra, Campinas, SP. Revista Brasileira de Botânica 31(2): 323-337. [TreeCo refID: 999]

GUIMARÃES, P.J.F. 1997. Estudos taxonômicos de Tibouchina sect. Pleroma (D. Don) Cogn. (Melastomastaceae). Tese (Doutorado). Universidade Estadual de Campinas, São Paulo. 191p. [TreeCo refID: 10043]

GUSSON, A.E.; LOPES, S.F.L.; DIAS NETO, O.C.; VALE, V.S.; OLIVEIRA, A.P. & SCHIAVINI, I. 2009. Características químicas do solo e estrutura de um fragmento de floresta estacional semidecidual em Ipiaçu, Minas Gerais, Brasil. Rodriguésia 60(2): 403-414. [TreeCo refID: 178]

HAIDAR, R.F. 2008. Fitossociologia, diversidade e sua relação com variáveis ambientais em florestas estacionais do Bioma Cerrado no Planalto Central e Nordeste do Brasil. Dissertação (Mestrado). Universidade de Brasília, Brasília, DF, Brasil. 255p. [TreeCo refID: 1256]

HAIDAR, R.F.; FELFILI, J.M.; PINTO, J.R.R. & FAGG, C.W. 2005. Fitossociologia da vegetação arbórea em fragmentos de floresta estacional, no Parque Ecológico Altamiro de Moura Pacheco, GO. Boletim do Herbário Ezechias Paulo Heringer 15: 19-46. [TreeCo refID: 2658]

HAMILTON, C.W. 1989. A revision of mesoamerican Psychotria subgenus Psychotria (Rubiaceae), part I: introduction and species 1-16. Annals of the Missouri Botanical Garden 76(1): 67-111. [TreeCo refID: 10044]

HAYDEN, S.M. & HAYDEN, W.J. 1996. A revision of Discocarpus (Euphorbiaceae). Annals of the Missouri Botanical Garden 83(2): 153-167. [TreeCo refID: 10045]

HERTZOG A.; PELLEGRINI, M.O.O. & SANTOS-SILVA F. 2016. Winteraceae do Rio Grande do Sul, Brasil. Rodriguésia 67(1): 251–260. [TreeCo refID: 10192]

IMAÑA-ENCINAS, J.; PAULA, J.E. & CONCEIÇÃO, C.A. 2012. Florística, volume e biomassa lenhosa de um fragmento de Mata Atlântica no municipio de Santa Maria de Jetibá, Espirito Santo. Floresta 42(3): 565 - 576. [TreeCo refID: 10046]

Inventário Florestal de Minas Gerais (Unpublished data). Literature compilation of species traits provided by the researchers from Minas Gerais state inventory. Universidade Federal de Lavras, Lavras. [TreeCo refID: 10427]

Inventário Florestal de Minas Gerais (Unpublished data). Primary data of maximum tree height and maximum diameter at breast height from the Minas Gerais state inventory. Universidade Federal de Lavras, Lavras. [TreeCo refID: 10426]

ISACKSSON, J.G.L. 2015. Árvores, propágulos e plântulas de duas Licania spp. (Chrysobalanaceae) nativas da floresta de várzea estuarina, Amapá. Monografia (Graduação). Universidade do Estado do Amapá, Macapá. 72p. [TreeCo refID: 10415]

ISACKSSON, J.G.L. 2018. Morfologia de frutos, sementes e plântulas de espécies neotropicais de Chrysobalanaceae como suporte filogenético. Dissertação (Mestrado). Instituto Nacional de Pesquisas da Amazônia, Manaus. 145p. [TreeCo refID: 10412]

IZA, O.B. 2002. Parâmetros de autoecologia de uma comunidade arbórea de Floresta Ombrófila Densa, no Parque Botânico do Morro Baú, Ilhota, SC. Dissertação (Mestrado). Universidade Federal de Santa Catarina, Florianópolis. 92p. [TreeCo refID: 842]

JORDANO, P. 1995. Angiosperm fleshy fruits and seed dispersers: a comparative analysis of adaptation and constraints in plant-animal interactions. American Naturalist 145(2): 163-191. (JORDANO, P. 2013. FRUBASE dataset. Version 3.0. Dryad Digital Repository. https://doi.org/10.5061/dryad.9tb73/1) [TreeCo refID: 10314]

KAASTRA, R.C. 1982. Pilocarpinae (Rutaceae). Flora Neotropica 33: 1-197. [TreeCo refID: 10437]

KILCA, R.V.; SOARES, J.C.W.; MEDEIROS, E.M. & JARENKOW, J.A. 2012. Cambios florísticos y estructurales entre dos comunidades arbóreas de un bosque ripario bajo condiciones ambientales contrastantes en la Pampa sur brasileña. Iheringia série Botânica 67(2):165-175. [TreeCo refID: 1152]

KLEIN, R.M. 1990. Espécies raras ou ameaçadas de Extinção do Estado de Santa Catarina. Vol. 2. IBGE, Rio de Janeiro. 287 p. [TreeCo refID: 10049]

KNAPP, S. 2002. Solanum Section Geminata (Solanaceae). Flora Neotropica 84: 1-404. [TreeCo refID: 10154]

KUBITZKI, K. & RENNER, S. 1982. Lauraceae I (Aniba and Aiouea). Flora Neotropica 31: 1-124. [TreeCo refID: 10134]

KURTZ, B.C. & ARAUJO, D.S.D. 2000. Composição florística e estrutura do componente arbóreo de um trecho de Mata Atlântica na Estação Ecológica Estadual do Paraíso,

Cachoeiras de Macacu, Rio de Janeiro, Brasil. Rodriguesia 51(78/115): 69-112. [TreeCo refID: 531]

LANDRUM, L. 1981. A Monograph of the Genus Myrceugenia (Myrtaceae). Flora Neotropica 29: 1-135. [TreeCo refID: 10141]

LANDRUM, L. 1986. Campomanesia, Pimenta, Blepharocalyx, Legrandia, Acca, Myrrhinium, and Luma (Myrtaceae). Flora Neotropica 45: 1-178. [TreeCo refID: 10155]

LEAL, T.DE.S. 2015. Florística e fitossociologia de Cerrado Sentido Restrito em regeneração natural no município de Pirassununga, estado de São Paulo. Dissertação (Mestrado). Universidade Estadual Paulista "Júlio de Mesquita Filho", Rio Claro. 66p. [TreeCo refID: 3348]

LEITÃO FILHO, H.F. 1972. Contribuição ao conhecimento taxonômico da Tribo Vernonieae no Estado de Sãoo Paulo. Tese (Doutorado). Escola Superior de Agricultura "Luiz de Queiroz", Piracicaba. 217p. [TreeCo refID: 10050]

LERF (Unpublished data). Compilation of species traits from the literature from the Laboratório de Ecologia e Restauração Florestal. ESALQ/USP, Piracicaba. (http://www.lerf.eco.br) [TreeCo refID: 10417]

LIMA JUNIOR, M.J.V. (ed). 2010. Manual de Procedimentos para Análise de Sementes Florestais. Universidade Federal do Amazonas, Manaus. 146p. [TreeCo refID: 10053]

LIMA, A.L.A. & RODAL, M.J.N. 2010. Phenology and wood density of plants growing in the semi-arid region of northeastern Brazil.Journal of Arid Environments 74(11): 1363-1373. [TreeCo refID: 10051]

LIMA, H.C. 1986. Tribo Dalbergieae (Leguminosae - Papilionoideae) em estudo morfológico dos frutos, sementes e plântulas e sua aplicação na sistemática. Dissertação (Mestrado). Universidade Federal do Rio de Janeiro, Rio de Janeiro. 127p. [TreeCo refID: 10052]

LIMA, J.A.S. 2009. Biomassa arbórea e estoques de nutrientes em fragmentos florestais da APA Rio São João: o efeito da fragmentação sobre a Mata Atlântica da Baixada Litorânea Fluminense. Tese (Doutorado). Universidade Estadual do Norte Fluminense, Campo dos Goytacazes. 180p. [TreeCo refID: 3028]

LIMA, M.E.L.; CORDEIRO, I. & MORENO, P.R.H. 2011. Estrutura do componente arbóreo em Floresta Ombrófila Densa Montana no Parque Natural Municipal Nascentes de Paranapiacaba (PNMNP), Santo André, SP, Brasil. Hoehnea 38(1): 73-96. [TreeCo refID: 894]

LINDENMAIER, D.S. & BUDKE, J.C. 2006. Florística, diversidade e distribuição espacial das espécies arbóreas em uma floresta estacional na bacia do rio Jacuí, sul do Brasil. Instituto Anchietano de Pesquisas Botânica 57: 193-216. [TreeCo refID: 638]

LINGNER, D.V.; SCHORN, L.A.; VIBRANS, A.C.; MEYER, L.; SEVEGNANI, L.; GASPER, A.L.; SOBRAL, M.G.; KRÜGER, A.; KLEMZ, G.; SCHMIDT, R. & ANASTÁCIO-JUNIOR, C. 2013. Fitossociologia do componente arbóreo/arbustivo da floresta ombrófila densa em Santa Catarina. In: VIBRANS, A.C.; SEVEGNANI, L.; GASPER, A.L. & LINGNER, D.V. (eds.) Inventário Florístico Florestal de Santa Catarina, Vol. IV, Floresta Ombrófila Densa. Edifurb, Blumenau. 159-200. | MEYER, L.L.; SEVEGNANI, A.L.G.; SCHORN, L.A.; VIBRANS, A.C.; LINGNER, D.V.; SOBRAL, M.; KLEMZ, G.; SCHMITT, R.; ANASTACIO-JR, C. & BROGNI, E. 2013. Fitossociologia do componente arbóreo/arbustivo da Floresta Ombrófila Mista em Santa Catarina. In: VIBRANS, A.C.; SEVGNANI, L.; GASPER, A.L. & LINGNER, D.V. (orgs.) Inventário florístico florestal de Santa Catarina (IFFSC): Floresta Ombrófila Mista. Edifurb, Blumenau. 157-189. | SCHORN, L.A.; LINGNER, D.V.; VIBRANS, A.C.; GASPER, A.L.; SEVEGNANI, L.; SOBRAL, M.; MEYER, L.; KLEMZ, G.; SCHMITT, R.; ANASTACIO-JR, C.; PASQUALLI, V.R. 2013. Estrutura do componente arbóreo/arbustivo da Floresta Estacional Decidual em Santa Catarina. In: VIBRANS, A. C.; SEVGNANI, L.; GASPER, A. L.; LINGNER, D. V. (eds.). Inventário florístico florestal de Santa Catarina (IFFSC): Floresta

Estacional Semidecidual. Edifurb, Blumenau. Vol. 2. 142-163. [TreeCo refID: 2394|2668|2773]

LOBÃO, A.Q. 2009. Filogenia de Guatteria (Annonaceae) e revisão taxonômica das espécies da Floresta Atlântica. Tese (Doutorado). Jardim Botânico do Rio de Janeiro, Rio de Janeiro. 156p. [TreeCo refID: 10054]

LOPES, J.DE.C. & MELLO-SILVA, R. 2012. Annonaceae do Parque Estadual de Ibitipoca, Minas Gerais. Boletim de Botânica 30(2): 157-164. [TreeCo refID: 10055]

LOPES, S.D.F.; SCHIAVINI, I.; PRADO-JÚNIOR, J.A.; GUSSON, A.E.; SOUZA-NETO, A.R.; VALE, V.S. & DIAS-NETO, O.C. 2011. Ecological characterization and diametric distribution of arboreal vegetation in remanescent of seasonal semideciduous forest gloria's experimental farm, Uberlandia, MG. Bioscience Journal 27(2): 322-335. [TreeCo refID: 89]

LOREGIAN, A.C.; SILVA, B.B.; ZANIN, E.M.; DECIAN, V.S.; HENKE-OLIVEIRA, C. & BUDKE, J.C. 2012. Padrões espaciais e ecológicos de espécies arbóreas refletem a estrutura em mosaicos de uma floresta subtropical. Acta Botanica Brasilica 26(3): 593-606. [TreeCo refID: 639]

LORENZI, H. 1992. Árvores brasileiras: manual de identificação e cultivo de plantas arbóreas nativas do Brasil, Vol. 1. Editora Plantarum, Nova Odessa. 384p. [TreeCo refID: 10057]

LORENZI, H. 1998. Árvores brasileiras: manual de identificação e cultivo de plantas arbóreas nativas do Brasil, Vol. 2. Editora Plantarum, Nova Odessa. 351p. [TreeCo refID: 10058]

LORENZI, H. 2009. Árvores brasileiras: manual de identificação e cultivo de plantas arbóreas nativas do Brasil, Vol. 3. Editora Plantarum, Nova Odessa. 384p. [TreeCo refID: 10059]

LORENZI, H.; NOBLICK, L.R.; KAHN, F. & FERREIRA, E. 2010. Flora brasileira: Arecaceae (Palmeiras). Instituto Plantarum, Nova Odessa. 382p. [TreeCo refID: 10056]

MAAS, P.J.M.; WESTRA, L.Y.T. & CHATROU, L.W. 2003. Duguetia (Annonaceae). Flora Neotropica Monograph 88: 1-274. [TreeCo refID: 10060]

MAAS, P.J.M.; WESTRA, L.Y.TH.; BROWN, K.S.; MAAS, P.; TER WELLE, B.J.H.; WEBBER, A.C.; LE THOMAS, A.; WAHA, M.; VAN DER HEIJDEN, E.; BOUMAN, F.; CAVÉ, A.; LEBOEUF, M.; LAPRÉVOTE, O.; KOEK-NOORMAN, J.; MORAWETZ, W. & HEMMER, W. 1992. Rollinia. Flora Neotropica 57: 1-188. [TreeCo refID: 10137]

MAÇANEIRO, J.P.; SEUBERT, R.C. & SCHORN, L.A. 2015. Fitossociologia de uma Floresta Pluvial Subtropical primária no sul do Brasil. Floresta 45(3): 555-566. [TreeCo refID: 3453]

MAGNAGO, L.F.S. (Unpublished data). Measurements and compilations of fruits and seed traits performed by Luiz F. Silva Magnago. Universidade Federal do Sul da Bahia, Itabuna. [TreeCo refID: 10421]

MAGNAGO, L.F.S. 2009. Gradiente vegetacional e pedológico em mata de restinga no estado do Espírito Santo. Dissertação (Mestrado). Universidade Federal de Viçosa, Viçosa. 122p. [TreeCo refID: 24]

MANSANO, V.F. & LIMA, J.R. 2007. O gênero Swartzia Schreb. (Leguminosae, Papilionoideae) no estado do Rio de Janeiro. Rodriguésia 58(2): 469-483. [TreeCo refID: 10183]

MANTOVANI, A.; MORELLATTO, L.P.C. & REIS, M.S. 2004. Fenologia reprodutiva e produção de sementes em Araucaria angustifolia (Bertol.) Kuntze. Revista Brasileira de Botânica 27(4): 787-796. [TreeCo refID: 10407]

MANTOVANI, M.; RUSCHEL, A.R.; PUCHALSKI, Â.; SILVA, J.Z.; REIS, M.S. & NODARI, R.O. 2005. Diversidade de espécies e estrutura sucessional de uma formação secundária da floresta ombrófila densa. Scientia Forestalis 67(1): 14-26. [TreeCo refID: 830]

MANZATTO, A.G.; FURLAN, A.; CESAR, O. & PAGANO, S.N. 1999. Vegetação lenhosa do SESC Interlagos, São Paulo, SP. Universidade Estadual Paulista "Júlio de Mesquista Filho", Rio Claro. 74p. [TreeCo refID: 600]

MARCHIORI, J.N.C. 1997. Dendrologia das angiospermas: das magnoliáceas às flacurtiáceas. Editoria da Universidade Federal de Santa Maria, Santa Maria. 271p. [TreeCo refID: 10127]

MARCHIORI, J.N.C. 1997. Dendrologia das angiospermas: Myrtales. Editoria da Universidade Federal de Santa Maria, Santa Maria. 304p. [TreeCo refID: 10128]

MARCHIORI, J.N.C. 2000. Dendrologia das angiospermas: das bixáceas às rosáceas. Editora da Universidade Federal de Santa Maria, Santa Maria.240p. [TreeCo refID: 10126]

MARCHIORI, N.M.; ROCHA, H.R.; TAMASHIRO, J.Y. & AIDAR, M.P.M. 2016. Composição da comunidade arbórea e biomassa aérea em uma floresta atlântica secundária, Parque Estadual da Serra do Mar, São Paulo, Brazil. CERNE 22(4): 501-514. [TreeCo refID: 3626]

MARÇON, S.L. 2009. Composição florística e estrutura do componente arbustivo-arbóreo do Parque Natural Municipal da Cratera da Colônia, São Paulo, SP. Dissertação (Mestrado). Universidade de São Paulo, Ribeirão Preto. 120p. [TreeCo refID: 1020]

MARCONDES-FERREIRA NETO, W. 1988. Aspidosperma Mart., nom. cons. (Apocynaceae) : estudos taxonomicos. Tese (Doutorado). Universidade Estadual de Campinas, Campinas. 431p. [TreeCo refID: 10402]

MARQUES, S.D.S.; OLIVEIRA, J.D.S.; PAES, J.B.; ALVES, E.S.; SILVA, A. & FIEDLER, N.C. 2012. Estudo comparativo da massa específica aparente e retratibilidade da madeira de pau-brasil (Caesalpinia echinata Lam.) nativa e de reflorestamento. Revista Árvore 36(2): 373-380. [TreeCo refID: 10063]

MARTINEZ-YRIZAR, A.; SARUKHAN, J.; PEREZ-JIMENEZ, A.; RINCON, E.; MAASS, J.M.; SOLIS-MAGALLANES, A. & CERVANTES L. 1992. Above-Ground Phytomass of a Tropical Deciduous Forest on the Coast of Jalisco, Mexico. Journal of Tropical Ecology 8(1): 87-96. [TreeCo refID: 10064]

MARTINI, A.M.Z.; FIASCHI, P.; AMORIM, A.M. & PAIXÃO, J.L. 2007. A hot-point within a hot-spot: a high diversity site in Brazil's Atlantic Forest. Biodiversity and Conservation 16(11) :3111-3128. [TreeCo refID: 10065]

MARTINS, F.R. 1991. Estrutura de uma floresta mesófila. Editora da UNICAMP, Campinas. 214 p. [TreeCo refID: 1022]

MARTINS, L.T. 2012. Caracterização dendrométrica e crescimento de dez espécies florestais nativas em plantios homogêneos no estado do Espírito Santo. Dissertação (Mestrado). Universidade Federal do Espírito Santo, Jerônimo Monteiro. 102p. [TreeCo refID: 10066]

MARTINS, R. 2005. Florística, estrutura fitossociológica e interações interespecíficas de um remanescente de floresta ombrófila densa como subsídio para a recuperação de áreas degradadas pela mineração de carvão, Siderópolis, SC. Dissertação (Mestrado). Universidade Federal de Santa Catarina, Florianópolis. 93p. [TreeCo refID: 2572]

MATOS, M.Q. & FELFILI, J.M. 2010. Florística, fitossociologia e diversidade da vegetação arbórea nas matas de galeria do Parque Nacional de Sete Cidades (PNSC), Piauí, Brasil. Acta Botanica Brasilica 24(2): 483-496. [TreeCo refID: 1589]

MAZINE, F.F. 2002. Estudo taxonômico das espécies de Myrtaceae ocorrentes nos campos de altitude do Parque Nacional do Caparaó (ES/MG). Dissertação (Mestrado). Universidade de São Paulo, São Paulo. 77p. [TreeCo refID: 10067]

MAZINE, F.F. 2006. Estudos taxonomicos em Eugenia L. (Myrtaceae), com enfase em Eugenia secrt. Racemosa O. Berg. Tese (Doutorado). Universidade de São Paulo, São Paulo. 239p. [TreeCo refID: 10068]

MEDEIROS, M.B & WALTER, B.M.T. 2012. Composição e estrutura de comunidades arbóreas de Cerrado sensu stricto no norte do Tocantins e sul do Maranhão. Revista Árvore 36(4): 673-683. [TreeCo refID: 1900]

MEDEIROS, M.B; WALTER, B.M.T. & SILVA, G.P. 2008. Fitossociologia do Cerrado sensu stricto no município de Carolina, Maranhão, Brasil. CERNE 14(4): 285-294. [TreeCo refID: 1894]

MEIFA, M.N.E. & CASTILLO, M.U. 1992. Poder calorífico de cinco espécies de Bombacaceas. Revista Florestal Del Peru 19(1):93-97. [TreeCo refID: 10069]

MEIRELLES, A.C. & SOUZA, L.A.G. 2015. Germinação natural de oito espécies de Swartzia (Fabaceae, Faboideae) da Amazônia. Scientia Amazonia 4(3): 84-92. [TreeCo refID: 10408]

MEIRELLES, J. 2015. Filogenia de Miconia seção Miconia subseção Seriatiflorae e revisão taxonômica do clado Albicans (Melastomataceae, Miconieae). Universidade Estadual de Campinas, Campinas. 219p. [TreeCo refID: 10185]

MELLO-SILVA, R.; LOPES, J.C. & PIRANI, J.R. 2012. Flora da Serra do Cipó, Minas Gerais: Annonaceae. Boletim de Botânica 30(1): 23-35. [TreeCo refID: 10027]

MELO, E. 1996. Levantamento das espécies de Coccoloba (Polygonaceae) da restinga do Estado da Bahia, Brasil. Sitientibus 15: 49-59. [TreeCo refID: 10028]

MELO, E. 1999. Levantamento da família Polygonaceae no estado da Bahia, Brasil: espécies do semi-árido. Rodriguésia 50 (76-77): 29-47. [TreeCo refID: 10030]

MELO, M.M.R.F.; BARROS, F.; CHIEA, S.A.C.; KIRIZAWA, M.; JUNG-MENDAÇOLLI, S.L. & WANDERLEY, M.G.L. (eds.). 2009. Flora Fanerogâmica da Ilha do Cardoso. Vol.14. Instituto de Botânica de São Paulo, São Paulo. 118p. [TreeCo refID: 10131]

MELO, M.M.R.F.; BARROS, F.; CHIEA, S.A.C.; KIRIZAWA, M.; JUNG-MENDAÇOLLI, S.L. & WANDERLEY, M.G.L. (org.). 2008. Flora Fanerogâmica da Ilha do Cardoso. Vol.13. Instituto de Botânica, São Paulo. 143p. [TreeCo refID: 10130]

MENDONÇA FILHO, C.V.; TOZZI, A.M.G.A. & MARTINS, E.R.F. 2007. Revisão taxonômica de Machaerium sect. Oblonga (Benth.) Taub. (Leguminosae, Papilionoideae, Dalbergieae). Rodriguésia 58(2):283-312. [TreeCo refID: 10161]

MENDONÇA, N.T. 2005. Florística e fitossociologia em fragmento de Mata Atlântica – Serra da Bananeira, Estação Ecológica de Murici, Alagoas. Dissertação (Mestrado). Universidade Federal Rural de Pernambuco, Recife. 83p. [TreeCo refID: 2637]

MISSIO, F.F.; SILVA, A.C.; HIGUCHI, P.; LONGHI, S.J.; BRAND, M.A.; RIOS, P.D.; DALLA ROSA, A.; BUZZI JUNIOR, F.; BENTO, M.A.; GONÇALVES, D.A.; LOEBENS, R. & PSCHEIDT, F. 2017 Atributos funcionais de espécies arbóreas em um fragmento de Floresta Ombrófila Mista em Lages, SC. Ciência Florestal 27: 215-224. [TreeCo refID: 10070]

MITCHELL, J.D. & DALY, D.C. 1991. Cyrtocarpa Kunth (Anacardiaceae) in South America. Annals of the Missouri Botanical Garden 78(1): 184-189. [TreeCo refID: 10071]

MORAES, P.L.R. 2007. Taxonomy of Cryptocarya species of Brazil (Vol. 3). Belgian Focal Point to the Global Taxonomy Initiative-Royal Belgian Institute of Natural Sciences. 191p. [TreeCo refID: 10400]

MOREAU, J.S. 2014. Estrutura e interação entre vegetação e ambiente de uma Floresta Ombrófila Densa das Terras Baixas, Espírito Santo. Dissertação (Mestrado). Universidade Federal do Espírito Santo, Jerônimo Monteiro. 96p. [TreeCo refID: 2890]

NASCIMENTO, A.R.T.; FELFILI, J.M. & MEIRELLES, E.M. 2004. Florística e estrutura da comunidade arbórea de um remanescente de Floresta Estacional Decidual de encosta, Monte Alegre, GO, Brasil. Acta Botanica Brasilica 18(3): 659-669. [TreeCo refID: 1251]

NEGRELLE, R.R.B. 2006. Composição florística e estrutura vertical de um trecho de Floresta Ombrófila Densa de planície quaternária. Hoehnea 33(3): 261-289. [TreeCo refID: 831]

NEGRELLE, R.R.B. 2013. Tree Species Composition and Estructure of A Remnant of A Semidecidual Seasonal Alluvial Forest Remnant in Pantanal Matogrossense, Brazil. Revista Arvore 37(6): 989-999. [TreeCo refID: 3079]

NÓBREGA, M.G.G.; RAMOS, A.E. & SILVA JUNIOR, M.C. 2001. Composição florísitica e estrutura na mata de galeria do Cabeça de Veado, no Jardim Botânico de Brasília, FDF. Boletim do Herbário Ezechias Paulo Heringer 8: 44-65. [TreeCo refID: 1264]

NOGUEIRA JÚNIOR, L.R. 2010.Estoque de carbono na fitomassa e mudanças nos atributos do solo em diferentes modelos de restauração da Mata Atlântica. Tese (Doutorado). Escola Superior de Agricultura "Luiz de Queiroz", Piracicaba. 94 p. [TreeCo refID: 10072]

NOGUEIRA, E.M. 2008. Wood density and tree allometry in forests of Brazil's 'arc of deforestation' implications for biomass and emission of carbon from land-use change in Brazilian Amazonia. Tese (Doutorado). INPA/UFAM, Manaus. 130p. [TreeCo refID: 10073]

NUNES, E.S.; GUIMARÃES JÚNIOR, J.B.; OLIVEIRA, R.J. & GUIMARÃES NETO, R.M. 2012. determinação da densidade básica da madeira de Qualea parviflora Mart. e Qualea grandiflora Mart.(pau-terra) para produção de carvão vegetal. XXI Seminário de Iniciação Científica IV Seminário em Desenvolvimento Tecnológico e Inovação/ Teresina (PI), 24 a 26 de Outubro de 2012. Resumo expandido. ISSN: 1518-7772. [TreeCo refID: 10074]

OLIVEIRA-FILHO, A.T. 2017. NeoTropTree, Flora arbórea da Região Neotropical: Um banco de dados envolvendo biogeografia, diversidade e conservação. Universidade Federal de Minas Gerais. <http://www.neotroptree,info> Acessado: abril, 2019. [TreeCo refID: 10156]

OLIVEIRA-FILHO, A.T. TreeAtlan 2.0, Flora arbórea da América do Sul cisandina tropical e subtropical: Um banco de dados envolvendo biogeografia, diversidade e conservação. Universidade Federal de Minas Gerais. Disponivel em: <http://www.icb.ufmg.br/treeatlan/>. Acessodo: 2013. [TreeCo refID: 10190]

OLIVEIRA-FILHO, A.T.; CARVALHO, D.A.; FONTES, M.A.L.; Van Den BERG, E.; CURI, N. & CARVALHO, W.A.C. 2004. Variações estruturais do compartimento arbóreo de uma floresta semidecídua alto-montana na chapada das Perdizes, Carrancas, MG. Revista Brasileira de Botânica 27(2): 291-309. [TreeCo refID: 219]

OLIVEIRA-FILHO, A.T.; VILELA, E.A.; GAVILANES, M.L. & CARVALHO, D.A. 1994e. Effect of flooding regime and understorey bamboos on the physiognomy and tree species composition of a tropical semideciduous forest in Southeastern Brazil. Vegetatio 113(2): 99-124. [TreeCo refID: 280]

OLIVEIRA, A.M. 2011. Caracterizaçãoo de uma comunidade de árvores e sua infestação por lianas em uma floresta decídua. Dissertação (Mestrado). Universidade Estadual Paulista "Júlio de Mesquita Filho", Botucatu. 99p. [TreeCo refID: 10075]

OLIVEIRA, G.M.V. 2014. Densidade da madeira em Minas Gerais: amostragem, espacialização e relação com variáveis ambientais. Tese (Doutorado). Universidade Federal de Lavras, Lavras. 125p. [TreeCo refID: 10076]

OLIVEIRA, M.M.A. 1999. Frugivoria por aves em um fragmento de floresta de restinga no estado do Espírito Santo, Brasil. Tese (Doutorado), Universidade Estadual de Campinas, Campinas. 153p. [TreeCo refID: 10077]

PASETTO, M.R. 2008. Composição florística e estrutura de fragmento de Floresta Ombrófila Densa Submontana no município de Siderópolis, Santa Catarina. Trabalho de Conclusão de Curso (Graduação). Universidade do Extremo Sul Catarinense - UNESC, Criciúma. 44p. [TreeCo refID: 2715]

PAULA, A. & SOARES, J.J. 2010. Estrutura horizontal de um trecho de floresta ombrófila densa das terras baixas na Reserva Biológica de Sooretama, Linhares, ES. Floresta 41(2): 321-334. [TreeCo refID: 8]

PAULA, A; SILVA, A.F.; MARCO-JÚNIOR, P.; SANTOS, F.A.M. & SOUZA, A.L. 2004. Sucessão ecológica da vegetação arbórea em uma Floresta Estacional Semidecidual, Viçosa, MG, Brasil. Acta Botanica Brasilica 18(3): 401-699. [TreeCo refID: 224]

PAULA, J. E.; IMAÑA-ENCINAS, J. & SUGIMOTO, N. 1998. Levantamento quantitativo em três hectares de vegetação de cerrado. Pesquisa Agropecuária Brasileira 33(5): 613-620. [TreeCo refID: 2410]

PAULA, J.E. & ALVES, J.L.H. 2010. 922 Madeiras nativas do Brasil: anatomia, dendrologia, dendrometria, produção e uso. Ed. Cinco Continentes, Porto Alegre. 461p. [TreeCo refID: 10078]

PAULA, J.E., IMAÑA-ENCIMAS, J.; PEREIRA, B.A.S. 1993. Inventário de um hectare de Mata Ripária. Pesquisa Agropecuária Brasileira 28(2): 143-152. [TreeCo refID: 2411]

PAULA, J.E.; IMAÑA-ENCINAS, J. & PEREIRA, B.A.S. 1996. Parâmetros volumétricos e da biomassa da mata ripária do Córrego dos Macacos. CERNE 2(2): 91-105. [TreeCo refID: 1882]

PEDREIRA, G. & SOUSA, H.C. 2011. Comunidade arbórea de uma mancha florestal permanentemente alagada e de sua vegetação adjacente em Ouro Preto-MG, Brasil. Ciência Floresta 21(4): 663-675. [TreeCo refID: 103]

PEIXOTO, A.L. 1987. Revisão Taxonomica do Genero Mollinedia Ruiz et Pavon (Monimiaceae, Monimioideae). Universidade Estadual de Campinas, Campinas. 401p. [TreeCo refID: 10079]

PENNINGTON, T. 1990. Sapotaceae. Flora Neotropica 52: 1-770. [TreeCo refID: 10136]

PENNINGTON, T., STYLES, B. & TAYLOR, D.A.H. 1981. Meliaceae, with Accounts of Swietenioideae and Chemotaxonomy. Flora Neotropica 28: 1-470. [TreeCo refID: 10133]

PENNINGTON, T.D. 1990. Sapotaceae. Flora Neotropica Monograph 52: 1-770. [TreeCo refID: 10080]

PENNINGTON, T.D. 1997. The genus Inga: botany. The Royal Botanical Garden, Kew. 844p. [TreeCo refID: 10081]

PENNINGTON, T.R. 2003. Monograph of Andira (Leguminosae-Papilionoideae). Systematic Botany Monographs 64: 1-143. [TreeCo refID: 10123]

PERDIZ, R.O.; FERRUCCI, M.S. & AMORIM, A.M.A. 2014. Sapindaceae em remanescentes de florestas montanas no sul da Bahia, Brasil. Rodriguésia 65(4): 987-1002. [TreeCo refID: 10162]

PEREIRA, M.S. 2007. O genêro Coussarea Aubl. (Rubiaceae, Rubioideae, Coussareae) na Mata Atlântica. Tese (Doutorado). Universidade Federal de Pernambuco, Recife. 136p. [TreeCo refID: 10175]

PEREIRA, Z.V. & KINOSHITA, L.S. 2013. Rubiaceae Juss. of Parque Estadual das Várzeas do Rio Ivinhema, Mato Grosso do Sul State, Brazil. Hoehnea 40(2): 205-251. [TreeCo refID: 10082]

PILLAR, V.D. & SOSINSKI, E. 2003. An improved method for searching plant functional types by numerical analysis. Journal of Vegetation Science 14: 323-332. (TRY dataset 77: FAPESP Brazil Rainforest Database). [TreeCo refID: 10425]

PIÑA- RODRIGUES, F.C.M.; FREIRE, J.M.; LELES, P.S.S.; BREIER, T.B. (Org.). Parâmetros técnicos para produção de sementes florestais. Editora da UFRRJ, Seropédica. 188p. [TreeCo refID: 10083]

PINHEIRO, K.; ALVES, M. 2007 Espécies arbóreas de uma área de Caatinga no sertão de Pernambuco, Brasil: dados preliminares. Revista Brasileira de Biociências 5(sup): 426-428. [TreeCo refID: 3506]

PIRANI, J.R. 1987. Flora da Serra do Cipó: Burseraceae. Boletim de Botânica 9: 211-218. [TreeCo refID: 10084]

PIRANI, J.R. 1998. A revision of Helietta and Balfourodendron (Rutaceae-Pteleinae). Brittonia 50(3): 348-380. [TreeCo refID: 10177]

POSSETTE, R.F.DA.S. & RODRIGUES, W.A. 2010. O gênero Inga Mill. (Leguminosae - Mimosoideae) no estado do Paraná, Brasil. Acta Bot. Bras. 24(2): 354-368. [TreeCo refID: 10180]

PRADO-JÚNIOR, J.A.; LOPES, S.F.; VALE, V.S.; OLIVEIRA, A.P.; GUSSON, A.E.; DIAS-NETO, O.C. & SCHIAVINI, I. 2011. Estrutura e caracterização sucessional da comunidade arbórea de um remanescente de floresta estacional semidecidual, Uberlândia, MG. Caminhos de Geografia 12(39): 81-93. [TreeCo refID: 235]

PRADO-JÚNIOR, J.A.P.; VALE, V.S.; OLIVEIRA, A.P.; GUSSON, A.E.; DIAS-NETO, O. C.; LOPES, S.F. & SCHIAVINI, I. 2010. Estrutura da comunidade arbórea em um fragmento de Floresta Estacional Semidecidual localizada na Reserva Legal da Fazenda Irara, Uberlândia, MG. Bioscience Journal 26(4): 638-647. [TreeCo refID: 100]

PRADO JUNIOR, J.A.; FARIA, S.; SCHIAVINI, I.; VALE, V.; OLIVEIRA, A. P.; GUSSON, A. E.; DIAS, N.; OLAVO, C. & STEIN, M. 2012. Fitossociologia, caracterização sucessional e síndromes de dispersão da comunidade arbórea de remanescente urbano de Floresta Estacional Semidecidual em Monte Carmelo, Minas Gerais. Rodriguésia 63(3): 489-499. [TreeCo refID: 1307]

PRANCE, G. & MORI, S. 1979. Lecythidaceae: Part I: The Actinomorphic-Flowered New World Lecythidaceae (Asteranthos, Gustavia, Grias, Allantoma, & Cariniana). Flora Neotropica 21(1): 1-270. [TreeCo refID: 10140]

PRANCE, G.T.; PLANA, V.; EDWARDS, K.S. & PENNINGTON, R.T. 2007. Proteaceae. Flora Neotropica 100: 1-218. [TreeCo refID: 10434]

PROENÇA, C. 1986. Revisão de Siphoneugena (Myrtaceae, Myrteae). Dissertação (Mestrado). Universidade Federal do Rio de Janeiro, Rio de Janeiro. 163p. [TreeCo refID: 10159]

QUIRINO, W.F.; VALE, A.T.; ANDRADE, A.P.A.; ABREU, V.L.S. & AZEVEDO, A.D.S. 2005. Poder calorífico da madeira e de materiais ligno-celulósicos.Revista da Madeira 89: 100-106. [TreeCo refID: 10085]

Rede speciesLink. Data retrieved for record descriptions or images from the speciesLink netwrok. http://www.splink.org.br. [TreeCo refID: 10428]

REIS, A. 1993. Manejo e conservação das florestas catarinenses (Trabalho apresentado para o Concurso de Professor Titular de Botânica Aplicada). Universidade Federal de Santa Catarina, Florianópolis. [TreeCo refID: 10086]

REITZ, P.R. 1965. Plano de coleção. In: REITZ, R. (ed). Flora Ilustrada Catarinense. Herbário Barbosa Rodrigues, Itajaí. 71p. [TreeCo refID: 10087]

REITZ, R. (ED.). 1965 - 1985. Flora Ilustrada Catarinense. Herbário Barbosa Rodrigues, Itajaí. [TreeCo refID: 10176]

REITZ, R.; KLEIN, R.M. & REIS, A. 1979. Madeiras do Brasil - Santa Catarina. REITZ, R. (Ed.). Editora Lunardelli, Florianópolis. 320 p.|REITZ,R.; KLEIN, R.M. & REIS, A. 1983. Projeto Madeira do Rio Grande do Sul. Sellowia 34-35. 525p. [TreeCo refID: 10116]

RENNER, S. & HAUSNER, G. 2005. Siparunaceae. Flora Neotropica 95: 1-247. [TreeCo refID: 10433]

RIBEIRO, B.R. 2014. Onde estão os recrutas? A matriz de pasto pode acelerar alterações na assembléia de árvores grandes em áreas fragmentadas. Monografia (Graduação). Universidade Federal de Alfenas, Alfenas. 39p. [TreeCo refID: 3851]

RIBEIRO, S.C.; FEHRMANN, L.; SOARES, C.P.B.; JACOVINE, L.A.G.; KLEINN, C. & GASPAR, R.O. 2011. Above-and belowground biomass in a Brazilian Cerrado. Forest Ecology and Management 262(3): 491-499. [TreeCo refID: 10088]

ROCHA, D.S.B & AMORIM, A.M.A. 2012. Heterogeneidade altitudinal na Floresta Atlântica setentrional: um estudo de caso no sul da Bahia, Brasil. Acta Botanica Brasilica 26(2): 309-327. [TreeCo refID: 10089]

RODRIGUES, A.V.; BONES, F.L.V.; SCHNEIDERS, A.; OLIVEIRA, L.Z.; VIBRANS, A.C. & GASPER, A.L. 2018. Plant trait dataset for tree-like growth forms species of the subtropical Atlantic Rain Forest in Brazil. Data 3(2): 16. [TreeCo refID: 10090]

RODRIGUES, I.A. 1982. Contribuição à sistemática das espécies do gênero Inga P. Mill. (Leg. Mim.) ocorrentes no Estado do Rio de Janeiro. Dissertação Mestrado, UFRJ, Rio de Janeiro. [TreeCo refID: 10092]

RODRIGUES, I.M.C. & GARCIA, F.C.P. 2007. Papilionoideae (Leguminosae) arbóreas e lianas na estacão de pesquisa, treinamento e educação ambiental (EPTEA), Mata do Paraíso, Viçosa, Zona da Mata Mineira. Revista Árvore 31(3):521-532. [TreeCo refID: 10091]

RODRIGUES, L.A.; CARVALHO, D.A.; OLIVEIRA-FILHO, A.T.; BOTREL, R.T. & SILVA, E.A. 2003. Florística e estrutura da comunidade arbórea de um fragmento florestal em Luminárias, MG. Acta Botanica Brasilica 17(1): 71-87. [TreeCo refID: 75]

RODRIGUES, R.R.; GANDOLFI, S. & SOUZA, V.C. 2006. Diversidade, dinâmica e conservação em florestas do estado de São Paulo: 40,96ha de parcelas permanentes. Universidade de São Paulo, Piracicaba, Brazil, 68p. [TreeCo refID: 936]

RODRIGUES, V.H.P.; LOPES, S.F.; ARAÚJO, G.M. & SCHIAVINI, I. 2010. Composição, estrutura e aspectos ecológicos da floresta ciliar do rio Araguari no Triângulo Mineiro. Hoehnea 37(1): 87-105. [TreeCo refID: 76]

ROHWER, J. 1993. Lauraceae: Nectandra. Flora Neotropica 60: 1-332. [TreeCo refID: 10142]

ROSEIRA, D.S. 1990. Composição florística e estrutura fitossociológica do bosque com Araucaria angustifolia (Bert.) Kuntze no Parque Estadual João Paulo II, Curitiba, Paraná. Dissertação (Mestrado). Universidade Federal do Paraná, Curitiba. 111p. [TreeCo refID: 411]

ROYAL BOTANIC GARDENS KEW. 2017. Seed Information Database (SID). Version 7.1. <http://data.kew.org/sid/> Accessed on: March 2017. [TreeCo refID: 10094]

SÁ, C.F.C. & ARAUJO, D.S.D. 2009. Estrutura e florística de uma floresta de Restinga em Ipitangas, Saquarema, Rio de Janeiro, Brasil. Rodriguésia 60: 147-170. [TreeCo refID: 2848]

SALES, H.R.; SOUZA, S.C.A.; LUZ, G.R.; MORAIS-COSTA, F.; AMARAL, V.B.; SANTOS, R.M.; VELOSO, M.D.M. & NUNES, Y.R.F. 2009. Flora arbórea de uma Floresta Estacional Decidual na APA Estadual do Rio Pandeiros, Januária, MG. Biota 2(3): 31-41. [TreeCo refID: 244]

SALIS, S.M.; LEHN, C.R.; PADILHA, D.R.C. & MATTOS, P.P. 2012. Changes in the structure due to strong winds in forest areas in the Pantanal, Brazil. CERNE 18(3): 387-395. [TreeCo refID: 2350]

SALOMÃO, N.A.; DAVIDE, A.C.; FIRETTI, F.; SOUSA E SILVA, F.C.; CALDAS, L.S.; WETZEL, M.M.V.S.; TORRES, R.A.A. & GONZÁLES, S. 2003. Germinação de sementes e produção de mudas de plantas do Cerrado. Rede de Sementes do Cerrado, Brasília. 93p. [TreeCo refID: 10095]

SALYWON, A.M. & LANDRUM, L.R. 2007. Curitiba (Myrtaceae): A New Genus from the Planalto of Southern Brazil. Brittonia 59(4): 301-307. [TreeCo refID: 10096]

SAMPAIO, A.B.; WALTER, B.M.T. & FELFILI, J.M. 2000. Diversidade e distribuição de espécies arbóreas em duas matas de galeria do riacho Fundo, Distrito Federal. Acta Botanica Brasilisa 14(2): 197-214. [TreeCo refID: 1268]

SAMPAIO, D. 2009. Revisão taxonômica das espécies neotropicais extra-amazônicas de Sloanea L. (Elaeocarpaceae) na América do Sul. Tese (Doutorado). Universidade Estadual de Campinas, Campinas. 168p. [TreeCo refID: 10165]

SANTANA, G.C. 2010. Estrutura de uma floresta ombrófila densa montana com monodominância de dossel por Eremanthus erythropappus (DC.) Macleish (candeia) na serra da Mantiqueira, em Itamonte, Minas Gerais. Dissertação (Mestrado). Universidade Federal de Lavras, Lavras. 58p. [TreeCo refID: 245]

SANTIN, D.A. 1989. Revisão taxonômica do gênero Astronium Jacq. e revalidação do gênero Myracrodruon Fr. Allem. (Anacardiaceae). Dissertação de mestrado, Universidade Estadual de Campinas, Campinas. 187p. [TreeCo refID: 10160]

SANTOS, I.S. & PEIXOTO, A.L. 2001. Taxonomia do gênero Macropeplus Perkins (Monimiaceae, Monimioideae). Rodriguesia 52(81): 65-105. [TreeCo refID: 10097]

SANTOS, K. 2003. Caracterização florística e estrutural de onze fragmentos de mata estacional semidecidual da área de proteção ambiental do município de Campinas – SP. Tese (Doutorado). Universidade de Campinas, Campinas. 225p. [TreeCo refID: 2809]

SARTORI, Â.L.B.; LEWIS, G.P.; MANSANO, V.F. & TOZZI, A.M.G.A. 2015. A revision of the genus Myroxylon (Leguminosae: Papilionoideae). Kew Bulletin 70(4): 48. [TreeCo refID: 10098]

SCHORN, L.A. 2005. Estrutura e dinâmica de estágios sucessionais de uma floresta ombrófila densa em Blumenau, Santa Catarina. Tese (Doutorado). Universidade Federal do Paraná, Curitiba. 192p. [TreeCo refID: 855]

SCHORN, L.A.; GASPER, A.L. de; MEYER, L.; VIBRANS, A.C. 2012. Síntese da estrutura dos remanescentes florestais em Santa Catarina. In: Vibrans, A.C., Sevegnani, L.; Gasper A.L.; Lingner, D.V. (Org.). 2012. Inventário Florístico Florestal de Santa Catarina, Vol. 1, Diversidade e Conservação dos remanescentes florestais. Edifurb, Blumenau, 1 ed: 125-140 (Classificação consensuada utilizada pela FURB a partir de 2011 para o IFSC) [TreeCo refID: 10125]

SECCO, R.S. 2004. Alchorneae (Euphorbiaceae) (Alchornea, Aparisthmium e Conceveiba). Flora Neotropica 93: 1-194. [TreeCo refID: 10432]

SECRETARIA DO MEIO AMBIENTE DO ESTADO DE SÃO PAULO. 1991. Anais do 2° Simpósio Brasileiro sobre Tecnologia de Sementes Florestais. Atibaia, São Paulo, 16 a 19 de outubro de 1989. Instituto Florestal, São Paulo. 319p. [TreeCo refID: 10100]

SENA, C.M. & GARIGLIO, M.A. 2008. Sementes Florestais: Colheita, Beneficiamento e Armazenamento. MMA, Secretaria de Biodiversidade e Florestas, Departamento de Florestas, Programa Nacional de Florestas, Natal. 28p. [TreeCo refID: 10101]

SENA, L.H.M. 2014. Conservação de sementes e produção de mudas de pitombeira (Talisia esculenta (A. St. Hil.) Radlk.). Dissertação (Mestrado). Universidade Federal Rural de Pernambuco, Recife. 122p. [TreeCo refID: 10414]

SENNA, L.M. 1984. Maprounea Aubl. (Euphorbiaceae). Considerações taxonômicas e anatômicas das espécies sul-americanas. Rodriguesia 36(61): 51-78. [TreeCo refID: 10164]

SEVEGNANI, L. (Unpublished data) Information on species traits compiled by Lucia Sevegnani. FURB, Blumenau. [TreeCo refID: 10422]

SEVILHA, A.C.; PAULA, A; LOPES, W.P. & SILVA, A.F. 2001. Fitosociologia de estrato arbóreo de um trecho de Floresta Estacional no Jardim Botânico da Universidade Federal de Viçosa (face sudoeste), Viçosa, Minas Gerais. Revista Árvore 25(4): 431-443. [TreeCo refID: 256]

SILVA-JÚNIOR, J.F. 2004. Estudo fitossociológico em um remanescente de floresta atlântica visando dinâmica de espécies florestais arbóreas no município do Cabo de Santo Agostinho, PE. Dissertação (Mestrado). Universidade Federal Rural de Pernambuco, Recife. 74p. [TreeCo refID: 1570]

SILVA JÚNIOR, M.C.DA. & PEREIRA, B.A.S. 2009. 100 Arvores do Cerrado-Matas de Galeria. Editora Rede de Sementes do Cerrado, Brasilia. 288p. [TreeCo refID: 10061]

SILVA JÚNIOR, M.C.DA. 2005. 100 Árvores do cerrado: guia de campo. Editora Rede de Sementes do Cerrado, Brasilia. 278p. [TreeCo refID: 10001]

SILVA, A.C.; HIGUCHI, P.; AGUIAR, M.D.; NEGRINI, M.; FERT NETO, J. & HESS, A.F. 2012. Relações florísticas e fitossociologia de uma floresta ombrófila mista montana secundária em Lages, Santa Catarina. Ciência Florestal 22(1): 193-206. [TreeCo refID: 828]

SILVA, A.C.D.; VAN DEN BERG, E.; HIGUCHI, P.; OLIVEIRA-FILHO, A.T.; MARQUES, J.J.G.D.S.; APPOLINÁRIO, V.; PIFANO, D.S.; OGOSUKU, L.M. & NUNENS, M. 2009. Tree community floristic and structure of alluvial forest fragments in São Sebastião da Bela Vista, Minas Gerais, Brazil. Brazilian Journal of Botany 32(2): 283-297. [TreeCo refID: 60]

SILVA, C.V. & REIS, M.S. 2009. Produção de pinhão na região de Caçador, SC: aspectos da obtenção e sua importância para comunidades locais. Ciência Florestal 19(4): 363-374. [TreeCo refID: 10406]

SILVA, M.J. & TOZZI, A.M.G.A. 2012. Revisão taxonômica de Lonchocarpus s. str. (Leguminosae, Papilionoideae) do Brasil. Acta Botanica Brasilica 26(2): 357-377. [TreeCo refID: 10104]

SILVA, M.S.; SANTOS, F.A.R.; SILVA, C.R.A. & SILVA, L.B. 2012. Características das fibras e densidade básica da madeira de quatro espécies de Mata Atlântica (Serra da Jibóia, Elísio

Medrado, Bahia, Brasil): qualificação para uso e preservação. I Simpósio sobre a Biodiversidade da Mata Atlântica: 129-133. [TreeCo refID: 10062]

SILVA, R.K.S. 2009. Fitossociologia do componente arbóreo em áreas ciliares e de nascentes de um fragmento de floresta ombrófila densa de terras baixas, em Sirinhaém, Pernambuco. Dissertação (Mestrado). Universidade Federal Rural de Pernambuco, Recife. 80p. [TreeCo refID: 2740]

SILVEIRA, N.M.; ALVES, J.D.; DOUSSEAU, S. & ALVARENGA, A.A. 2013. Technology seed Sebastiania membranifolia Mull Arg (Euphorbiaceae). Cerne 19(4): 669-75. [TreeCo refID: 10410]

SILVEIRA, P. 2008. Métodos indiretos de estimativa do conteúdo de biomassa e do estoque de carbono em um fragmento de floresta ombrófila densa. Tese (Doutorado). Universidade Federal do Paraná, Curitiba. 129p. [TreeCo refID: 10105]

SLEUMER, H.O. 1980. Flacourtiaceae. Flora Neotropica 22: 1-499. [TreeCo refID: 10132]

SLEUMER, H.O. 1984. Olacaceae. Flora Neotropica 38: 1-158. [TreeCo refID: 10438]

SLUSARSKI, S.R. & SOUZA, M.C. 2012. Analysis of floristic similarity between forest remnants from the upper Paraná river floodplain, Brazi. Acta Scientiarum, Biological Sciences 34(3): 343-352. [TreeCo refID: 1159]

SOBRAL, M. 2011. Eugenia (Myrtaceae) no Parana. Eduel, Londrina. 236 p. [TreeCo refID: 10166]

SOSINSKI, E.; JOLY, C.A. & PILLAR, V.D. (Unpublished data). TRY dataset 77: FAPESP Brazil Rainforest Database. [TreeCo refID: 10424]

SOUSA-JÚNIOR, P.R.C. 2006. Estrutura da comunidade arbórea e da regeneração natural em um fragmento de floresta urbana, Recife - PE. Dissertação (Mestrado). UFRPE, Recife. 91p. [TreeCo refID: 2636]

SOUZA, A.L.; BOINA, A.; SOARES, C.P.B.; VITAL, B.R.; GASPAR, R.O. & LANA, J.M. 2012. Estrutura fitossociológica, estoques de volume, biomassa, carbono e dióxido de carbono em floresta estacional semidecidual. Revista Árvore 36(1): 169-179. [TreeCo refID: 97]

SOUZA, I.M.; FUNCH, L.S. & QUEIROZ, L.P 2016. Flora da Bahia: Leguminosae – Hymenaea (Caesalpinioideae: Detarieae). Sitientibus, série Ciências Biológicas 16: http://dx.doi.org/10.13102/scb1092 [TreeCo refID: 10179]

SOUZA, J.S.; ESPÍRITO-SANTO, F.D.B.; FONTES, M.A.L.; OLIVEIRA-FILHO, A.T. & BOTEZELLI, L. 2003. Análise das variações florísticas e estruturais da comunidade arbórea de um fragmento de floresta semidecídua às margens do rio Capivari, Lavras-MG. Revista Árvore 27(2): 185-206. [TreeCo refID: 82]

SOUZA, M.C. 2009. Estudos taxônomicos em Myrtaceae no Brasil: Revisão de Neomitranthes Kausel ex D.Legrand e contribuição ao conhecimento da diversidade e conservação de Plinia L. (Myrtaceae Juss.) no Dominío Atlântico. Tese (Doutorado). Jardim Botânico do Rio de Janeiro, Rio de Janeiro. [TreeCo refID: 10107]

SPINA, A.P. 2004. Estudos taxonomico, micro-morfologico e filogenetico do genero Himatanthus Willd. ex Schult. (Apocynaceae: Rauvolfioideae - Plumerieae). Tese (Doutorado). Universidade Estadual de Campinas, Campinas. 191p. [TreeCo refID: 10178]

SÜHS, R.B.; PUTZKE, J. & BUDKE, J.C. 2010. Relações florístico-geográficas na estrutura de uma floresta na região central do Rio Grande do Sul, Brasil. Floresta 40(3): 635-646. [TreeCo refID: 741]

TABANEZ, A.A.J.; VIANA, V.M. & DIAS, A.S. 1997. Conseqüências da fragmentação e do efeito de borda sobre a estrutura, diversidade e sustentabilidade de um fragmento de floresta de Planalto de Piracicaba, SP. Revista Brasileira de Biologia 57(1): 47-60. [TreeCo refID: 1077]

TAMASHIRO, J.Y. 1989. Estudos taxonomicos e morfologicos do genero Piptadenia sensu Dentham no sudoeste do Brasil: avaliação das modificações taxonomicas recentemente propostas. Dissertação (Mestrado). Universidade Estadual de Campinas, Campinas. 99p. [TreeCo refID: 10188]

TAVARES, R.P. 2010. Morfoanatomia foliar de espécies de Brunfelsia L. do Sul do Brasil. Dissertação (Mestrado). Universidade Federal de Santa Catarina, Florianópolis. 76p. [TreeCo refID: 10191]

THOMAS, W.W.; CARVALHO, A.M.V.; AMORIM, A.M.; HANKS, J.G. & SANTOS, T.S. 2008. Diversity of Woody Plants in the Atlantic Coastal Forest of Southern Bahia, Brazil. In: W.W. THOMAS (ed.). The Atlantic Coastal Forest of Northeastern Brazil. Memoirs of the New York Botanical Garden 100: 21-66. [TreeCo refID: 10109]

TOMASETTO, F. 2003. Composição florística e estrutura do componente arbóreo de um trecho de floresta estacional semidecidual na Estação Ecológica de Paulo de Faria – SP. Dissertação (Mestrado). Universidade Estadual Paulista, Rio Claro. 133 p. [TreeCo refID: 1344]

TONIATO, M.T.Z.; OLIVEIRA-FILHO, A.T. 2004. Variations in tree community composition and structure in a fragment of tropical semideciduous forest in southeastern Brazil related to different human disturbance histories. Forest Ecology and Management 198(1): 319-339. [TreeCo refID: 889]

TOZZI, A.M.G.A. 1989. Estudos taxônomicos dos gêneros Lonchocarpus Kunth e Deguelia Aubl. no Brasil. Tese (Doutorado). Universidade Estadual de Campinas, Campinas. 341p. [TreeCo refID: 10110]

TOZZI, A.M.G.A.; MELHEM, T.S.; FORERO, E.; FORTUNA-PEREZ, A.P.; WANDERLEY, M.G.L.; MARTINS, S.E.; ROMANINI, R.P.; PIRANI, J. R.; MELO, M.M.R.F.DE.; KIRIZAWA, M.; YANO, O. & CORDEIRO, I (eds.). 2016. Flora Fanerogâmica do Estado de São Paulo, Vol. VIII, Leguminosae. Instituto de Botânica, São Paulo. 441p. [TreeCo refID: 10036]

VALE, A.T. (Unpublished data). Measurements of wood specific gravity made by Ailton T. Vale for Cerrado species. Universidade de Brasília, Brasília. [TreeCo refID: 10419]

VALE, A.T. 2000. Caracterização da biomassa lenhosa de um cerrado sensu stricto da região de Brasília para uso energético. Tese (Doutorado). Universidade Estadual de São Paulo "Júlio de Mesquita Filho", Botucatu. 111p. | VALE, A.T.; BRASIL, M.A.M.; LEÃO, A.L. 2002. Quantificação e caracterização energética da madeira e casca de espécies do cerrado. Ciência Florestal, 12(1):71-80. | VALE, A.T. & FELFILI, J.M. 2005. Dry biomass distribution in a cerrado sensu stricto site in Brazil central. Revista Árvore, 29(5): 661-669. [TreeCo refID: 10119]

VALE, A.T.; DIAS, Í.S. & SANTANA, M.A.E. 2010. Relações entre propriedades químicas, físicas e energéticas da madeira de cinco espécies de cerrado. Ciência Florestal 20(1): 137-145. [TreeCo refID: 10111]

VAZ, A.M.S.F. & TOZZI, A.M.G.A 2003. Bauhinia ser. Cansenia (Leguminosae: Caesalpinioideae) no Brasil. Rodriguésia 54(83): 55-143. [TreeCo refID: 10122]

VEIGA, L.G. 2010. Estoque de madeira morta ao longo de um gradiente altitudinal de Mata Atlântica no nordeste do estado de São Paulo. Dissertação (Mestrado). Universidade Estadual de Campinas, Campinas. 71p. [TreeCo refID: 10112]

VENZKE, T.S.L. 2012. Florística, estrutura e síndrome de dispersão de sementes em estágios sucessionais de mata ciliar no município de Arroio do Padre, RS, Brasil. Dissertação (mestrado). Universidade Federal de Viçosa, Viçosa, 82p. [TreeCo refID: 798]

VIANI, R.A.G.; COSTA, J.C.; ROZZA, A.F.; BUFO, L.B.V.; FERREIRA, M.A.P. & OLIVEIRA, A.C.P. 2011. Caracterização florística e estrutural de remanescentes florestais de Quedas do Iguaçu, Sudoeste do Paraná. Biota Neotropica 11(1): 115-128. [TreeCo refID: 360]

VIGNOLI-SILVA, M. 2009. O gênero Cestrum L. (Solanaceae) no Brasil extra-amazônico. Tese (Doutorado). Universidade Federal do Rio Grande do Sul, Porto Alegre. 317p. [TreeCo refID: 10173]

VILELA, E.A.; OLIVEIRA-FILHO, A.T.; CARVALHO, D.A. & CURI. N. 1998. Estudos florísticos e fitossociológicos em remanescentes de florestas ripárias do Baixo Rio Paranaíba e Alto Rio São Francisco. Boletim técnico 01000-GE/PA-1, Companhia Energética de Minas Gerais (CEMIG) , Belo Horizonte. 23p. [TreeCo refID: 274]

WANDERLEY, M.G.L.; SHEPHERD, G.J. & GIULIETTI, A.M. (coords.). 2002. Flora Fanerogâmica do Estado de São Paulo. Vol. 2. Ed. Hucitec, São Paulo. 386p. [TreeCo refID: 10144]

WANDERLEY, M.G.L.; SHEPHERD, G.J.; GIULIETTI, A.M. & MELHEM, T.S. (coords.). 2003. Flora Fanerogâmica do Estado de São Paulo. Vol. 3. Ed. Hucitec, São Paulo. 386p. [TreeCo refID: 10145]

WANDERLEY, M.G.L.; SHEPHERD, G.J.; MELHEM, T.S. & GIULIETTI, A.M. (coords.) 2007. Flora Fanerogâmica do Estado de São Paulo. Vol. 5. Ed. Hucitec, São Paulo. 523p. [TreeCo refID: 10147]

WANDERLEY, M.G.L.; SHEPHERD, G.J.; MELHEM, T.S. & GIULIETTI, A.M. (coords.). 2005. Flora Fanerogâmica do Estado de São Paulo. Vol. 4. Ed. Hucitec, São Paulo. 437p. [TreeCo refID: 10146]

WANDERLEY, M.G.L.; SHEPHERD, G.J.; MELHEM, T.S.; GIULIETTI, A.M. & MARTINS, S.E. (coords.) 2009. Flora Fanerogâmica do Estado de São Paulo. Vol. 6. Ed. Hucitec, São Paulo. 330p. [TreeCo refID: 10148]

WANDERLEY, M.G.L.; SHEPHERD, G.J.; MELHEM, T.S.; GIULIETTI, A.M. & MARTINS, S.E. (coords.) 2012. Flora Fanerogâmica do Estado de São Paulo. Vol. 7. Ed. Hucitec, São Paulo. 393p. [TreeCo refID: 10149]

WEBSTER, G.L. 1982. Systematic status of the genus Kleinodendron (Euphorbiaceae). Taxon 31(3): 535-539. [TreeCo refID: 10431]

WIESBAUER, M.B.; GIEHL, E.L.H. & JARENKOW, J.A. 2008. Padrões morfológicos de diásporos de árvores e arvoretas zoocóricas no Parque Estadual de Itapuã, RS, Brasil. Acta Bot Brasilica 22(2):425-435. [TreeCo refID: 10113]

WITTMANN F.; SCHÖNGART J.; PAROLIN P.; WORBES M.; PIEDADE MTF. & JUNK W.J. 2006b. Wood specific gravity of trees in Amazonian white-water forests in relation to flooding. IAWA J, 27:255–266. apud WITTMANN, F.; ZORZI, B.T.; TIZIANEL, F.A.T.; URQUIZA, M.V.S.; FARIA, R.R.; SOUZA, N.M.; MÓDENA, É.S.; GAMARRA, R.M. & ROSA, A.L.M. 2008. Tree species composition, structure, and aboveground wood biomass of a riparian forest of the Lower Miranda River, southern Pantanal, Brazil. Folia Geobotânica 43(4): 397-411. [TreeCo refID: 10168]

WITTMANN, F.; SCHÖNGART, J.; DE BRITO, J.M.; WITTMANN, A.DEO.; PIEDADE, M.T.F.; PAROLIN, P.; JUNK, W.J. & GUILLAUMET, J.-L. 2010. Manual of trees in Central Amazonian várzea floodplains: taxonomy, ecology, and use. Editora INPA, Manaus. 286p. apud WITTMANN, F.; ZORZI, B.T.; TIZIANEL, F.A.T.; URQUIZA, M.V.S.; FARIA, R.R.; SOUZA, N.M.; MÓDENA, É.S.; GAMARRA, R.M. & ROSA, A.L.M. (2008). Tree species composition, structure, and aboveground wood biomass of a riparian forest of the Lower Miranda River, southern Pantanal, Brazil. Folia Geobotânica 43(4), 397-411. [TreeCo refID: 10169]

WITTMANN, F.; ZORZI, B.T.; TIZIANEL, F.A.T.; URQUIZA, M.V.S.; FARIA, R.R.; SOUSA, N.M.; MÓDENA, É.D.S.; GAMARRA, R.M. & ROSA, A.L.M. 2008. Tree species composition, structure, and aboveground wood biomass of a riparian forest of the lower Miranda River, Southern Pantanal, Brazil. Folia Geobotanica 43(4): 397-411. [TreeCo refID: 10118]

YUNCKER, T.G. 1972. The Piperaceae of Brazil I: Piper -Group I, II, III, IV. Hoehnea 2: 19-366. [TreeCo refID: 10342]

YUNCKER, T.G. 1973. The Piperaceae of Brazil II: Piper: Group V; Ottonia; Pothomorphe; Sarcorhachis. Hoehnea 3: 29-284. [TreeCo refID: 10343]

ZAMA, M.Y.; BOVOLENTA, Y.R.; CARVALHO, E.D.S.; RODRIGUES, D.R.; ARAUJO, C.G.D.; SORACE, M.A.D.F. & LUZ, D.G. 2012. Floristic composition and diaspore dispersal syndromes of shrubs and tree species in Parque Estadual Mata São Francisco, Paraná State, Brazil. Hoehnea 39(3): 369-378. [TreeCo refID: 10117]

ZAPPI, D. 2003. Revision of Rudgea (Rubiaceae) in Southeastern and Southern Brazil. Kew Bulletin 58(3): 513-596. [TreeCo refID: 10115]

ZECHINI, A.A.; SCHUSSLER, G.; SILVA, J.Z.; MATTOS, A.G.; PERONI, N.; MANTOVANI, A. & REIS, M.S. 2012. Produção, comercialização e identificação de variedades de pinhão no entorno da Floresta Nacional de Três Barras–SC. Biodiversidade Brasileira 2(2): 74-82. [TreeCo refID: 10404]

ZICKEL, C.S. 1989. Revisão taxonômica do gênero Lamanonia Vell. (Cunoniaceae). Dissertação (Mestrado). Universidade Estadual de Campinas, Campinas. 116p. [TreeCo refID: 10182]
